# Harnessing the Biosynthetic Diversity of Actinomycetes: Discovery of Unique Natural Products through Comparative Genomic and Metabolic Analysis

**DOI:** 10.1101/2024.12.06.627119

**Authors:** Yuta Kikuchi, Hayama Tsutsumi, Yoshihiro Watanabe, Hiroki Nakahara, Sho Ito, Yoshihiko Noguchi, Yuta Awano, Masanobu Kasuga, Masato Iwatsuki, Tomoyasu Hirose, Toshiaki Sunazuka, Yuki Inahashi

**Affiliations:** Ōmura Satoshi Memorial Institute, Kitasato University, 5-9-1 Shirokane, Minato-ku, Tokyo 108-8641, Japan; Graduate School of Infection Control Sciences, Kitasato University, 5-9-1 Shirokane, Minato-ku, Tokyo 108-8641, Japan; DIC Central Research Laboratories, 631, Sakado, Sakura, Chiba 285-8668

**Keywords:** natural products, genome analysis, actinomycetes molecular networking, biosynthesis

## Abstract

Actinomycetes, rich in biosynthetic gene clusters (BGCs), are important sources of natural products (NPs). However, the rate of discovering novel NPs from actinomycetes has declined, indicating the need for a new strategy to obtain NPs. Herein, we present a strategy for the efficient discovery of novel NPs. First, we performed a comprehensive analysis of BGCs in actinomycetes, evaluating the average number and types of BGCs per genus to identify prolific NP producers. Our analysis revealed that certain actinomycetes strains, such as those in the family *Pseudonocardiaceae*, possess a greater number of BGCs than *Streptomyces*. In addition, these strains tend to possess strain-specific BGCs compared with others. To facilitate the identification of strain-specific compounds, we developed a comparative metabolic analysis method using molecular networking. Applying this method to eight *Pseudonocardiaceae* strains, we successfully discovered two novel peptides, lentindoles A (**1**) and B (**2**), featuring a 6/5/6 and 6/5/5 tricyclic ring system, respectively, along with their possible biosynthetic precursor, lentindole C (**3**). Our result demonstrates that the effectiveness of our developed strategy, which is expected to accelerate the discovery of novel NPs.

## Introduction

Natural products (NPs) isolated from microorganisms have played a crucial role in drug development for decades.^[1]^ To address challenges such as drug-resistant pathogens, emerging infections, and neglected diseases, the discovery of novel NPs from microorganisms is essential.^[2,3]^ However, recently, the rate of discovering new NPs has decreased, while the rediscovery of known NPs has become a common problem.^[4,5]^ This may be attributed to the focus of such research mainly on *Streptomyces*, which have been the primary sources of NPs, and the reliance on activity-based screening methods.^[6]^ Therefore, alternative sources of NPs and innovative methods for discovering novel NPs are urgently needed.

Rare actinomycetes, excluding *Streptomyces*, represent a promising alternative source of NPs because recent genomic analyses have revealed that their biosynthetic potential is comparable to that of *Streptomyces*.^[7]^ In our previous work, guided by physicochemical properties such as ultraviolet (UV) spectra and molecular weight, we discovered several overlooked NPs from rare actinomycetes, such as actinoallolides that exhibited antitrypanosomal activity.^[8]^ These findings suggested that untapped microorganisms with prolific NP-producing capabilities could serve as valuable sources of novel NPs. To accelerate the discovery of NPs, selecting rare actinomycetes genera with prolific NP-producing potential is essential. Genomic analysis for assessing the number of secondary metabolite biosynthetic gene clusters (sBGCs) has also been shown to be effective in identifying promising NP producers.^[9–11]^ However, accurately evaluating biosynthetic potential requires more detailed analyses, such as examining the diversity of BGCs within genera and the average number of BGCs per genus.

Molecular networking (MN) is a recently developed technique for metabolomic analysis using MS/MS.^[12–14]^ This technique allows the connection and visualization of structurally similar compounds based on MS/MS spectra similarity, enabling the detection of many “dark matter” metabolites. Furthermore, recently, Global Natural Products Social Molecular Networking (GNPS) was demonstrated to facilitate the identification of both primary and secondary known metabolites, and aid in the analysis of these “dark matters”. The MN-based approach has recently expanded the strategies available for novel NPs discovery, resulting in the identification of many new compounds.^[15]^

In this study, we combined comprehensive genomic and comparative metabolomic analyses for the rapid discovery of novel NPs. We first analyzed the numbers and diversity of sBGCs among *Actinomycetota* to search for prolific NP producers. Using this analysis, we focused on rare actinomycetes, specifically the family *Pseudonocardiaceae*, and employed MN-based method created in this study to detect NPs produced exclusively by individual strains (**Figure 1**). Consequently, we discovered lentindoles A (**1**) and B (**2**), two novel NPs with a 6/5/6 or 6/5/5 fused ring systems, along with their possible biosynthetic precursor lentindole C (**3**) (**Figure 1 and 5A)**.

**Figure 1.**
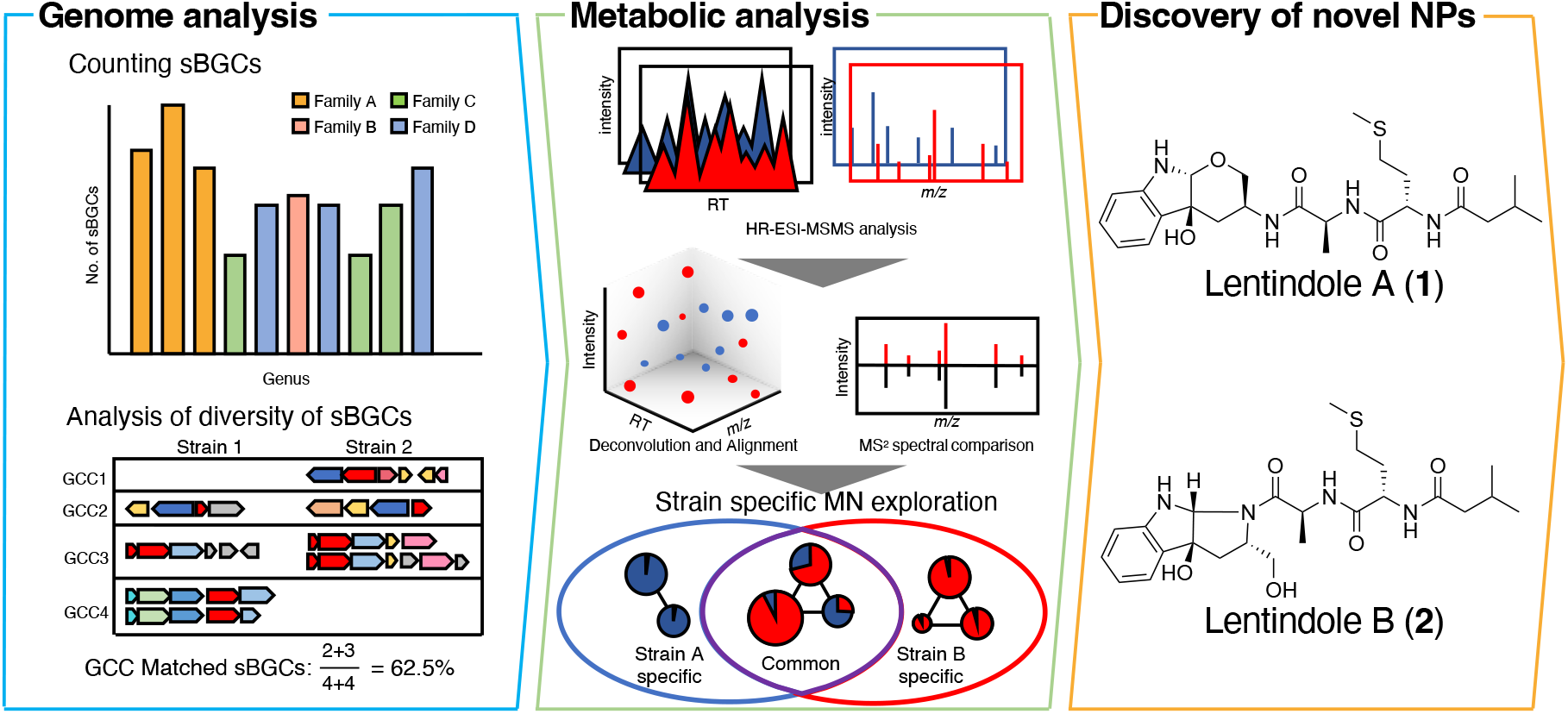
Overview of our strategy for identifying prolific producers of natural products and discovering novel natural products, along with the novel natural products obtained in this study.

## Results and Discussion

### Comprehensive analysis of biosynthetic gene clusters in *Actinomycetota*

To assess the potential for NP production in each genus of *Actinomycetota*, we initially estimated the average number of sBGCs (**Figure 1**). A total of 6781 *Actinomycetota* genome data deposited in the National Center for Biotechnology Information Reference Sequence Database^[16]^ was analyzed using antiSMASH6.0^[17]^ to detect the sBGCs of each *Actinomycetota* strain. Subsequently, we counted the number of sBGCs related to the biosynthesis of polyketide (PK), nonribosomal peptide (NRP), terpene (TP), and ribosomally synthesized and posttranslationally modified peptides (RiPPs) (PNTR-BGC) in each strain (Scheme S1, detailed sBGC type used to count the number of sBGCs described in Table S2). Next, we calculated the number of sBGCs in each genus by averaging the counted number of sBGCs. This analysis generated an average number of PNTR-BGCs in each of the 325 genera (Figure 2, Figure S1, Table S1).

**Figure 2.**
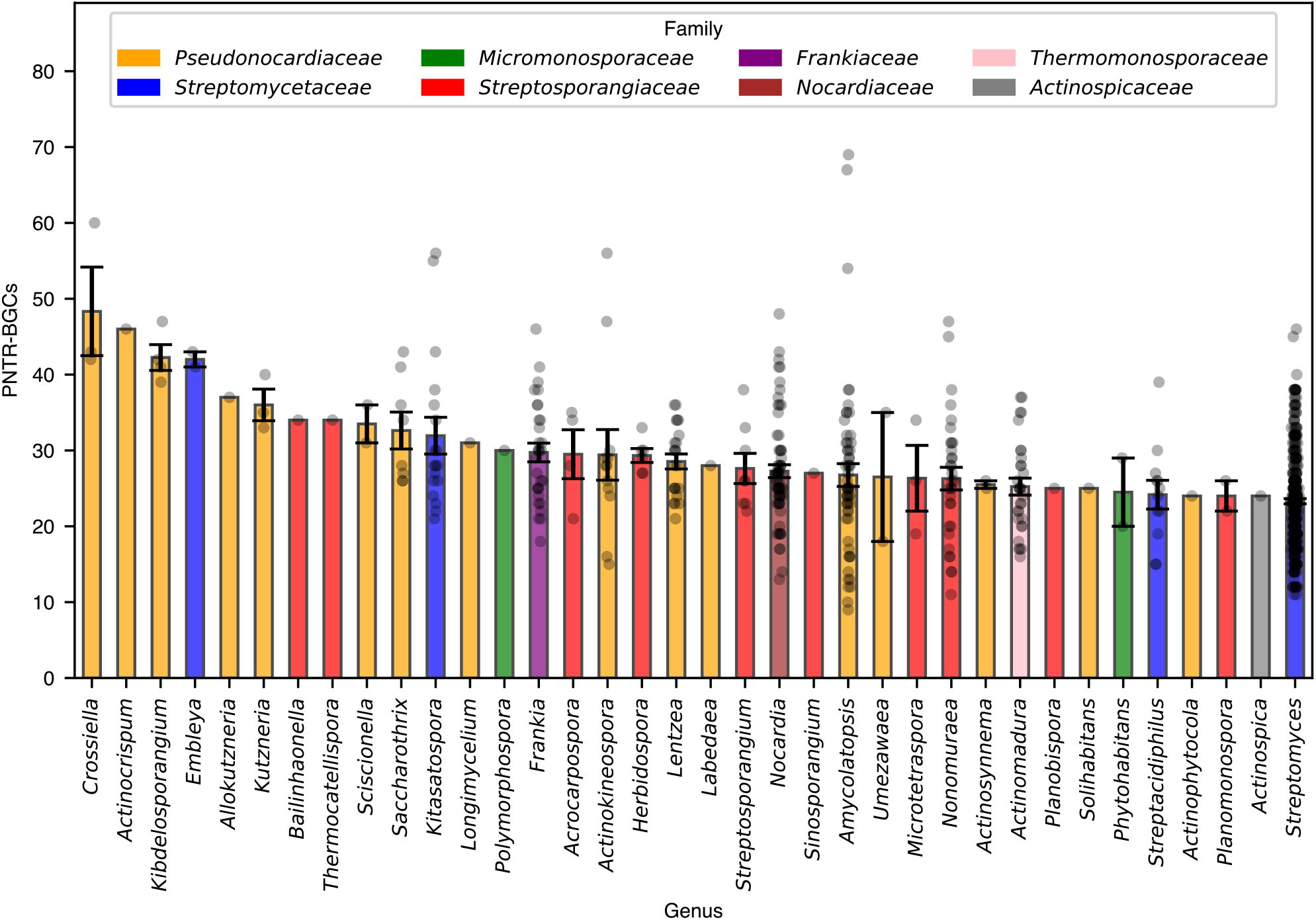
Comparison of the average number of PNTR-BGCs among genera within actinomycetes. In this figure, genera with a higher average number of PNTR-BGCs than *Streptomyces* are shown. Genera belonging to *Pseudonocardiaceae, Micromonosporaceae, Frankiaceae, Thermomonosporaceae, Streptomycetaceae, Streptosporangiaceae, Nocardiaceae*, and *Actinospicaceae* are colored yellow, green, purple, pink, blue, red, orange and brown, respectively.

We identified 35 genera harboring a number of PNTR-BGCs equal to or greater than that in *Streptomyces* (23.3 PNTR-BGCs), which is known as a prolific NP producer (**Figure 2, Table S1**). Classification of these genera at the family level revealed that they belonged to *Actinospicaceae* (1 genus), *Frankiaceae* (1 genus), *Micromonosporaceae* (2 genera), *Nocardiaceae* (1 genus), *Pseudonocardiaceae* (16 genera), *Streptomycetaceae* (3 genera), *Streptosporangiaceae* (10 genera), and *Thermomonosporaceae* (1 genus) (**Figure 2, Table S1**). Because one sBGC is involved in the biosynthesis of one secondary metabolite (and its associates such as derivative and intermediates), we hypothesized that these 35 genera with a high average number of sBGC may biosynthesize a wide variety of secondary metabolites. Next, we investigated the diversities of PNTR-BGC in the genus *Streptomyces* and some families of rare actinomycetes (*Streptomycetaceae* excluding *Streptomyces, Pseudonocardiaceae, Micromonosporaceae, Nocardiaceae*, and *Streptosporangiaceae*) by comparing types of gene cluster clans (GCCs) between strains using BiG-SCAPE,^[18]^ because these selected families contain some genera harboring a high number of PNTR-sBGCs, with some of them having more sBGCs than *Streptomyces* (**Figure 1, Scheme S1**). To estimate the taxonomic relationships between strains and the differences in PNTR-BGCs, we calculated the similarities in the 16S rRNA gene and GCCs between strains (**Figure 3**). In *Streptomyces*, the concordance rates of GCCs between the two strains ranged from 10% to 100% (**Figure 3A**). Notably, two strains, which exhibited a low 16S rRNA gene similarity (< 98.7%) and were identified as distinct species,^[19]^ had different types of GCCs, with the concordance rate ranging from 10% to 40% (**Figure 3A**). In most cases, the concordance rates of GCCs between different species were within the range of 10–20% (**Figure 3A**). These findings revealed that the species diversities are correlated with the diversities of GCCs. We observed similar trends in the families *Micromonosporaceae, Pseudonocardiaceae, Streptomycetaceae* (excluding *Streptomyces*), and *Streptosporangiaceae*; however, the concordance rates of GCCs between different strains of *Streptomycetaceae* were slightly higher (20–40%) than those in other families (**Figure 3B–E**). In contrast, the concordance rates of GCCs between species in family *Nocardiaceae* were high (60– 90%), with strains possessing similar PNTR-BGCs. This indicated that strains belonging to the family *Nocardiaceae* may produce structurally related NPs (**Figure 3E**). These results clearly revealed that *Micromonosporaceae, Pseudonocardiaceae, Streptomycetaceae*, and *Streptosporangiaceae* have specific sBGCs at species level and the ability to produce diverse secondary metabolites. This also suggested that exploring NPs only produced by one species belonging to any of these four families could lead to the discovery of novel NPs.

**Figure 3.**
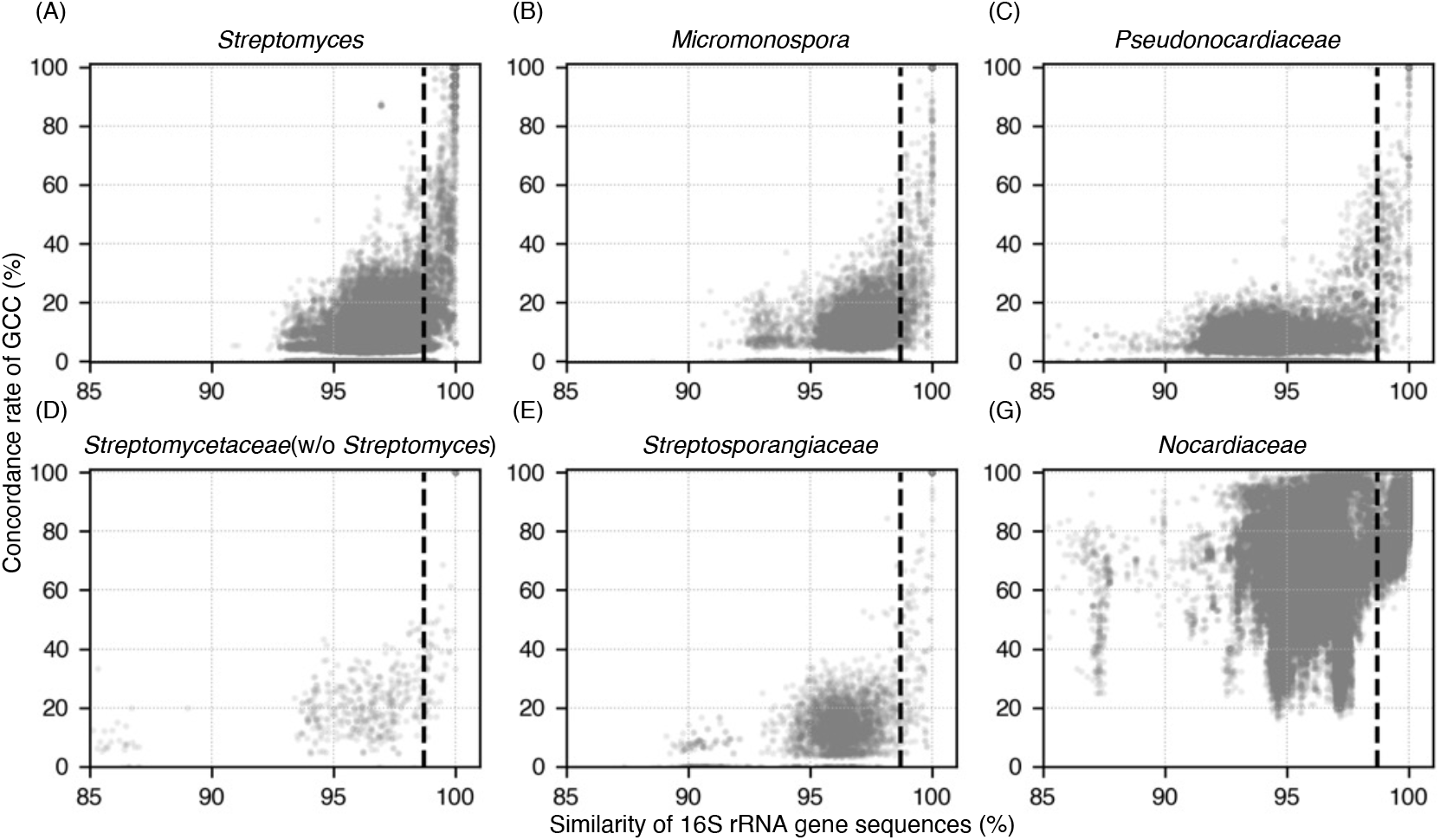
Scatter plot of the concordance correlation of GCCs and similarity of 16S rDNA sequences among *Streptomyces, Micromonosporaceae, Pseudonocardiaceae, Streptomycetaceae* (excluding *Streptomyces*), *Streptosporangiaceae* and *Nocardiaceae*. Each node represents the result of a comparison between two strains. The dotted lines indicate 98.7% sequence similarity, as sequence similarities below this threshold typically indicate different species.

### Exploration of novel metabolites using molecular networking analysis

Our comprehensive sBGC analysis of *Actinomycetota* suggested that some genera possess many PNTR-BGCs, with each species possessing specific PNTR-BGCs. Notably, mining strain-specific NPs from various strains that possess many sBGCs, such as strains belonging to *Pseudonocardiaceae*, would lead to the discovery of novel NPs. This prompted us to obtain strain-specific compounds from culture broths. To this end, we constructed a Python script, which is a molecular network-based method for easily identifying strain-specific compounds (https://github.com/microbialfunc/MNAnalyzer). In this method, (i) culture broths of strains belonging to the same family, prepared using the one strain many compounds (OSMAC) approach,^[20]^ were analyzed by LC-HRMS, (ii) obtained data were deconvoluted using MS-DIAL^[21]^ (see Materials and methods) (iii) deconvoluted data were analyzed by feature-based molecular networking (FBMN)^[22]^ in Global Natural Products Social Molecular Networking (GNPS)^[23]^, (iv) the production of each compound was compared based on the intensity, and (v) data were sharpened to visualize strain-specific compounds using Cytoscape^[24]^ (**Figure 1, Scheme S2**). In this study, we prepared 160 broths by culturing eight strains in 20 different medium conditions (listed in **Tables S3** and **S4)** and applied our method to search for strain-specific NPs in the prepared broths. As a result, we detected a total of 10494 compounds, 2102 of which were identified as components of the culture media, whereas the remaining 8392 compounds were identified as metabolites (**Figure S2**). Subsequently, among the 8392 metabolites, 41 were identified as known compounds based on the MS/MS spectra database of GNPS, whereas the rest (8351 metabolites) were unidentified metabolites. Among the 8351 unidentified metabolites, 1256 formed molecular networks (MNs) with one or more other nodes, whereas 7095 unknown metabolites were singleton (**Figure S2**). In addition, among the unidentified compounds forming MNs, 496 formed MNs with medium components, 45 formed MNs with known metabolites, 130 formed MNs with both known metabolites and medium components, and 585 formed MNs only with other unknown metabolites (Figure S2). Subsequently, to identify strain-specific unknown metabolites, we investigated whether all compounds in each MN and compounds forming singleton nodes were produced by a sole strain based on ion intensity. As a result, we detected 114 MNs (Figure S2). We focused on the MN containing four compounds produced by *Lentzea* sp. OK19-0192, in which two of the four compounds showed an *m/z* of 493.25, whereas the others showed *m/z* values of 477.25 and 409.19, respectively, because other strains did not produce all these compounds (Figure 4) and no similar compounds with the same accurate mass were found in the Dictionary of Natural Products^[25]^ and Natural Products Atlas.^[26,27]^ The compound with the *m/z* of 409.19 formed the same MN as that of the compound with the *m/z* of 493.25 despite a mass difference of *m/z* 84.06, suggesting that it shares a similar structure with the latter but lacks the isovaleric acid moiety. In addition, it had the same retention time as that of the compound with the *m/z* of 493.25 (Figure 4B). These observations suggested that the compound with the *m/z* of 409.19 is likely to be an artifact generated during MN analysis.

**Figure 4.**
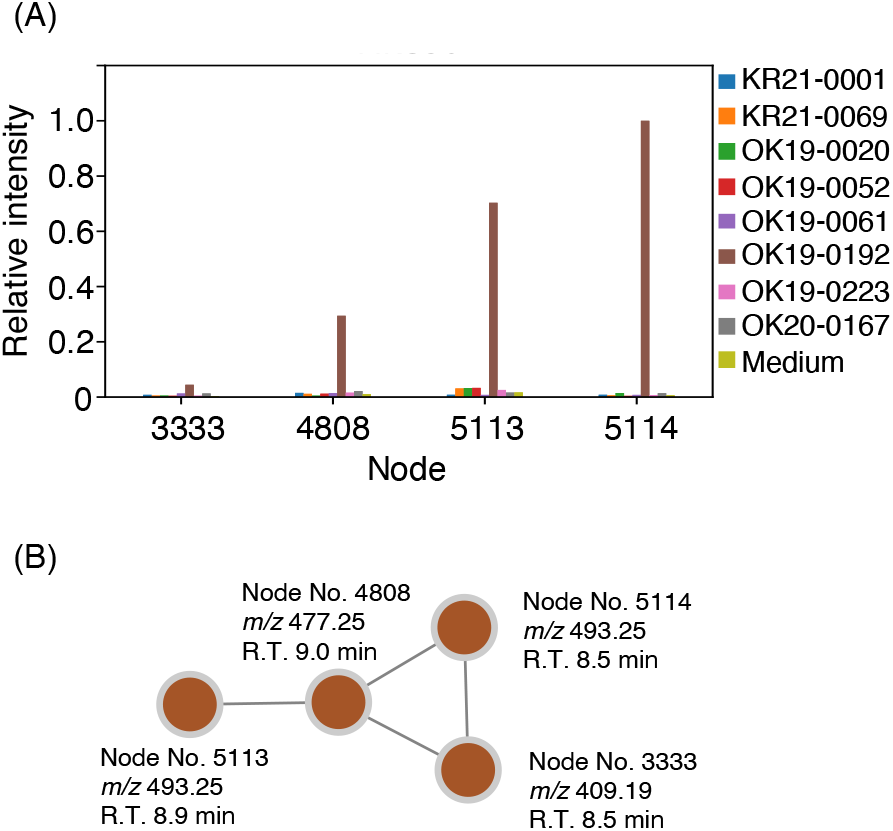
(A) Comparison of the productivity of each target compound across strains based on ion intensity. (B) The molecular network consists of target compounds specifically produced by one strain among the tested strains.

### Isolation and structure elucidation of lentindoles A (1), B (2) and C (3)

The procedure for the isolation and purification of lentindoles A (**1**), B (**2**), and C (**3**) **(Figure 5A**) is summarized in **Scheme S3**. The physicochemical properties of **1** and **2** are shown in **Table S5**. Compounds **1, 2**, and **3** exhibited an *m/z* of 493.2468 [M+H]^+^ (calcd. *m/z* of 493.2479; C_24_H_37_N_4_O_5_S), 493.2484 [M+H]^+^ (calcd. *m/z* of 493.2479; C_24_H_37_N_4_O_5_S), and 477.2530 [M+H]^+^ (calcd. *m/z* of 477.2479; C_24_H_37_N_4_O_4_S), respectively.

**Figure 5.**
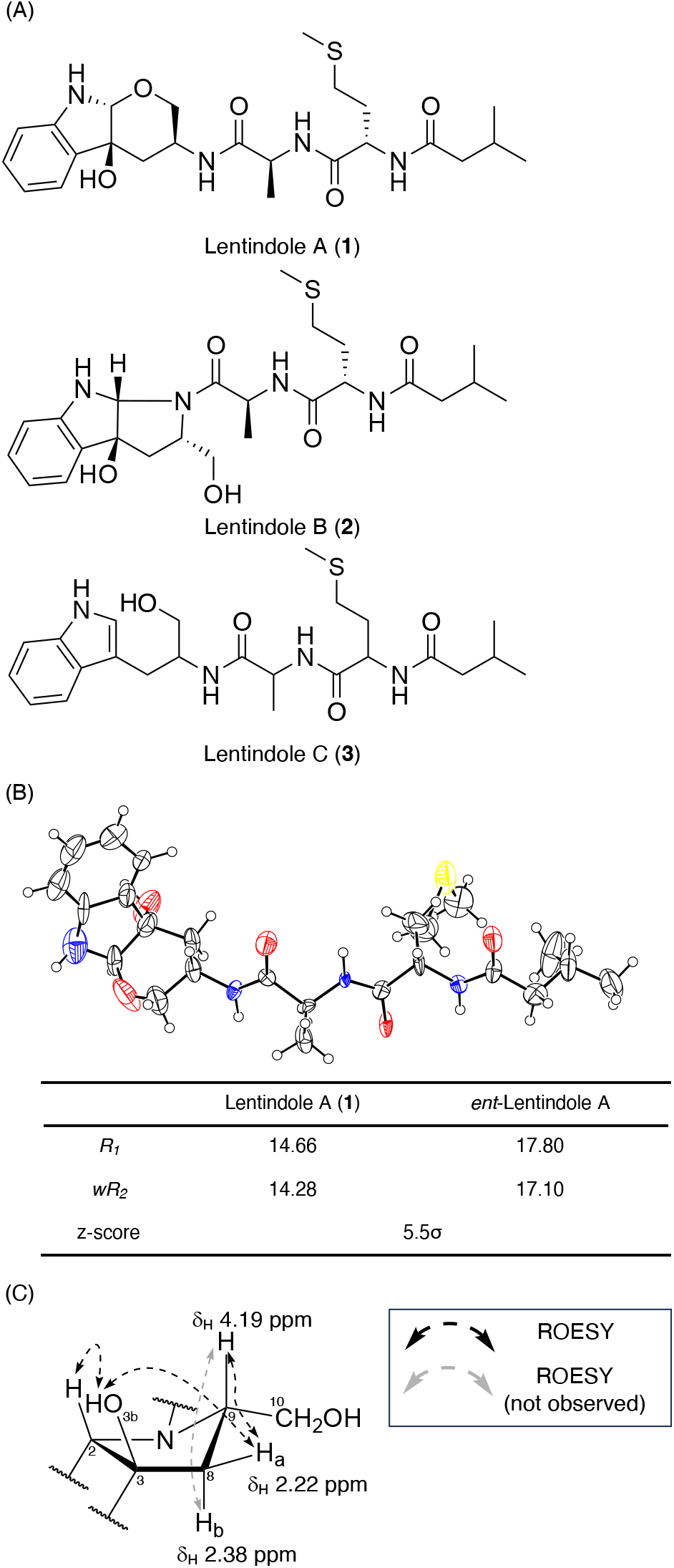
Structure of lentindoles A (**1**), B (**2**) and C (**3**) and stereochemical analysis of **1** and **2** using NMR and 3D ED/MicroED, respectively. (A) Structure of lentindoles A–C (**1–3**). (B) ORTEP obtained from crystal structure of **1**, and statistics calculated from the dynamical refinement result. (C) Key ROESY correlations of 2-pyrrolidin-2-ylmethanol moiety of **2**.

The elucidation of the planner structures of **1** and **2** is described in the **Supporting Information**. Detailed ^1^H NMR, ^13^C NMR, COSY, HMQC and HMBC analyses of **1** revealed that it has 3-amino-3,4,9,9a-tetrahydropyrano[2,3-*b*]indol-4a(2*H*)-ol (THPyra), alanine (Ala), methionine (Met), and isovaleric acid (Iva) moieties in the sequence of Iva-Met-Ala-THPyra (**Figure 5A, Figure S3A and S9–13**).

The ^1^H NMR of **2** indicated it to be a mixture of two compounds. However, these were equilibrating species, as the ratio of proton signals changed from 1:1 (measured in DMSO-*d*_6_) to 1:2.5 (measured in CDCl_3_) (**Figure S4A**), with the proton signals corresponding to the same position in the planar structure getting closer at 60°C (**Figure S4B**). These findings suggested that the two structures observed in NMR analysis formed a rotamer,^[28]^ and **2** was not a mixture of stereoisomers. The ^1^H NMR of **2** was similar to that of **1**, except for the presence of an additional hydroxy proton and the absence of the amide proton. Detailed 2D NMR analyses, including ROESY, revealed that **2** has a structure similar to **1** with a 2-(hydroxymethyl)-2,3,8,8a-tetrahydropyrrolo[2,3-*b*]indol-3a(1*H*)-ol (THPyro) moiety instead of a THPyra moiety (**Figure S3A**).

The absolute configuration of alanine and methionine moieties in **1** and **2** were determined using an advanced Marfey’s method.^[29,30]^ After the hydrolysis of **1** and **2**, the obtained amino acids were derivatized with D-FDLA, and the reaction products were analyzed by LC-HRMS. The comparison of authentic samples revealed that all amino acids in **1** and **2** had the L configuration (**Figure S5**). Furthermore, the stereochemistry of the tricyclic structure of **1** was determined using 3D electron diffraction/micro electron diffraction (3D ED/MicroED) (**Figure S6, Table S8**, CCDC 2406780). This analysis revealed that the relative stereochemistry of **1** should be 2*R*\*, 3*S*\*, 9*S*\*, 13*S*\*, and 17*S*\* (**Figure S7**). Considering **1** has L-alanine and L-methionine, the absolute configuration of **1** was determined as 2*R*, 3*S*, 9*S*, 13*S*, and 17*S* (**Figure 5A**). Moreover, the comparison of *R1* and *wR2* values calculated using **1** and its enantiomer (*ent*-**1**) without hydrogen atoms revealed that the values for **1** were lower than those for *ent*-**1** (**Figure 5B**). This suggested that the absolute configuration of **1** can be determined solely using 3D ED/MicroED.^[31,32]^

The absolute configuration of **1** and biosynthetic analysis (described later) indicated that the THPyra and THPyro moieties are likely synthesized from L-tryptophan. The ROESY spectra of **2** indicated that H-2, OH-3, and H-9 should be in the same orientation (**Figure 5C, Figure S3B and S19**). Therefore, the absolute configuration of **2** was determined to be 2*R*, 3*S*, 9*S*, 13*S*, and 17*S*.

The molecular formula of **3** (C_24_H_36_N_4_O_4_S) and MS/MS analysis indicated that **3** is a peptide consisting of L-tryptophanol, L-alanine, L-methionine, and isovaleric acid; however, the poor yield and low purity of compound **3** were insufficient for structure elucidation using NMR. Next, we synthesized **3** via liquid-phase peptide synthesis (LPPS) using soluble hydrophobic tag auxiliary described in **Supporting Information**. Because the retention time, *m/z*, and MS/MS spectra of compound **3** matched well with those of synthetic **3**, we concluded that **3** was *N*-(1-((1-hydroxy-3-(1*H*-indol-3-yl)propan-2-yl)amino)-1-oxopropan-2-yl)-2-(3-methylbutanamido)-4-(methylthio)butanamide (**Figure S7**).

### Biosynthetic implication of lentindoles

The THPyra moiety was reported to be formed from *N*-aminoacyl-L-tryptophanol such as L-valyl-L-tryptophanol by cytochrome P450s, TleB, and HinD^[33]^ via epoxidation and subsequent cyclization. Thus, we hypothesized that **1** was formed in the same manner using **3** as a substrate, which is presumably synthesized within an organism, as a substrate. To investigate whether the precursor was synthesized *de novo* in OK19-0192, we conducted isotope-labeled or fluorine-labeled compound feeding experiments. In the presence of 6-fluoro-DL-tryptophan, ^13^C-L-methionine, or L-alanine-*d*_4_, ion peaks with *m*/*z* of 511.2389 [M+H]^+^ (C_24_H_36_N_4_O_5_SF, calc. *m/z* of 511.2390 [M+H]^+^), *m*/*z* of 494.2520 [M+H]^+^ (C ^13^C_1_H_37_N_4_O_5_S, calc. *m/z* of 494.2519 [M+H]^+^) and *m*/*z* of 497.2719 [M+H^+^] (C_24_H_33_D_4_N_4_O_5_S, calc. *m/z* of 497.2736 [M+H]^+^) were observed (lentindole A and B; C_24_H_37_N_4_O_5_S, calc. *m*/*z* of 493.2464 [M+H]^+^), respectively (**Figure S8**). These results revealed that lentindoles are biosynthesized via nonribosomal or ribosomal peptide formation from L-tryptophane, L-alanine, and L-methionine, followed by successive modifications such as epoxidation and cyclization. Next, we analyzed the genome sequence of the strain OK19-0192 and searched for the BGC of lentindole. As a result, we discovered one potential BGC containing genes encoding polyketide synthase (PKS) with a reductase domain^[34]^ for synthesizing the tryptophanol moiety, nonribosomal peptide synthetase (NRPS) responsible for peptide bond formation, and flavin-dependent oxygenase, which presumably catalyzes oxidation of indole moiety (DDBJ accession number: LC852381, **Table S9**). Based on the detailed bioinformatic analysis of this BGC and structure of **1** and **2**, we proposed a predicted biosynthetic pathway, as described in **Scheme S4**. Initially, L-methionine is loaded onto an NRPS, LtdA and *N*-acylated by β-ketoacyl-(acyl-carrier-protein) synthase III LtdH using isovaleric-CoA. Then, *N*-isovaleryl-L-Met, L-Ala, and L-Trp are condensed by NRPSs LtdB and LtdI to form a *N*-isovaleryl tripeptide, which is tethered to the carrier protein (CP) of LtdI. This tripeptide is transferred onto the CP of LtdD and released as **3** through a four-electron reduction catalyzed by the terminal reductase domain of LtdD. The indole moiety in 3 is then oxidized by the luciferase-like monooxygenase LtdL to form 3-hydroxyindolenine, followed by ring closure by nucleophilic attack from either the hydroxy group of the tryptophanol moiety or the amide nitrogen between THPyro and L-Ala to the C2 carbon of indole to produce **1** or **2** in the same manner as for chemoenzymatically synthesized teleocidin analogs^[33]^ and notoamide D^[35]^, respectively. For a more detailed biosynthetic investigation, further studies are needed, including metabolic analysis of gene deletion mutants and biochemical characterization of biosynthetic enzymes. These will be the focus of our next research effort.

## Conclusion

We conducted comparative genomic and metabolic analyses to efficiently select prolific NP producers and search for novel NPs. Our genome analysis using 6781 *Actinomycetota* genome data showed that some rare actinomycetes, such as those belonging to *Pseudonocardiaceae*, have abundant and diverse sBGCs and are prolific NP producers. In this study, 29 genera of *Pseudonocardiaceae* were analyzed, and 16 of them were shown to have more PNTR-BGCs than *Streptomyces*. Of the remaining 13 genera, seven genera were found to have more than 10 PNTR-BGCs, whereas the other six had fewer than 10 PNTR-BGCs. Approximately 80% of genera in the family *Pseudonocardiaceae* possess a significant number of PNTR-BGCs, indicating that they are likely prolific producers of NPs. In addition, each prolific producer was also revealed to produce specialized and unique NPs. Thus, we constructed a method to easily and efficiently identify NPs specifically produced by a single strain by combining the OSMAC approach with MN analysis. This method enabled us to easily distinguish between the compounds that are medium components, metabolites converted from medium components, and common bacterial metabolites based on MS/MS spectrum and ion intensities of precursor ion. We applied this method to compare the metabolites from eight strains belonging to *Pseudonocardiaceae*. As a result, lentindoles A and B which are the first NPs with unique tetrahydropyranoindole (6/5/6 fused ring) and tetrahydropyrroloindole (6/5/5 fused ring) structure, respectively, were discovered from *Lentzea* sp. OK19-0192. While many NPs with indole-derived tricyclic systems have been reported, none contain a THPyra moiety, underscoring the effectiveness of our strategy. Our *in silico* biosynthetic analysis of lentindoles led to the discovery of a possible BGC. Notably, a similar BGC of lentindoles was only identified in *Lentzea aerocolonigenes* NBRC 13195^T^. This finding also highlights the potential of our methods to facilitate the discovery of novel secondary metabolites synthesized by unique BGCs. Further application of this strategy to explore microbial-produced NPs is expected to lead to the discovery of novel drug candidates.

## Supporting Information

The authors provide experimental procedures in the Supporting Information and have cited additional references therein^[36–45]^.

## Acknowledgements

We are grateful to Distinguished Emeritus Professor Satoshi ōmura (Kitasato University) for his helpful support and valuable guidance and suggestions. We are also grateful to Mr. Seizo Hino for his kind support. We thank Dr. Kenichiro Nagai, and Ms. Noriko Sato (School of Pharmacy, Kitasato University) for various instrumental analyses, Mr Toshihiro Suzuki, Mr Sho Hirata, Dr Eiji Okunishi (JEOL, Ltd), and Dr Akihito Yamano (Rigaku, Co.) for their support in 3D ED/Micro ED operation, and Dr. Paul B. Klar (Czech Academy of Sciences) for his support in analysis of crystal data obtained by 3D ED/Micro ED. This research was supported by Platform Project for Supporting Drug Discovery and Life Science Research (Basis for Supporting Innovative Drug Discovery and Life Science Research (BINDS)) from AMED under Grant Number JP22ama121035, Okinawan Create Leading Projects in Growing Fields 2017-2019 and 2020-2021, OKINAWA Prefectural Government and by JSPS KAKENHI Grant Numbers JP19H05685, JP22K15305 and JP23H04561.

## Entry for the Table of Contents

Comparative analysis of secondary metabolite biosynthetic gene clusters (sBGCs) in Actinomycetes revealed that *Pseudonocardiaceae* strains possess abundant, diverse and strain-specific sBGCs. Molecular networking identified lentindoles A and B, novel peptides with rare 6/5/6 and 6/5/5 tricyclic scaffolds, as strain-specific metabolites from a *Lentzea* strain. This approach offers a promising strategy for discovering novel natural products.

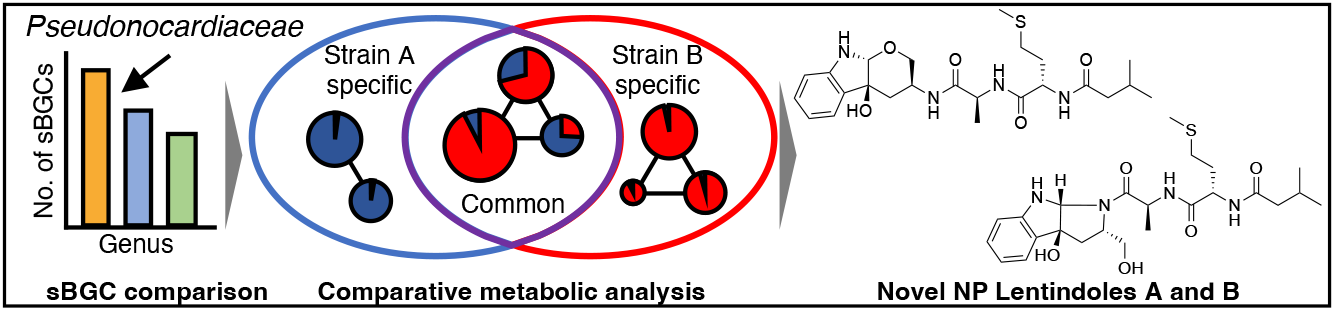

