## Supporting information for "Harnessing the Biosynthetic Diversity of Actinomycetes: Discovery of Unique Natural Products through Comparative Genomic and Metabolic Analysis"

Table of contents

Experimental Section ······ 3

1. General experiment ······ 3
2. Estimation of secondary metabolic ability in *Actinomycetota* ······ 4
2.1. Detection of sBGCs by antiSMASH and counting the number of sBGCs per genus ······ 4
2.2. Gene cluster clan analysis among the same family ······ 4
3. Exploration of novel secondary metabolites from *Pseudonocardiaceae* ······ 5
3.4. Synthesis procedures of lentindole C (3) and characterization data ······ 9
4. Feeding of labeled amino acids to analyze the biosynthesis pathway of lentindole ······ 15
5. Genome analysis ······ 15
6. Phylogenetic analysis ······ 15

### Experimental Section

#### 1. General experiment

Unless otherwise noted, commercial reagents were purchased from Sigma Aldrich, Combi-blocks, TCI, Strem Chemicals, FUJIFILM Wako Pure Chemical Co., Watanabe Chemical Industries LTD., BLDPharm and/or Kanto Chemical Co., and used without additional purification. Solvents were purchased from Sigma Aldrich, TCI and/or Kanto Chemical Co, and used without additional purification (stored over molecular sieves). THF and DCM were sparged with argon and dried over molecular sieves prior to use. 3,4,5-Tris(octadecyloxy)benzoic acid methyl ester (**4**) was prepared by following the literature procedure[1].

High-performance liquid chromatography-electron splay ionization-mass spectrometry analyses were carried out using a Triple TOF 5600+ System (AB Sciex, Framingham, MA, USA) equipped with an ExionLC AC system (AB Sciex, Framingham, MA, USA) with Capcell Core C18 column (3.0 i.d. × 100 mm; Osaka Soda Co., Ltd., Osaka, Japan). The mobile phase consisted of water containing formic acid (solvent A) and methanol containing formic acid (solvent B). The samples were subjected to a linear gradient of solvent A/solvent B (0–13 min 5–100% solvent B, 13-16 min 100% solvent B) as a mobile phase at 0.6 mL/min. NMR spectra were measured using an Agilent 400-MR DD2 (Agilent Technologies) with $^{1}$H NMR and $^{13}$C NMR at 500 and 125 MHz, respectively. Preparative liquid chromatography was performed using an HPLC Nexera system with LC-20AR as pumps, SPD-M40 as a photodiode array detector (Shimadzu Co., Ltd., Kyoto, Japan), equipped with Pegasil ODS SP100 column (20 i.d. × 250 mm; Senshu Scientific Co. Ltd., Tokyo, Japan). 3D ED/MicroED data were collected with an XtaLAB Synergy-ED (Rigaku Co., Tokyo, Japan, and JEOL Ltd). SCXRD data were collected using an XtaLAB Synergy-S (Rigaku Co.). Infrared attenuated total reflectance spectra (IR-ATR) were measured using an FT-4600 Fourier transform infrared spectrometer (JASCO Corporation, Tokyo, Japan) with diamond crystal prism. The UV spectra were measured using SpectraMax QuickDrop™ (Molecular Devices, LLC., San Jose, USA) spectrometers with

MeOH as the solvent. Optical rotation was measured with a JASCO P-2200 polarimeter (JASCO Co., Tokyo, Japan).

#### 2. Estimation of secondary metabolic ability in *Actinomycetota*

##### 2.1. Detection of sBGCs by antiSMASH and counting the number of sBGCs per genus

We analyzed 6781 sequenced *Actinomycetota* genomes (scaffold: 4148, complete: 2633, 325 genera) downloaded from the National Center for Biotechnology Information (NCBI) Reference Sequence (RefSeq) Database[2] using antiSMASH 6.0.0,[3] and counted the secondary metabolite biosynthetic gene clusters (sBGCs) related to the biosynthesis of polyketides (PKs), nonribosomal peptides (NRPs), terpenes (TPs), and ribosomally synthesized and post-translationally modified peptides (RiPPs) based on some types of BGCs detected using antiSMASH (types used for counting are listed in **Table S1**). These counted sBGCs were termed PNTR-BGCs. This analysis was carried out under the default condition except for Prodigal,[4] which was used for the gene annotation. Even if one BGC region contained some BGCs as indicated by antiSMASH analysis, that BGC region was counted as a single BGC. The PNTR-BGC counts in each genus were averaged.

##### 2.2. Gene cluster clan analysis among the same family

All PNTR-BGCs data (Genbank format files) generated using antiSMASH were analyzed using BiG-SCAPE with default settings, except for the glocal mode that was used for analysis. From the output result files, we obtained GCC information for each genus. The GCC concordance rates between two strains were calculated as follows. The total number of PNTR-BGCs was calculated by adding the number of PNTR-BGCs from each of two strains. The number of GCC-matched PNTR-BGCs was determined by excluding the number of GCC-unmatched PNTR-BGCs between the two strains from the total number of PNTR-BGCs. Dividing the number of GCC-matched PNTR-BGCs by the total number of BGCs gave the GCC concordance rate. For example, if strain A has three PNTR-BGCs classified as GCC1, GCC2, and GCC3, and strain B has four PNTR-BGCs classified as GCC1, GCC2, GCC4 and GCC5, the total number of PNTR-BGCs and number of GCC-matched PNTR-BGCs are 7 and 4 respectively, resulting in a concordance rate of 0.57 (= 4/7) (57%).

The 16S rRNA gene sequences of each genus were extracted from the gene annotation files used for antiSMASH analysis. The similarity between two strains was calculated using the BLAST algorithm. The correlation between the GCC concordance rate and similarities in the 16S rRNA sequence between two strains was plotted using Python matplotlib.

#### 3. Exploration of novel secondary metabolites from *Pseudonocardiaceae*

##### 3.1. Molecular networking analysis procedure

###### 3.1.1. Cultivation of strains

A 10 mL of seed medium (**Table S4**) in a 70-mL test tube was inoculated with one inoculating loop of each strain or 100 $\mu$L of the glycerol stock of each strain (the eight strains used in this study are listed in **Table S3**) and shaken at 300 rpm for 5 d at 27 °C. We applied the OSMAC (One Strain-Many Compound) approach on these eight strains using 20 different production media (**Table S4**). A 10 mL of each production medium in a 70-mL test tube was inoculated with 100 $\mu$L of the seed culture and shaken at 300 rpm for 6 d at 27 °C. An equal volume of ethanol was added to each culture broth (160 broths in total) and the 20 media used as controls, and the mixtures were shaken at 300 rpm for 2 h. Subsequently, the mixtures were centrifuged at 12 000 rpm and the supernatants were used for LC-HRMS analysis.

###### 3.1.2. Molecular Networking analysis to detect strain-specific natural products

A total of 181 samples (160 broth, 20 media, and one blank control) were analyzed using LC-HRMS. The analytical data were converted into abf format files using the *Reifycs* Analysis Base File Converter (URL: https://www.reifycs.com/abfconverter/), while the obtained spectra were deconvoluted and aligned using MS-DIAL[5] with optimized settings (**Table S10**). In our analysis with MS-DIAL, medium components were also detected using our in-house medium components library, which was constructed from LC-HRMS data from the 20 medium and one blank (MeOH) sample. The output data generated from MS-DIAL analysis was submitted to GNPS for molecular networking analysis with feature-based molecular networking[6] to cluster compounds based on structural similarities.

##### 3.2. Fermentation and Isolation of lentindoles A (1), B (2), and C (3) from *Lentzea* sp. OK19-0192

A 10 mL of seed medium in a 70-mL test tube was inoculated with 100 $\mu$L of the glycerol stock of *Lentzea* sp. OK19-0192 and shaken at 300 rpm for 3 days at 27°C. A 100 mL of seed medium in a 500-mL Erlenmeyer flask was inoculated with 1 mL of the first seed culture broth and shaken at 210 rpm for 6 days at 27°C. A 100 mL production medium in 500-mL Erlenmeyer flasks (total 60 flasks) was inoculated with 1 mL of the second seed culture broth and then rotary shaken at 210 rpm for 6 days at 27°C.

The procedure for isolation of lentindoles A–C (**1**-**3**) was summarized in **Scheme S1**. Compounds **1**-**3** were isolated from a 6-day-old culture broth guided by UV and mass-to-charge ratios. Six litter of the culture broth was centrifuged at 3000 rpm for 10 min. The supernatant was subjected to Diaion HP-20 column (55 i.d. × 200 mm), and compounds were eluted stepwise with 1.5 l MeOH aq. (0, 50, 100% (v/v)). The 100% MeOH fraction was dried *in vacuo*. The residual materials (1.7 g) were dissolved in a small amount of methanol and applied to a silica gel column (37 i.d. × 120 mm). The compounds were eluted stepwise with 300 mL of $CHCl_3$–MeOH (0:1, 100:1, 50:1, 10:1, 1:1, 0:1 (v/v)). The 10:1 fraction was dried *in vacuo*. The crude materials (527.2 mg) were dissolved in a small amount of MeOH and further purified by preparative HPLC equipped with a Pegasil ODS SP100 column (20 i.d. × 250 mm). The compounds were eluted using stepwise solvent with 45% MeOH aq. (0–20 min) and 55% MeOH aq. (20–60 min) at a flow rate of 7.0 mL $min^{-1}$. Compounds were detected at a UV 210 nm. Fractions with retention times of 9.8 min, 35.9 min, and 39.1 min were concentrated *in vacuo* to give lentindoles C (**3**), B (**2**), and a crude extract in this order. The crude extract was dried *in vacuo* and then further purified by an HPLC with 55% MeOH aq. as a mobile phase. Fractions with retention times of 23 min and 26 min were concentrated *in vacuo* to give **2** and **1**, respectively.

#### 3.3. Structure elucidation

##### 3.3.1. Planner structure elucidations by NMR

**Lentindole A (1)**

NMR experiments of the $^{1}H$, $^{13}C$, heteronuclear multiple quantum coherence (HMQC), heteronuclear multiple bond correlation (HMBC), and $^{1}H$-$^{1}H$ homonuclear correlation spectrometry (COSY) in DMSO-$d_6$ were used to elucidate the planar structure of **1** (**Figure S3, S9–S13, Table S6**). The $^{1}H$ and $^{13}C$ NMR and HMQC revealed that **1** had one oxymethylene carbon (C-10; $\delta_C$ 63.7), one quaternary carbon (C-3; $\delta_C$ 75.8), three carbonyl carbons (C-12; $\delta_C$ 171.8, C-16; $\delta_C$ 171.0, C-22; $\delta_C$ 171.9), four *sp*$^3$ methine carbons bonded to a heteroatom (C-9; $\delta_C$ 43.0, C-13; $\delta_C$ 48.1, C-17; $\delta_C$ 51.7), one *sp*$^3$ methine carbon bonded to two heteroatoms (C-2; $\delta_C$ 93.8), four methyl carbons (C-14; $\delta_C$ 18.3, C-20; $\delta_C$ 14.6, C-25; $\delta_C$ 22.3, C-26; $\delta_C$ 22.3), four *sp*$^3$ methylene carbons (C-8; $\delta_C$ 35.9, C-18; $\delta_C$ 31.6, C-19; $\delta_C$ 29.6, C-23; $\delta_C$ 44.4), two aromatic quaternary carbons (C-3a; $\delta_C$ 130.9, C-7a; $\delta_C$ 150.8), four aromatic methine carbons (C-4; $\delta_C$ 123.0, C-5; $\delta_C$ 117.52, C-6; $\delta_C$ 129.1, C-7; $\delta_C$ 109.02), and one *sp*$^3$ methine carbon (C-24; $\delta_C$ 25.6) (**Figure S3, S9–S11, Table S6**). $^{1}H$ NMR and HMQC revealed that five hydrogens (H-1; $\delta_H$ 6.43, H-3b; $\delta_H$ 5.33, H-11; $\delta_H$ 7.89, H-15; $\delta_H$ 7.93, H-21; $\delta_H$ 8.00) were not bonded to any carbon but bonded to either heteroatom (**Figure S3, S9–S11, Table S6**). $^{1}H$-$^{1}H$ COSY showed three spin systems of H-1/H-2 ($\delta_H$ 4.68), H-4 ($\delta_H$ 7.17)/H-5 ($\delta_H$ 6.65)/H-6 ($\delta_H$ 7.06)/H-7 ($\delta_H$ 6.58) and $H_2$-8 ($\delta_H$ 1.87, 2.38)/H-9 ($\delta_H$ 3.36)/$H_2$-10 ($\delta_H$ 3.03, 3.51)/H-11 (**Figure S3, S13**). The HMBC cross-peaks from H-4 to C-3 and C-7a, from H-5 to C-3a, from H-1 to C-3 and C-3a, from H-2 to C-3 and C-7a, from H-7 to C-3a, from H-3b to C-2, C-3 and C-3a, H-2 to C-10, from $H_2$-10 to C-2, and from $H_2$-8 to C-3 and C-3a were observed (**Figure S3, S12**). These results indicated that **1** should have a 6/5/6-fused tricyclic tetrahydropyranoindole (THPyra) moiety (**Figure S3**). Furthermore, the COSY and HMBC spectra indicated the presence of alanine (Ala), methionine (Met), and isovaleric acid (Iva) moieties. The HMBC correlations among those residues from H-11 to C-12, from H-15 to C-16, and from H-21 to C-22 established the sequence of these moieties to be Iva-Met-Ala-THPyra.

**Lentindole B (1)**

NMR experiments of the $^{1}$H, $^{13}$C, HSQC, HMBC, $^{1}$H-$^{1}$H COSY, and rotating frame nuclear Overhauser effect spectroscopy (ROESY) in DMSO-$d_6$, were used to elucidate the planar structure of **2** (**Figure S3, S14–S19, Table S7**). Compound **2** existed as a 1:1 mixture of two conformers (2-1 and 2-2) in DMSO-$d_6$, which were presumed to be rotamers. The NMR spectra of **2** were similar to those of **1**, but differences in chemical shifts of the tricyclic moiety were observed. $^{1}$H-$^{1}$H COSY showed three spin systems of H-1 ($\delta_H$ 6.47 (**2-1**), $\delta_H$ 6.77 (**2-2**))/H-2 ($\delta_H$ 5.18 (**2-1**), $\delta_H$ 5.51 (**2-2**)), H-4 ($\delta_H$ 7.17 (**2-1**), $\delta_H$ 7.19 (**2-2**),)/H-5 ($\delta_H$ 6.60 (**2-1**), $\delta_H$ 6.68 (**2-2**))/H-6 ($\delta_H$ 7.00 (**2-1**), $\delta_H$ 7.05 (**2-2**))/H-7 ($\delta_H$ 6.47 (**2-1**), $\delta_H$ 6.54 (**2-2**)), and $H_2$-8 ($\delta_H$ 2.41 (**2-1**), $\delta_H$ 2.22, 2.38 (**2-2**))/H-9 ($\delta_H$ 3.99 (**2-1**), $\delta_H$ 4.19 (**2-2**))/$H_2$-10 ($\delta_H$ 2.62, 3.29 (**2-1**), $\delta_H$ 2.51, 3.20 (**2-2**))/H-11($\delta_H$ 4.67 (**2-1**), $\delta_H$ 4.53 (**2-2**)) (**Figure S3, S18**). The HMBC cross-peaks from H-6 to C-7a ($\delta_c$ 149.7 (**2-1**), $\delta_c$ 149.1 (**2-2**)), from H-5 to C-3a ($\delta_c$ 132.2 (**2-1**), $\delta_c$ 133.3 (**2-2**)), from H-1 to C-3a, from H-2 to C-3 ($\delta_c$ 88.5 (**2-1**), $\delta_c$ 87.7 (**2-2**)), C-3a and C-7a, from H-3b ($\delta_H$ 5.69 (**2-1**), $\delta_H$ 5.76 (**2-2**)) to C-3, from H-2 to C-9 ($\delta_c$ 59.5 (**2-1**), $\delta_c$ 58.7 (**2-2**)), and from $H_2$-8 to C-3 were observed (**Figure S3, S17**). These results indicated that **2** has a tetrahydropyrroloindole (THPyro) moiety instead of THPyra moiety (**Figure S3**).

##### 3.3.2. Amino acid analysis by advanced Marfey's method

A 1.0 mg powder of compound was added to 200 $\mu$L of 6 M HCl and incubated overnight at 100 °C. Then, the reaction solution was diluted with water and dried *in vacuo*. After the removal of HCl completely, the dried residue was dissolved in 1 mL water. A 5 $\mu$L solution was mixed with 45 $\mu$L distilled water, 20 $\mu$L 1 M $NaHCO_3$ and a 50 $\mu$L 1% (w/v) D-FDLA in acetone and incubated at 37 °C for 1 h. The reaction solution was neutralized by the addition of 25 $\mu$L 1 M HCl and dried *in vacuo*. The residual materials were dissolved in 1 mL acetonitrile and analyzed by LC-HRMS to compare with the D-DLA derivative of amino acid as authentic standard.

##### 3.3.3. Structural analysis of compound 1 using 3D ED/ MicroED

Lentindole A (**1**) was crystallized under suitable conditions (in MeOH at r.t.) and crushed using a pair of microscopy glass slides. Prepared micro-crystals of **1** was used for 3D ED/MicroED measurement. The sample was applied to a grid (microgrid Cu200, JEOL Ltd) and kept at 293 K during data collection. 3D ED/MicroED data were collected (200 keV electrons λ = 0.0251 Å) using an XtaLAB Synergy-ED (Rigaku Co. and JEOL Ltd). The integrated control and processing software of the equipment was CrysAlisPro for ED (ver. 1.171.43.27a) for stage control, diffraction experiments, data integration, and reduction. Three datasets were merged to increase the completeness of the data. The crystal structure was solved using the intrinsic phasing approach with the SHELXT[7] to give the molecular structure and was refined using SHELXL[7] in Olex[8] to refine kinematically. The crystallographic data of lentindole in the standard CIF format file have been deposited at the Cambridge Crystallographic Data Centre (CCDC) as supplementary publication number CCDC 2406780. The crystallographic data were summarized in **Table S8**. Absolute configuration determination of Lentindole A (**1**) by 3D ED/MicroED was performed as previously reported[9–13].

##### 3.4. Synthesis procedures of lentindole C (3) and characterization data

Unless otherwise noted in this synthesis procedures, reactions were carried out in flame- or oven-dried glassware under a positive pressure of $N_2$ in anhydrous solvents using standard Schlenk techniques. Reaction temperatures above room temperature (20–25°C) were controlled by an IKA® temperature modulator or AS ONE Co. Oil Bath and monitored using liquid-in-glass thermometers. Reaction progress was monitored by thin-layer chromatography (TLC) on Sigma Aldrich/Millipore silica gel TLC plates (60 Å, F254 indicator). TLC plates were visualized by exposure to ultraviolet light (254 nm), and/or stained by submersion in aqueous potassium permanganate solution ($KMnO_4$), *p*-anisaldehyde, ceric ammonium molybdate, ninhydrin, or phosphomolybdic acid stains and heating with a heat gun or heating plate. Organic solutions were concentrated under reduced pressure on an EYELA temperature-controlled rotary evaporator equipped with a cooling condenser. Flash column chromatography was performed with glass columns using Kanto Chemical silica gel (60 N, spherical neutral, 40–50 $\mu$m particle size), using ACS grade solvents. All yields refer to spectroscopically ($^1$H and $^{13}$C NMR) pure material. High-resolution mass spectra (HRMS) were measured on a JEOL JMS-AX505HA, JEOL JMS-700 MStation and/or JEOL JMS-T100LP. Data acquisition and processing were performed using the XcaliburTM software. Optical rotations were measured on a JASCO P-2020 polarimeter.

**Compound 5**

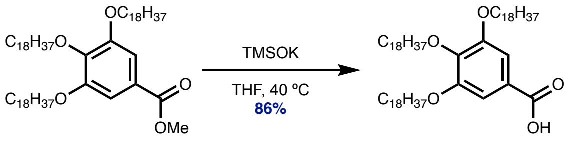

4 5

To a solution of **4** (94.1 mg, 0.10 mmol) in THF (2.00 mL, 0.05 M) was added TMSOK (38.5 mg, 3.0 equiv) at room temperature. After stirring at 40 ºC for 21 h, to the reaction mixture was added TBAF (1.0 M in THF, 200 $\mu$L, 0.20 mmol, 2.0 equiv) at 40 ºC. After stirring at 40 ºC for 2 h, the reaction mixture was cooled down to room temperature and poured into cold MeOH (100 mL, 0.02 M). The resulting suspension was stirred for 30 min at 0 ºC and filtered through a pad of Celite (washed with excess MeOH). The product collected on the filter cake was dissolved in $CHCl_3$ and then eluted by suction filtration using excess $CHCl_3$. The resulting filtrate was concentrated *in vacuo*. The resulting filtrate was concentrated *in vacuo* and dried under high vacuum, yielding **5** (80.1 mg, 86.4 $\mu$mol, 86%, NMR purity ≥95%) a colorless powder.

**Rf-value**: 0.62 (hexane/EtOAc = 2:1, stained with phosphomolybdic acid)

$^{1}$**H NMR** (500 MHz, $CDCl_3$, 45 ºC): δ 7.33 (br-s, 2H), 4.07-4.01 (m, 6H), 1.84-1.74 (m, 6H), 1.48 (d, $J$ = 7.4 Hz, 6H), 1.28 (s, 84H), 0.89 (t, $J$ = 6.5 Hz, 9H)

**COOH proton was not detected.*

$^{13}$**C NMR** (126 MHz, $CDCl_3$, 45 ºC): δ 171.5, 152.9, 143.5, 123.7, 109.0, 73.6, 69.4 (2C), 31.9 (3C), 30.4, 29.7- 29.4 (overlap, 39C), 26.1 (3C), 22.7 (3C), 14.0 (3C)

*Note: Fast Atom Bombardment (FAB) mass spectra of 5 was difficult to identify.*

**Compound 6**

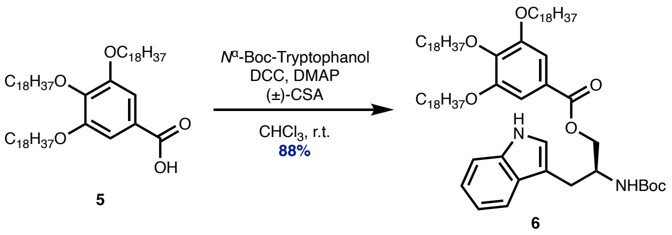

To a solution of **5** (253 mg, 0.27 mmol) in $CHCl_3$ (5.46 mL, 0.05 M) was added $N^{\alpha}$-Boc-Tryptophanol (95.0 mg, 0.33 mmol, 1.2 equiv), (±)-CSA (6.3 mg, 27.3 $\mu$mol, 0.1 equiv), DCC (124 mg, 0.60 mmol, 2.2 equiv) and DMAP (10.0 mg, 81.8 $\mu$mol, 0.3 equiv) at room temperature. After stirring at room temperature for 20 h, the reaction mixture was cooled down to 0 °C and poured into cold MeOH (27.3 mL, 0.01 M). The resulting suspension was stirred for 30 min at 0 ºC and filtered through a pad of Celite (washed with excess MeOH). The product collected on the filter cake was dissolved in $CHCl_3$ and then eluted by suction filtration using excess $CHCl_3$. The resulting filtrate was concentrated *in vacuo*. The resulting residue was purified by silica gel flash column chromatography (hexane:EtOAc = 10:1 to 4:1 gradient), yielding **6** (290 mg, 0.241 mmol, 88%) as a colorless powder.

**Rf-value**: 0.57 (hexane/EtOAc = 3:1, stained with phosphomolybdic acid)

$[\alpha]_D^{25}$ = +7.00 (c = 0.1, $CHCl_3$)

**$^1$H NMR** (500 MHz, $CDCl_3$): δ 8.43 (s, 1H), 7.67 (d, *J* = 8.0 Hz, 1H), 7.35 (d, *J* = 8.0 Hz, 1H), 7.30 (s, 2H), 7.19 (t, *J* = 7.5 Hz, 1H), 7.12 (t, *J* = 7.3 Hz, 1H), 7.04 (br-s, 1H), 4.86 (d, *J* = 8.0 Hz, 1H), 4.39 (br-s, 1H), 4.31 (d, *J* = 4.5 Hz, 2H), 4.07-3.97 (m, 6H), 3.14-3.03 (m, 2H), 1.85-1.74 (m, 6H), 1.52-1.46 (m, 6H), 1.43 (s, 9H), 1.36-1.16 (m, 84H), 0.90 (t, *J* = 6.8 Hz, 9H)

**$^{13}$C NMR** (126 MHz, $CDCl_3$): δ 166.4, 155.4, 152.8, 142.5, 136.2, 127.6, 124.4, 122.8, 122.0, 119.5, 118.7, 111.2, 110.9, 108.1, 79.4, 73.5, 69.1 (2C), 65.8, 60.4, 50.2, 31.9 (3C), 30.3, 29.7- 29.3 (overlap, 39C), 28.3 (3C), 27.5, 26.1 (2C), 26.0, 22.7 (3C), 14.1 (3C).

*Note: Fast Atom Bombardment (FAB) mass spectra of 6 was difficult to identify.*

**Compound 9**

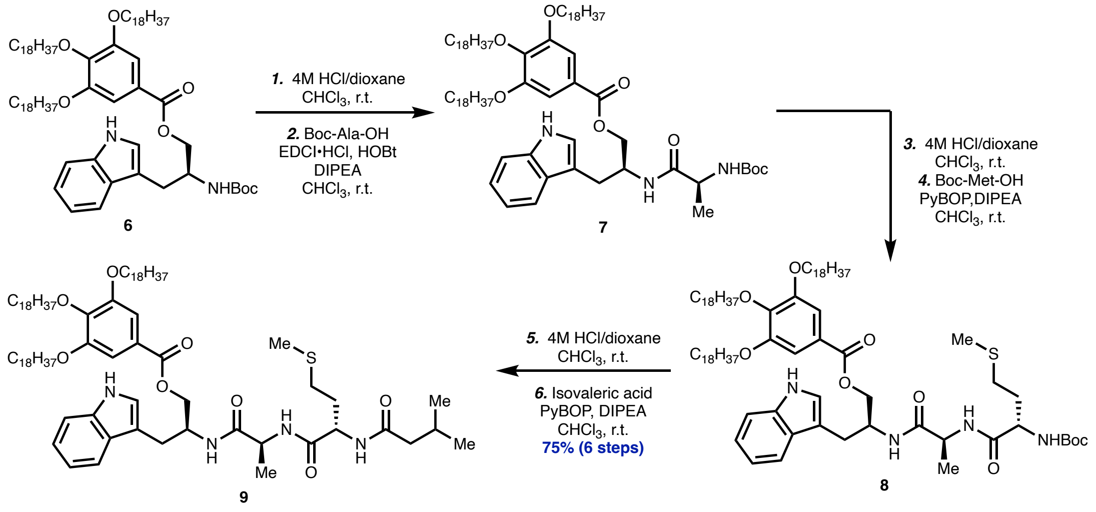

To a solution of **6** (152 mg, 0.13 mmol) in $CHCl_3$ (1.27 mL, 0.1M) was added 4M HCl/dioxane (1.27 mL, 0.1M) at room temperature. After stirring at room temperature for 2 h, the reaction mixture was concentrated *in vacuo* and dried under high vacuum. To this crude material (de-Boc compound) was added $CHCl_3$ (1.01 mL, 0.13 M), Boc-Ala-OH (26.4 mg, 0.14 mmol, 1.1 equiv), HOBt (5.1 mg, 38.0 $\mu$mol, 0.3 equiv), EDCI·HCl (28.2 mg, 0.15 mmol, 1.2 equiv) and DIPEA (50.8 $\mu$L, 0.29 mmol, 2.3 equiv) at room temperature. After stirring at room temperature for 5 h, the reaction mixture was washed with 1 M HCl and brine and concentrated *in vacuo*. This crude material **7** was used in the next reaction without further purification.

To a solution of crude material **7** in $CHCl_3$ (1.27 mL, 0.1M) was added 4M HCl/dioxane (1.27 mL, 0.1M) at room temperature. After stirring at room temperature for 1 h, the reaction mixture was concentrated *in vacuo* and dried under high vacuum. To this crude material (de-Boc compound) was added $CHCl_3$ (2.53 mL, 0.05 M), Boc-Met-OH (37.9 mg, 0.15 mmol, 1.2 equiv), PyBOP (98.9 mg, 0.19 mmol, 1.5 equiv) and DIPEA (66.2 μL, 0.38 mmol, 3.0 equiv) at room temperature. After stirring at room temperature for 11 h, the reaction mixture was cooled down to 0 °C and poured into cold MeOH (12.7 mL, 0.01 M). The resulting suspension was stirred for 30 min at 0 ºC and filtered through a pad of Celite (washed with excess MeOH). The product collected on the filter cake was dissolved in $CHCl_3$ and then eluted by suction filtration using excess $CHCl_3$. The resulting filtrate was concentrated *in vacuo* and dried under high vacuum, yielding **8** (163 mg, theoretically 178 mg) a pale-yellow powder. Compound **8** was used in the next reaction without further purification.

**A portion was used from the compound **8**.*

To a solution of crude material **8** (150 mg, 0.11 mmol) in $CHCl_3$ (1.07 mL, 0.1M) was added 4M HCl/dioxane (1.07 mL, 0.1M) at room temperature. After stirring at room temperature for 1 h, the reaction mixture was concentrated *in vacuo* and dried under high vacuum. To this crude material (de-Boc compound) was added $CHCl_3$ (2.14 mL, 0.05 M), isovaleric acid (14.1 μL, 0.13 mmol, 1.2 equiv), PyBOP (83.4 mg, 0.16 mmol, 1.5 equiv) and DIPEA (55.9 μL, 0.32 mmol, 3.0 equiv) at room temperature. After stirring at room temperature for 2 h, the reaction mixture was cooled down to 0 °C and poured into cold MeOH (10.7 mL, 0.01 M). The resulting suspension was stirred for 30 min at 0 ºC and filtered through a pad of Celite (washed with excess MeOH). The product collected on the filter cake was dissolved in $CHCl_3$ and then eluted by suction filtration using excess $CHCl_3$. The resulting filtrate was concentrated *in vacuo* and dried under high vacuum, yielding **9** (119 mg, 87.3 μmol, 82%, 75% from **9**) a pale-yellow powder.

**Rf-value**: 0.55 ($CHCl_3$/MeOH = 30:1, stained with phosphomolybdic acid)

**HRMS** (*m/z*): FAB [M+Na]$^+$ calculated for $C_{85}H_{148}N_4NaO_8S$: 1408.0916, found: 1408.0934.

**$^1$H NMR** (500 MHz, $CDCl_3$): δ 8.36 (br-s, 1H), 7.66 (d, *J* = 8.0 Hz, 1H), 7.37 (d, *J* = 8.0 Hz, 1H), 7.26 (s, 2H,* overlap with the $CHCl_3$ peak), 7.20 (t, *J* = 7.5 Hz, 1H), 7.12 (t, *J* = 7.5 Hz, 1H), 7.07 (s, 1H), 6.84 (br-s, 1H), 6.57 (br-s, 1H), 6.35 (br-s, 1H), 4.65 (m, 1H), 4.51 (m, 1H), 4.38-4.33 (m, 3H), 4.02-3.96 (m, 6H), 3.12-3.03 (m, 2H), 2.50-2.44 (m, 2H), 2.13-2.02 (m, 7H), 1.94 (m, 1H), 1.82-1.73 (m, 6H), 1.51-1.41 (m, 6H), 1.26-1.14 (m, 87H), 0.95-0.87 (m, 15H)

*Note: Compound 9 has poor solubility in various solvents and does not form a homogeneous solution, so $[\alpha]_D$ and $^{13}$C NMR were difficult to measure.*

**Compound 10**

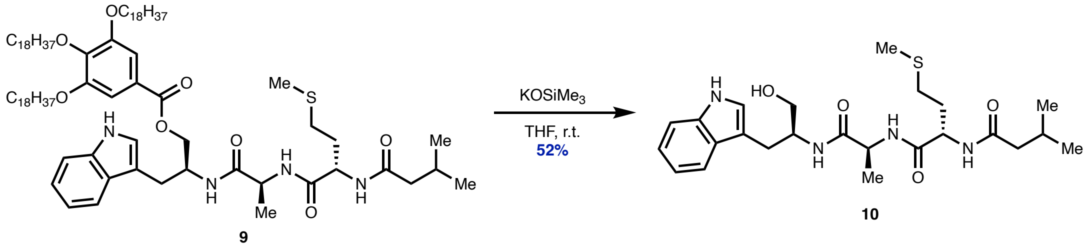

To a solution of crude material **9** (13.6 mg, 10.0 $\mu$mol) in $CHCl_3$ (0.2 mL, 0.05M) was added $KOSiMe_3$ (1.9 mg, 15.0 $\mu$mol, 1.5 equiv) at room temperature. After stirring at room temperature for 5 h, the reaction mixture was poured into cold MeOH (1.0 mL, 0.01 M). The resulting suspension was stirred for 30 min at 0 °C and filtered through a pad of Celite (washed with excess MeOH). The resulting filtrate was *in vacuo*. The resulting residue was purified by thin-layer preparative TLC ($CHCl_3$/MeOH = 10:1), yielding **10** (2.5 mg, 5.2 $\mu$mol, 52%) as yellow oil.

**Rf-value**: 0.49 ($CHCl_3$/MeOH = 10:1, stained with phosphomolybdic acid)

**HRMS** (*m/z*): ESI [M+H]$^+$ calculated for $C_{24}H_{37}N_4O_4S$: 477.2536, found: 477.2534.

**$[\alpha]_D^{25}$** = –13.2 (c = 0.1, $CHCl_3$)

**$^1$H NMR** (500 MHz, DMSO-$d_6$): δ 10.77 (s, 1H), 8.06 (d, *J* = 8.0 Hz, 1H), 7.94 (d, *J* = 7.0 Hz, 1H), 7.69 (d, *J* = 8.0 Hz, 1H), 7.60 (d, *J* = 8.0 Hz, 1H), 7.31 (d, *J* = 8.0 Hz, 1H), 7.10 (d, *J* = 1.5 Hz, 1H), 7.04 (t, *J* = 7.0 Hz, 1H), 6.95 (t, *J* = 7.5 Hz, 1H), 4.70 (br-s, 1H), 4.32 (dt, *J* = 13.0, 5.0 Hz, 1H), 4.23 (m, 1H), 3.91 (m, 1H), 3.33-3.30 (m, 2H), 2.87 (dd, *J* = 14.0, 7.5 Hz, 1H), 2.75 (dd, *J* = 14.0, 7.5 Hz, 1H), 2.47-2.41 (m, 2H), 2.04-1.99 (m, 5H), 1.98-1.85 (m, 3H), 1.76 (m, 1H), 1.18 (d, *J* = 7.0 Hz, 3H), 0.86-0.84 (m, 6H)

**$^{13}$C NMR** (126 MHz, DMSO-$d_6$): δ 171.9, 171.8, 170.9, 136.1, 127.5, 123.3, 120.8, 118.5, 118.2, 111.3, 111.1, 62.0, 51.7, 48.3, 44.4, 40.0, 31.6, 29.6, 26.4, 25.6, 22.3, 22.3, 18.4, 14.6.

#### 4. Feeding of labeled amino acids to analyze the biosynthesis pathway of lentindole

6-fluoro-DL-tryptophanol (β-Amino-6-fluoro-1H-indole-3-propanol) was prepared by following the literature procedure (WO2008001076 A1 2008-01-03)[14]. A 10 mL of seed medium in a 70-mL test tube was inoculated with 100 μL of the glycerol stock of *Lentzea* sp. OK19-0192 and shaken at 300 rpm for 2 days at 27°C. The production media were inoculated with 200 μL of the seed culture and supplemented with 500 μL of 1 M 6-fluoro-DL-tryptophan, L-alanine-$d_4$,$^{13}$C-L-methionine or 6-fluoro-DL-tryptophanole. Then, the test tubes were shaken at 300 rpm for 6 days at 27°C. Each culture broth was mixed with an equal amount of ethanol and centrifuged to remove cells. The supernatants were analyzed by LC-HRMS.

#### 5. Genome analysis

The genomic DNA of *Lentzea* sp. OK19-0192 was obtained by using the cetyltrimethylammonium bromide method. The genomes were sequenced using the Pacbio RSII technology (Macrogen, Inc., Seoul, Republic of Korea). The genome comprised a linear 10,360,722 bp and was annotated using DFAST. BGC in the genome was analyzed via antiSMASH6.0.0.[3]

#### 6. Phylogenetic analysis

The 16S rRNA gene was amplified by PCR using appropriate primer pairs (forward primer: 11F; 5′-AGTTTGATCATGGCTCAG-3′ and 520F; 5′-CGTCAATTCATTTGAGTT-3′ and reverse primer: 925R;

5′-CGTCAATTCATTTGAGTT-3′, 1510R; 5′-GGTTACCTTGTTACGACT-3′ and 1542R; 5′-AAGGAGGTGATCCAGCCGCA-3′). The PCR products were sequenced by Eurofins Genomics. Phylogenetic tree was constructed using neighbor-joining method[15] in the software package MEGA (version 11)[16]. Evolutionary distance matrices were generated according to Kimura's two-parameter model[17].

#### Supplementary Figure

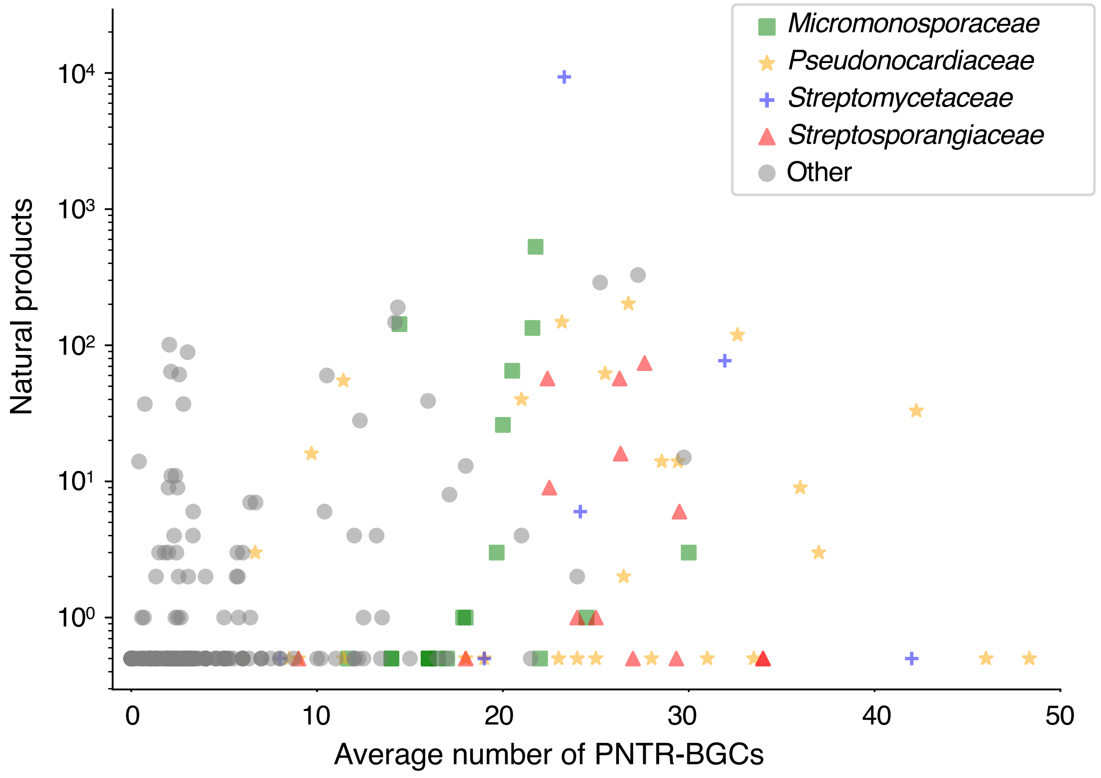

**Figure S1**. Number of NPs and average number of PNTR-BGCs of each genus in *Actinomycetota*. The number of NPs was estimated based on the data from the Dictionary of Natural Products[18] displayed in logarithmic scale. A dot indicates a genus, and their colors follow the legend shown in the figure; green: *Micromonosporaceae*, yellow: *Pseudonocardiaceae*, blue: *Streptomycetaceae*, red: *Streptosporangiaceae*, gray: other family.

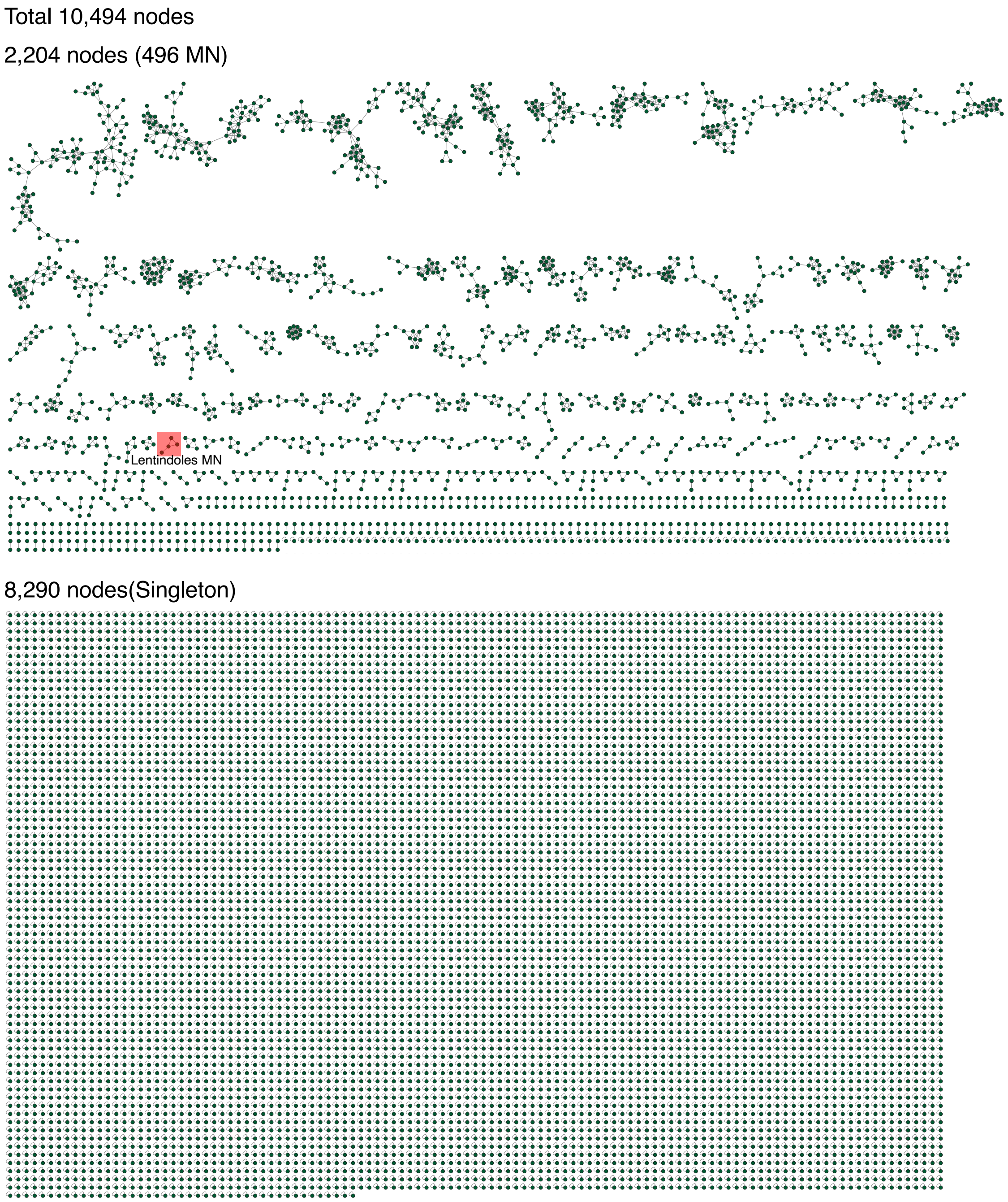

**Figure S2**. Overview of molecular networks.

Total 10,494 nodes were detected, 2,204 nodes were formed 496 MNs, and 8,290 nodes were singletons.

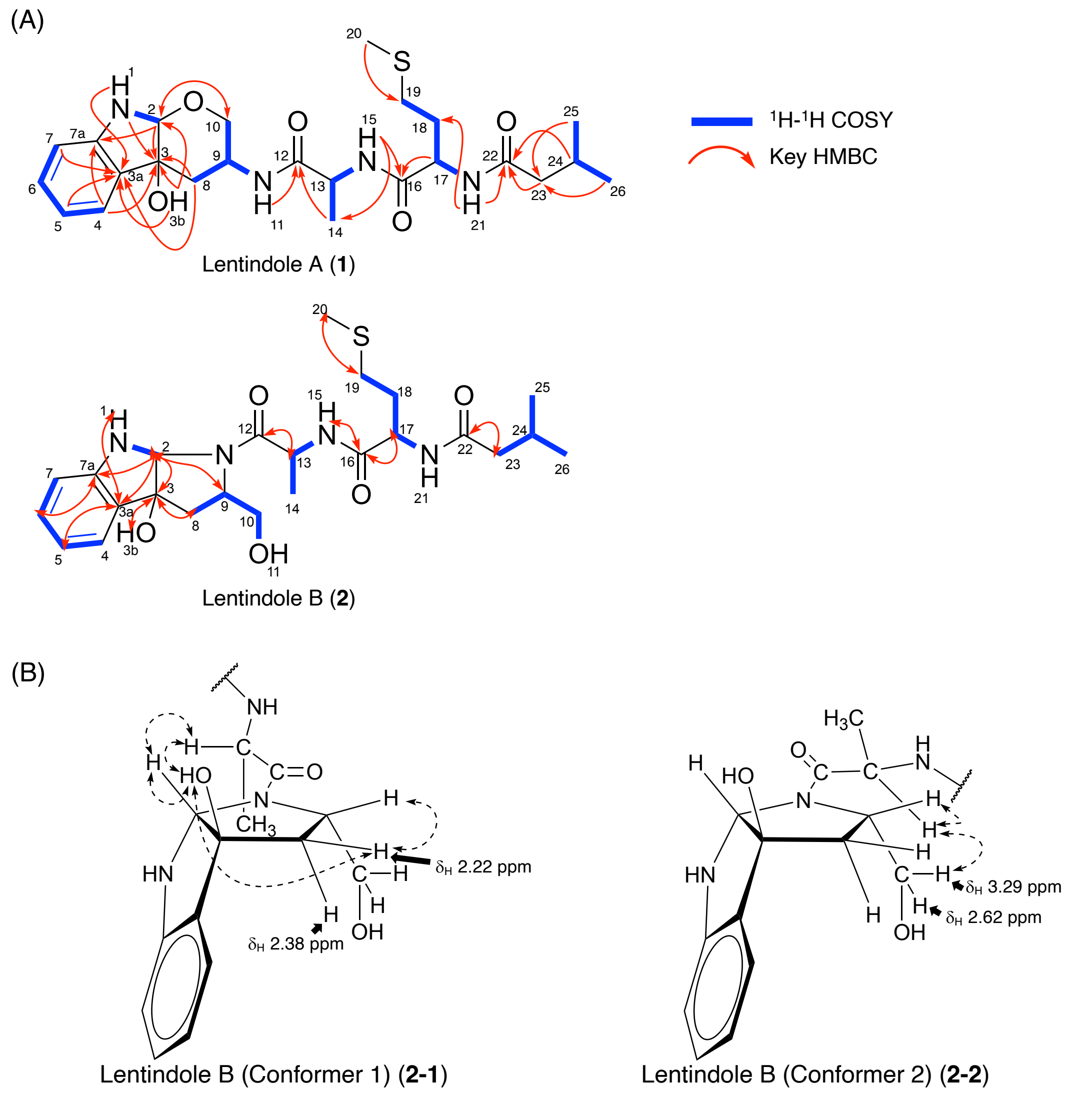

**Figure S3**. Planner structure elucidations of lentindoles A (**1**) and B (**2**) by 2D NMR experiments.

(A) COSY and key HMBC correlations of lentindoles A (**1**) and B (**2**).

(B) ROESY correlations of the two conformers of lentindole B (**2**).

(A)

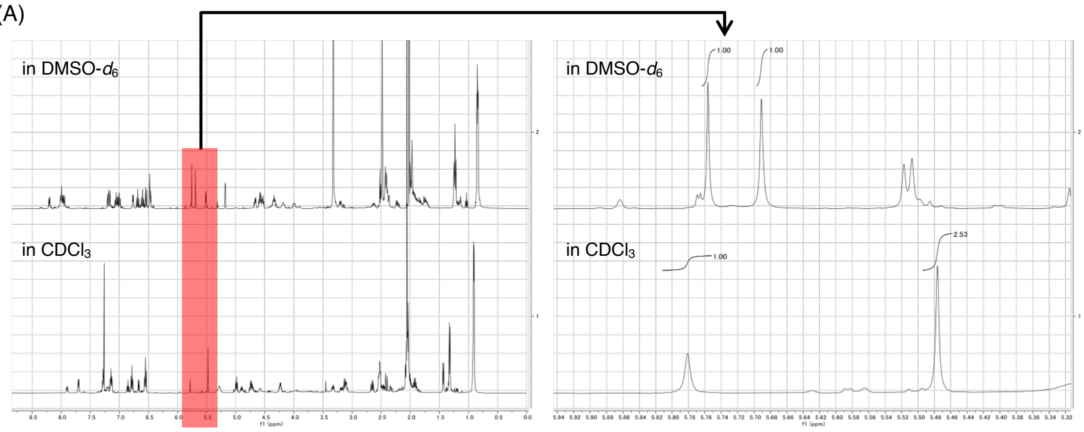

(B)

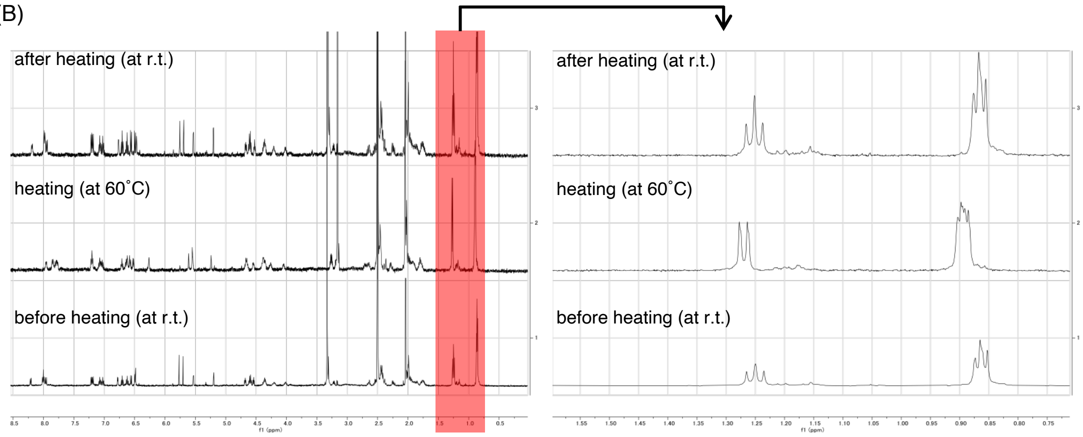

**Figure S4.** Changes in $^1$H NMR (500 MHz) of lentindole B (**2**) spectra under different temperature and solvent conditions.

(A) The integration ratio of the two hydroxy proton signals (H-3b; $\delta_H$ 5.69 (**2-1**), $\delta_H$ 5.76 (**2-2**)) in DMSO-$d_6$ was 1:1, but 2.5:1 ($\delta_H$ 5.47, $\delta_H$ 5.78 (conformer unassigned)) in $CD_3Cl$, showing that the conformer ratio changes depending on the solvent used for the $^1$H NMR measurement.

(B) $^1$H NMR measurements were performed at r.t., followed by heating to 60 °C, and then again after cooling back to r.t. in DMSO-$d_6$. While the methyl proton signals of the alanine moiety ($H_3$-14; $\delta_H$ 1.22 (**2-1**), $\delta_H$ 1.24 (**2-2**)) appeared as two overlapping doublets like a triplet signal at r.t., these signals merged into a single doublet at 60 °C as they moved closer together. Upon cooling back to r.t., these signals reverted to two overlapping doublets like a triplet signal.

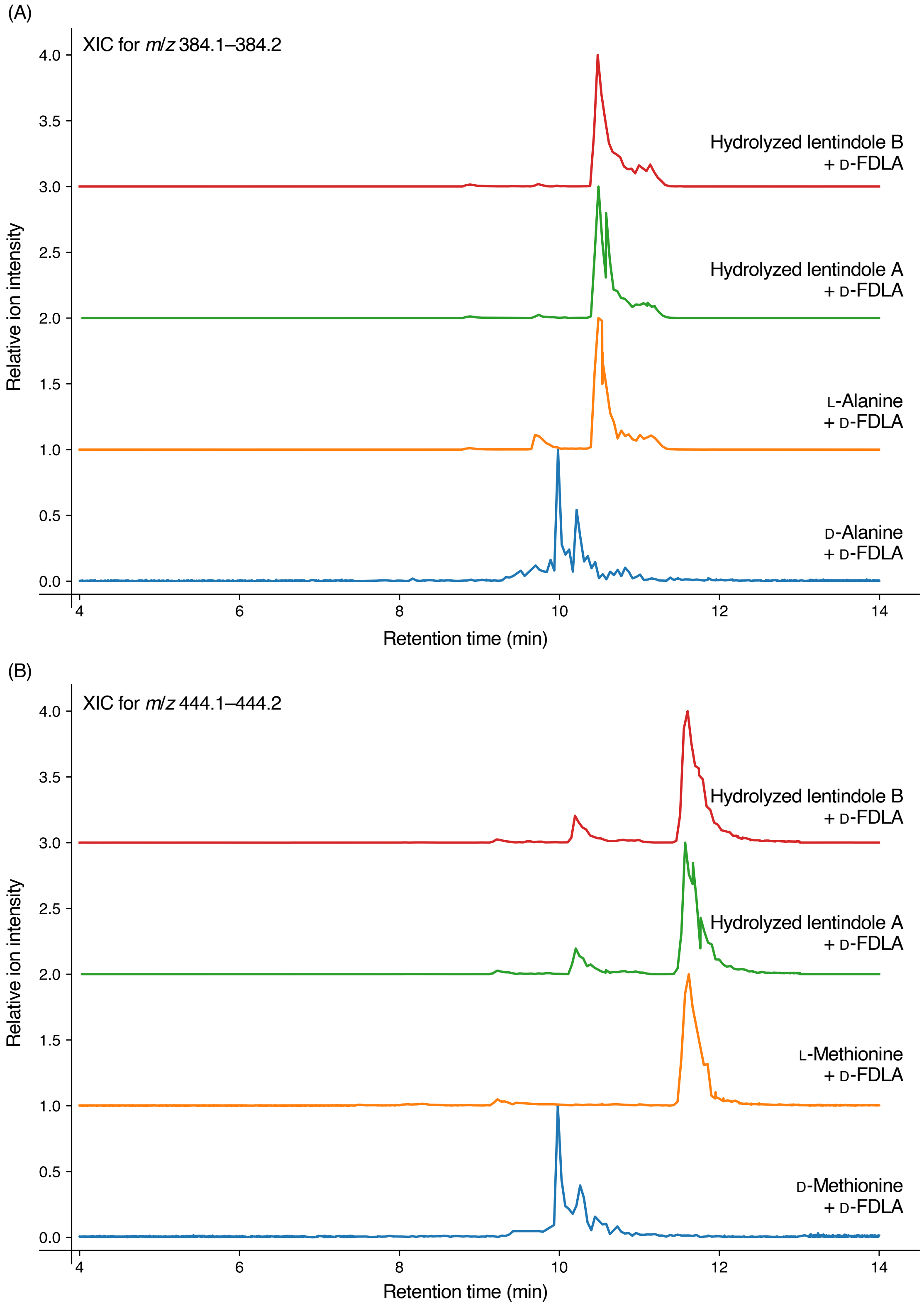

**Figure S5.** Extracted ion chromatogram (XIC) of D-FDLA derivatives.

(A) XIC for *m*/*z* 384.1–384.2

(B) XIC for *m*/*z* 444.1–444.2.

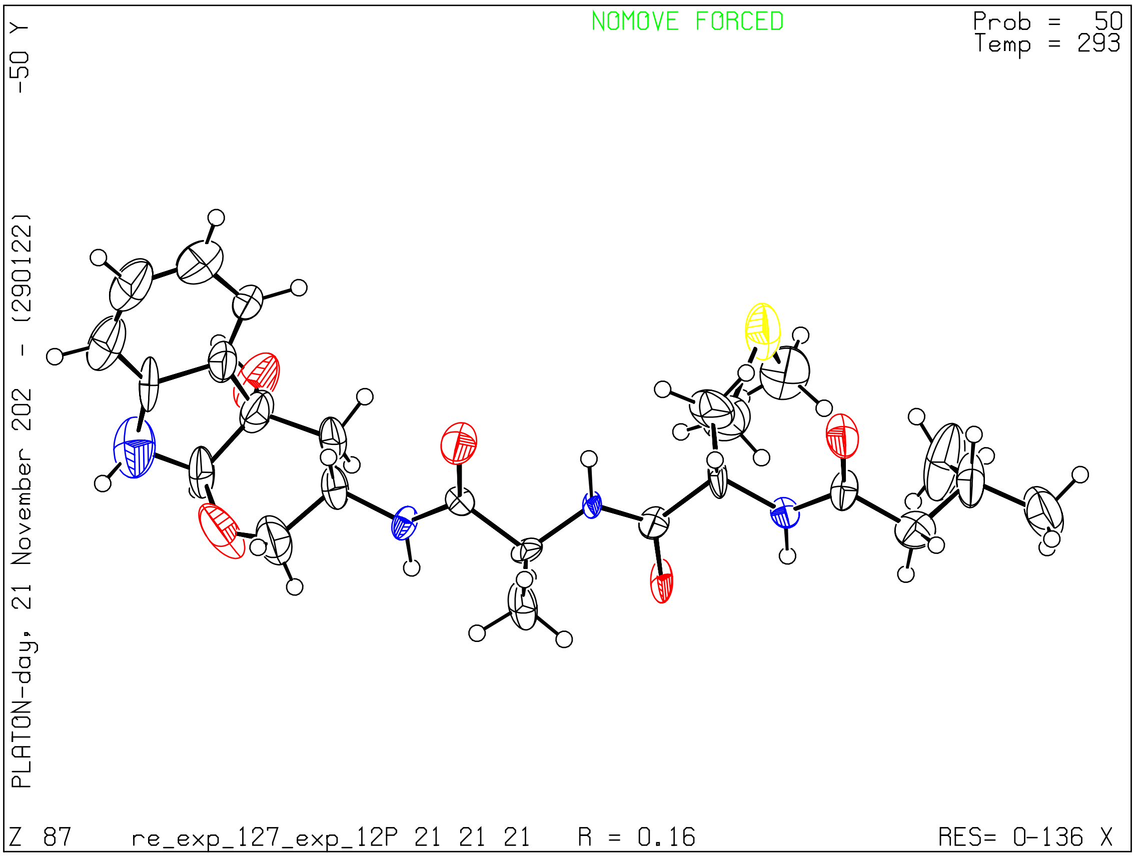

**Figure S6.** ORTEP of lentindole A (**1**)

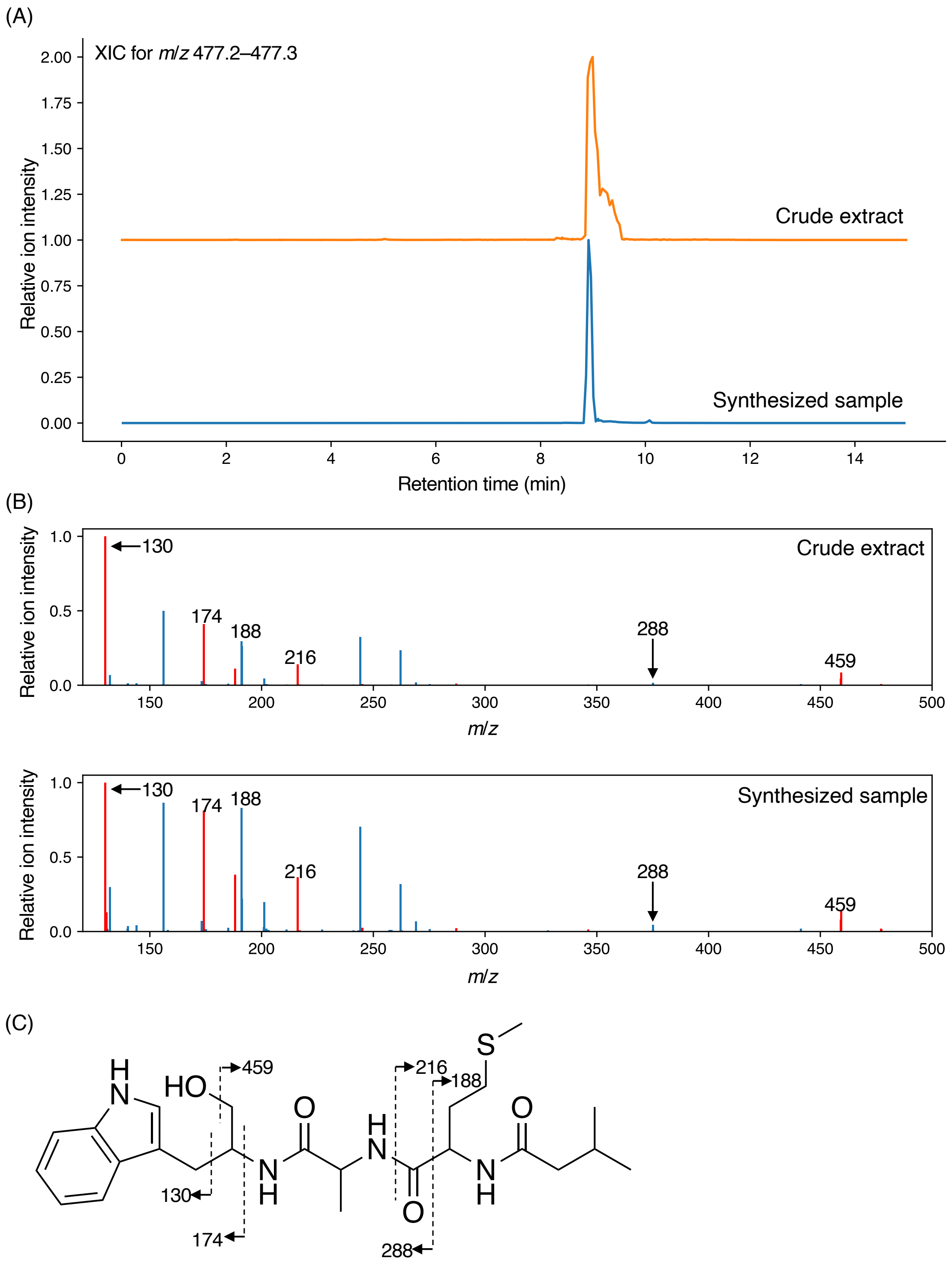

**Figure S7**. Mass spectrometry analysis of lentindole C (**3**) and synthetic **3**

(A) XIC for *m*/*z* 477.2–477.3 of crude extract and synthesized sample.

(B) MS/MS spectra of crude extract and synthesized sample.

(C) MS/MS fragments assignment of **3**.

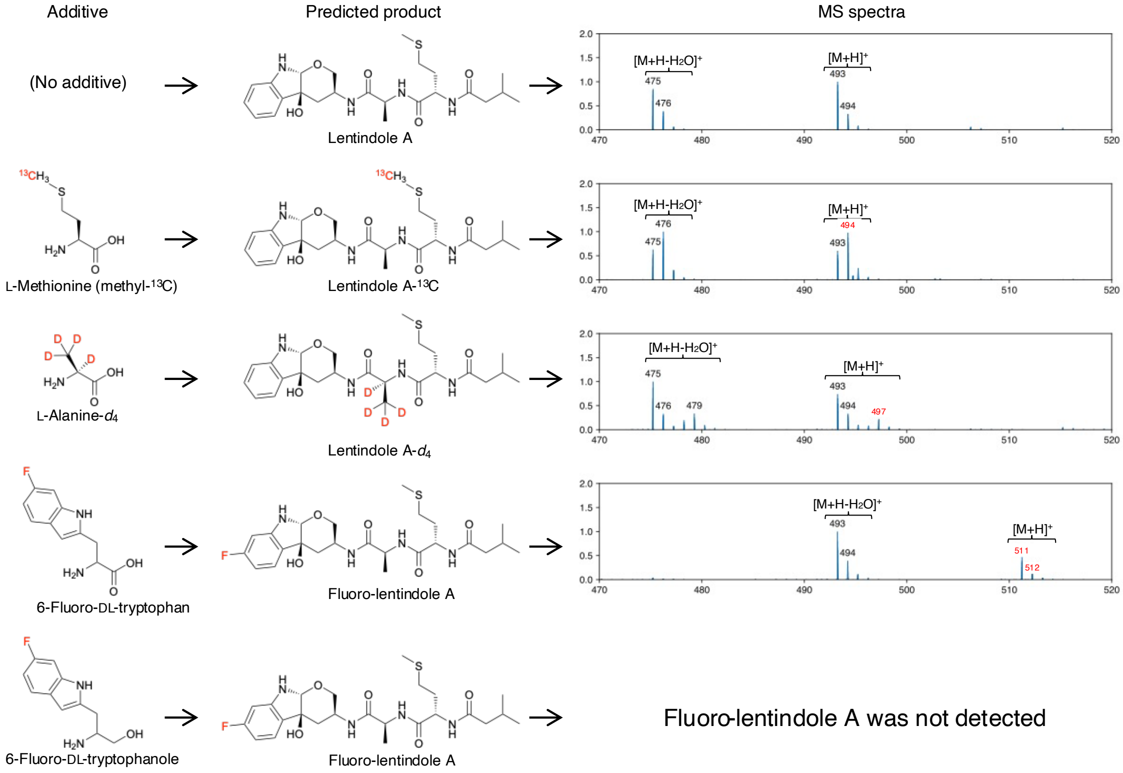

**Figure S8.** Detection of labeled lentindole A (**1**)

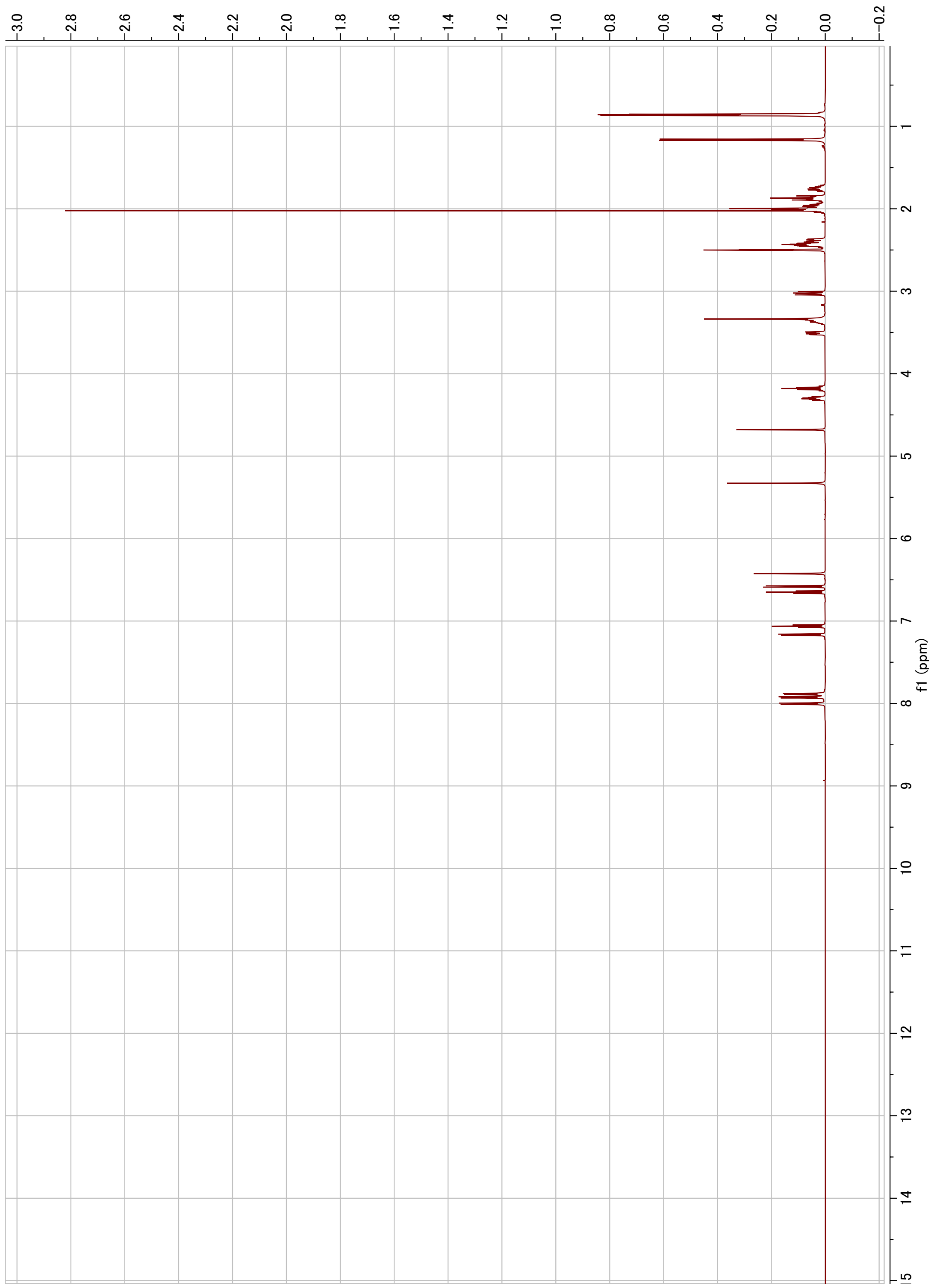

**Figure S9.** $^{1}$H NMR of lentindole A (1) (500 MHz, DMSO-$d_6$)

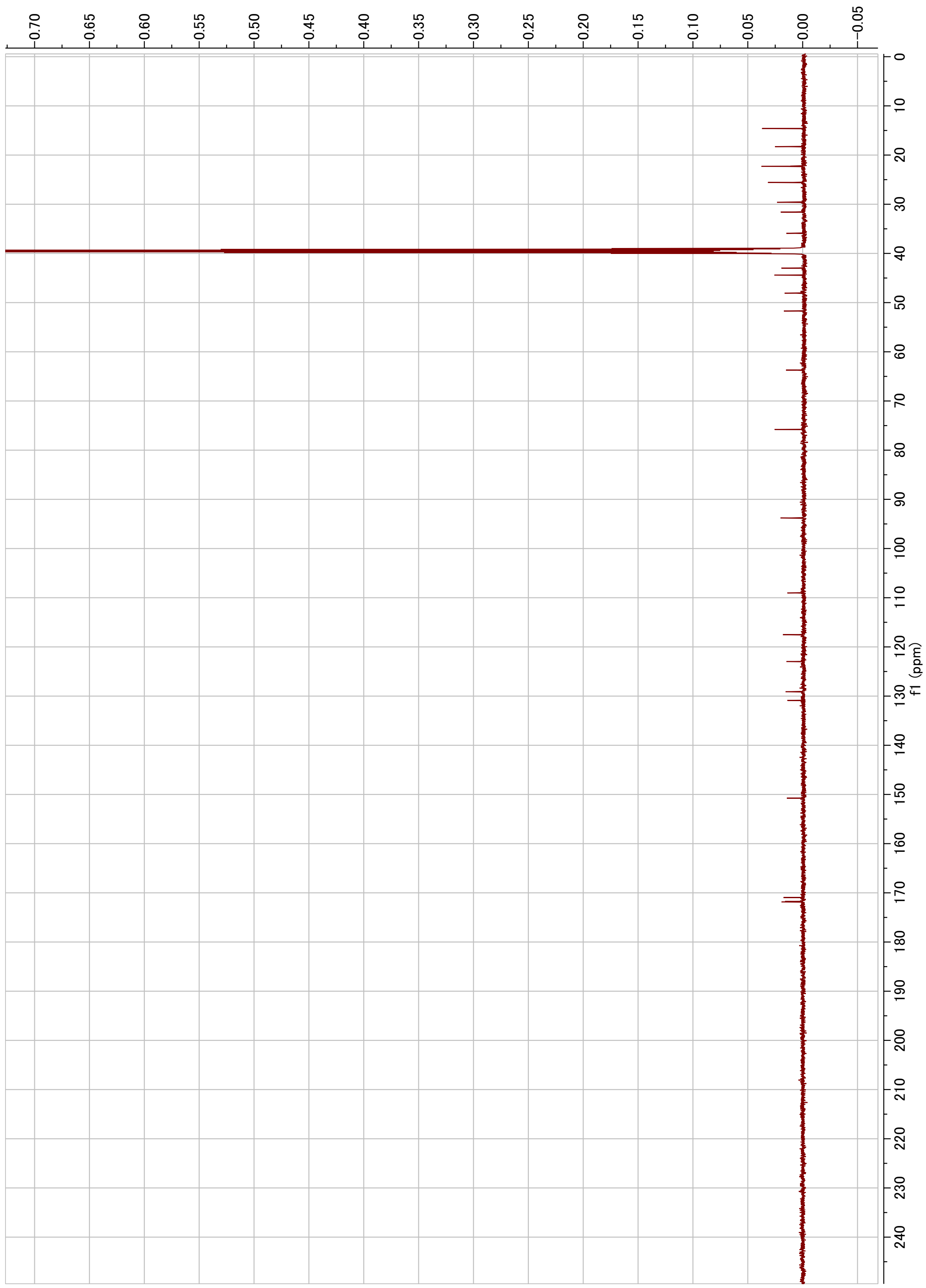

**Figure S10.** $^{13}$C NMR of lentindole A (1) (125 MHz, DMSO-$d_6$)

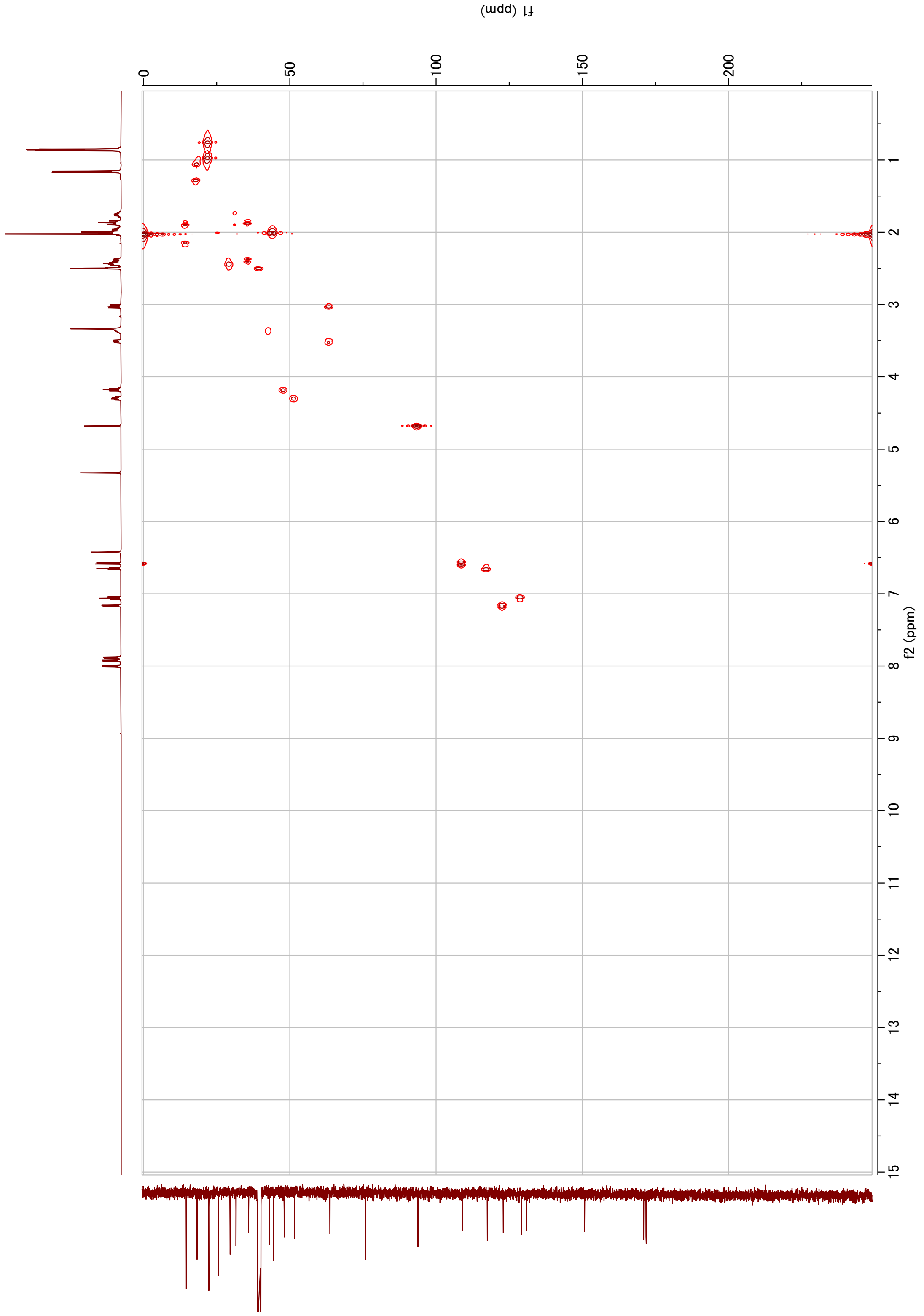

**Figure S11.** HMQC of lentindole A (1) ($^{1}$H:500 MHz, $^{13}$C:125 MHz, DMSO-$d_6$)

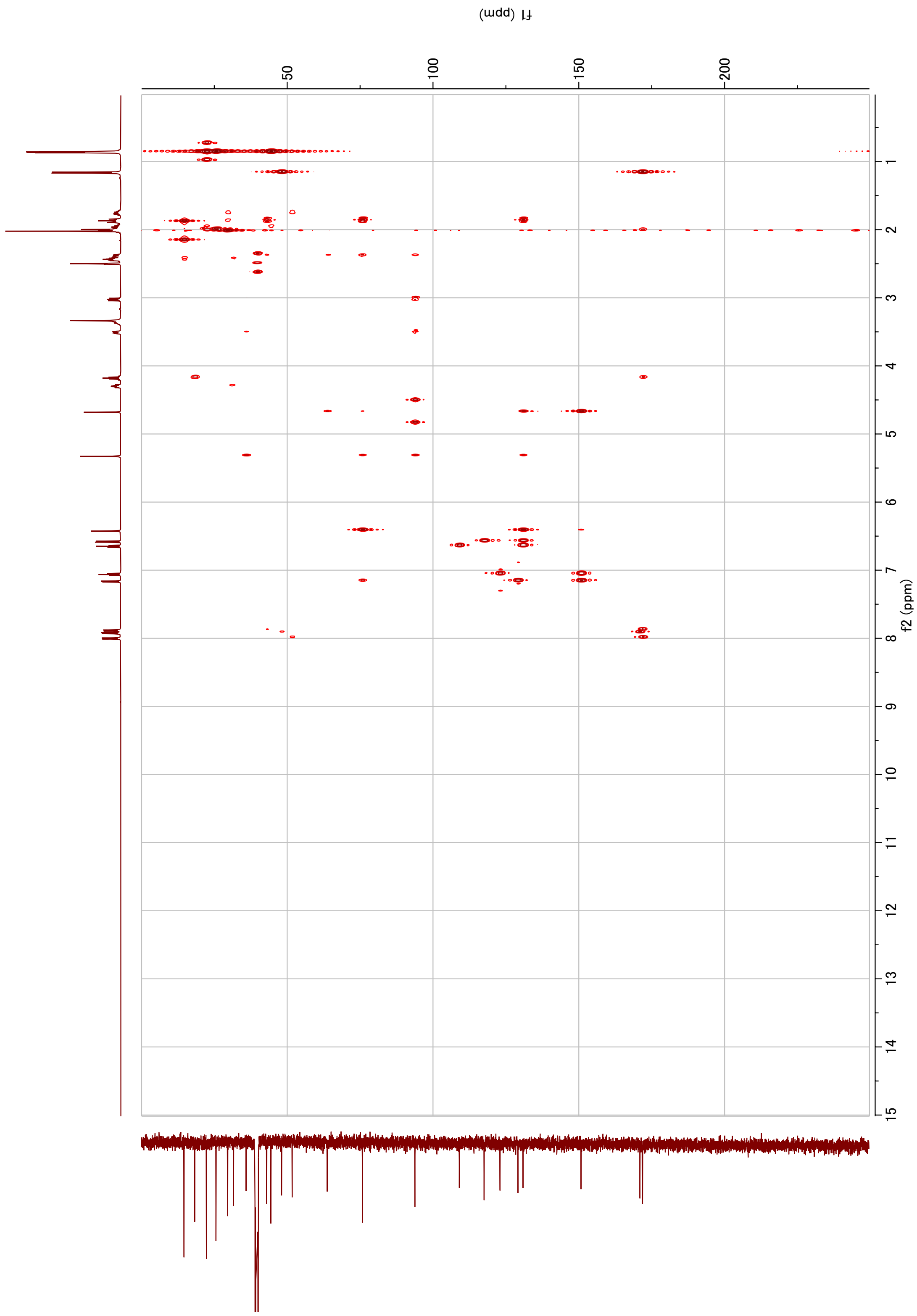

**Figure S12.** HMBC of lentindole A (1) ($^{1}$H:500 MHz, $^{13}$C:125 MHz, DMSO-$d_6$)

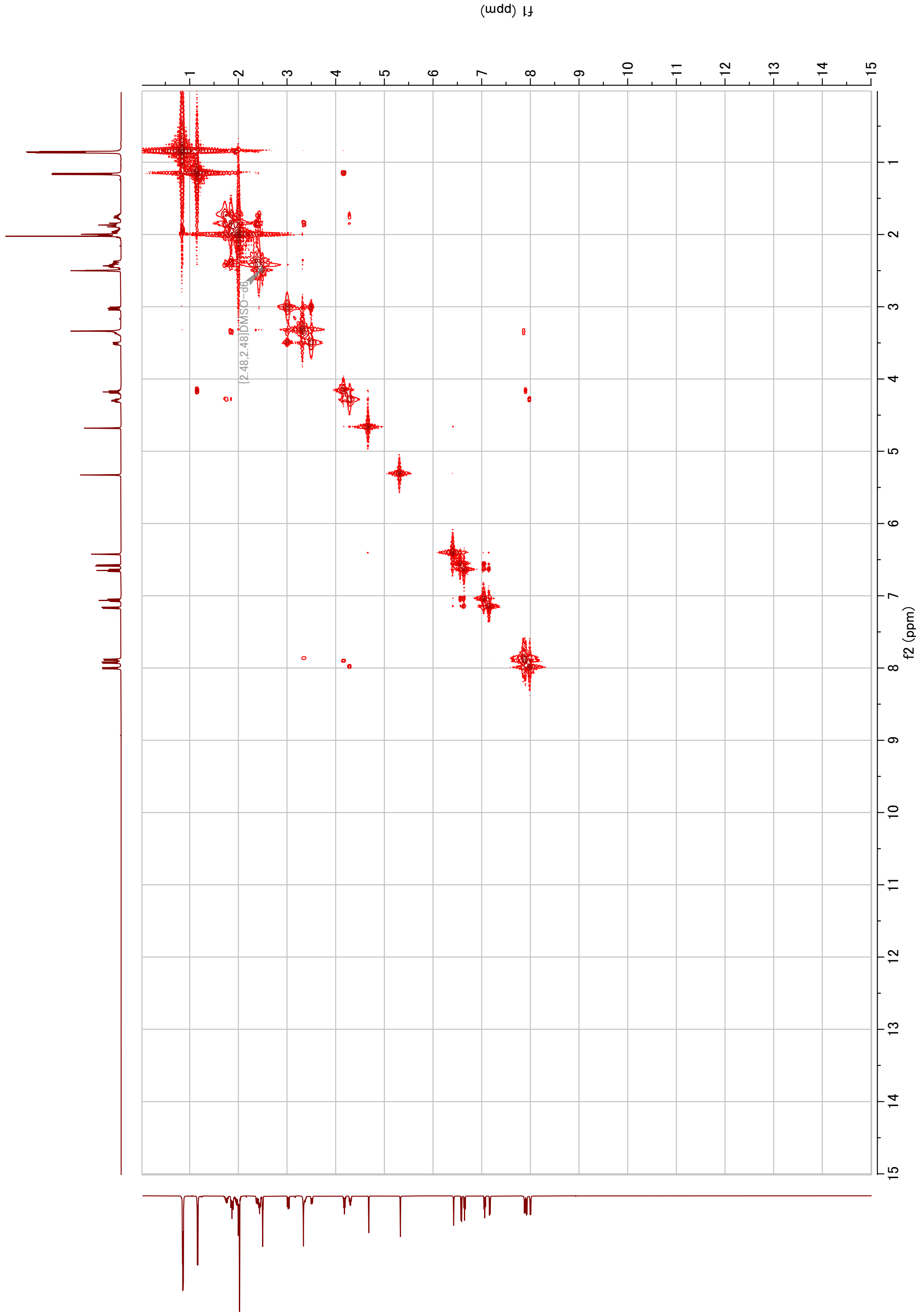

**Figure S13.** $^{1}H$-$^{1}H$ COSY of lentindole A (**1**) (500 MHz, DMSO-$d_6$)

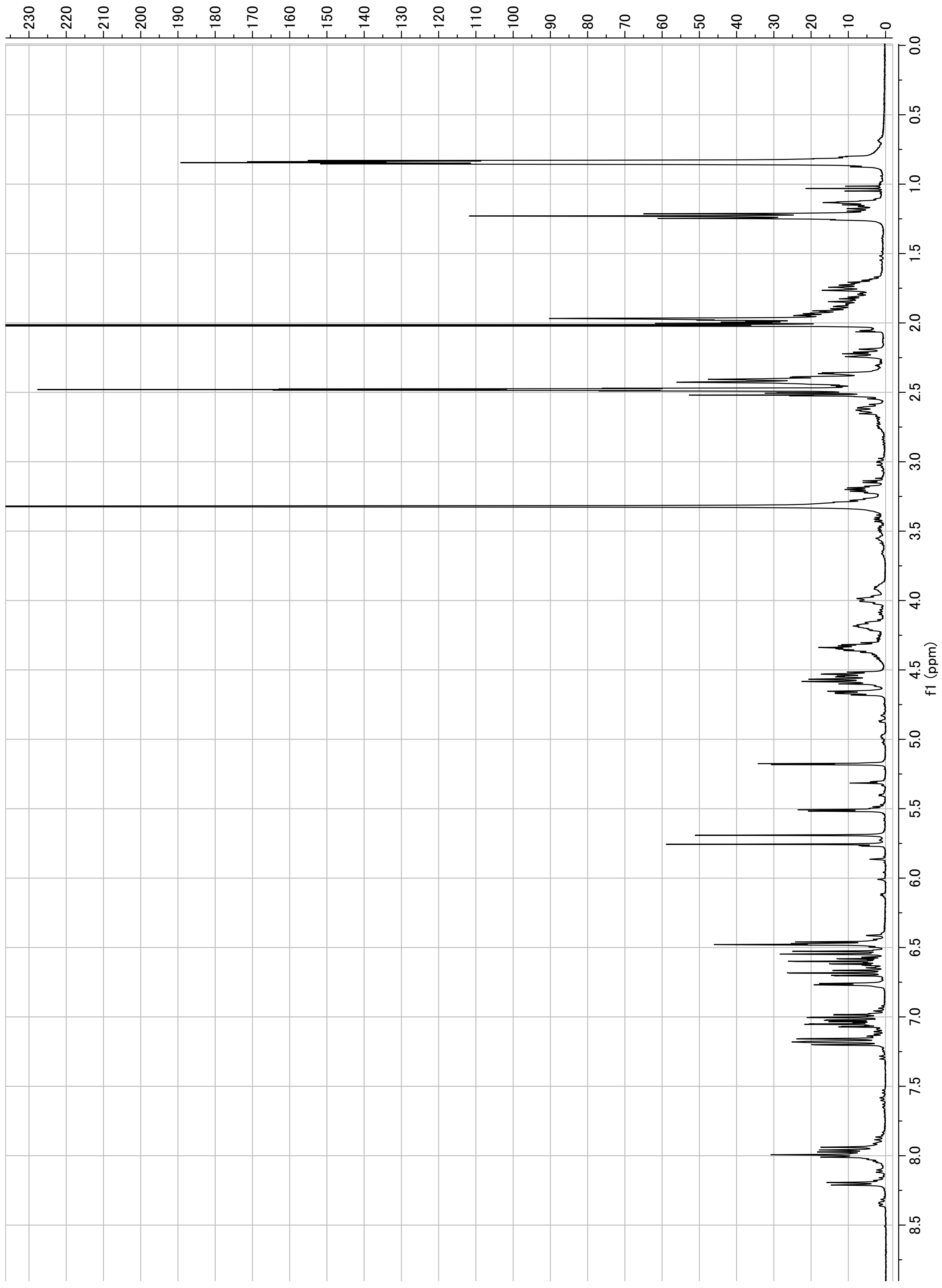

**Figure S14.** $^{1}$H NMR of lentindole B (**2**) (400 MHz, DMSO-$d_6$)

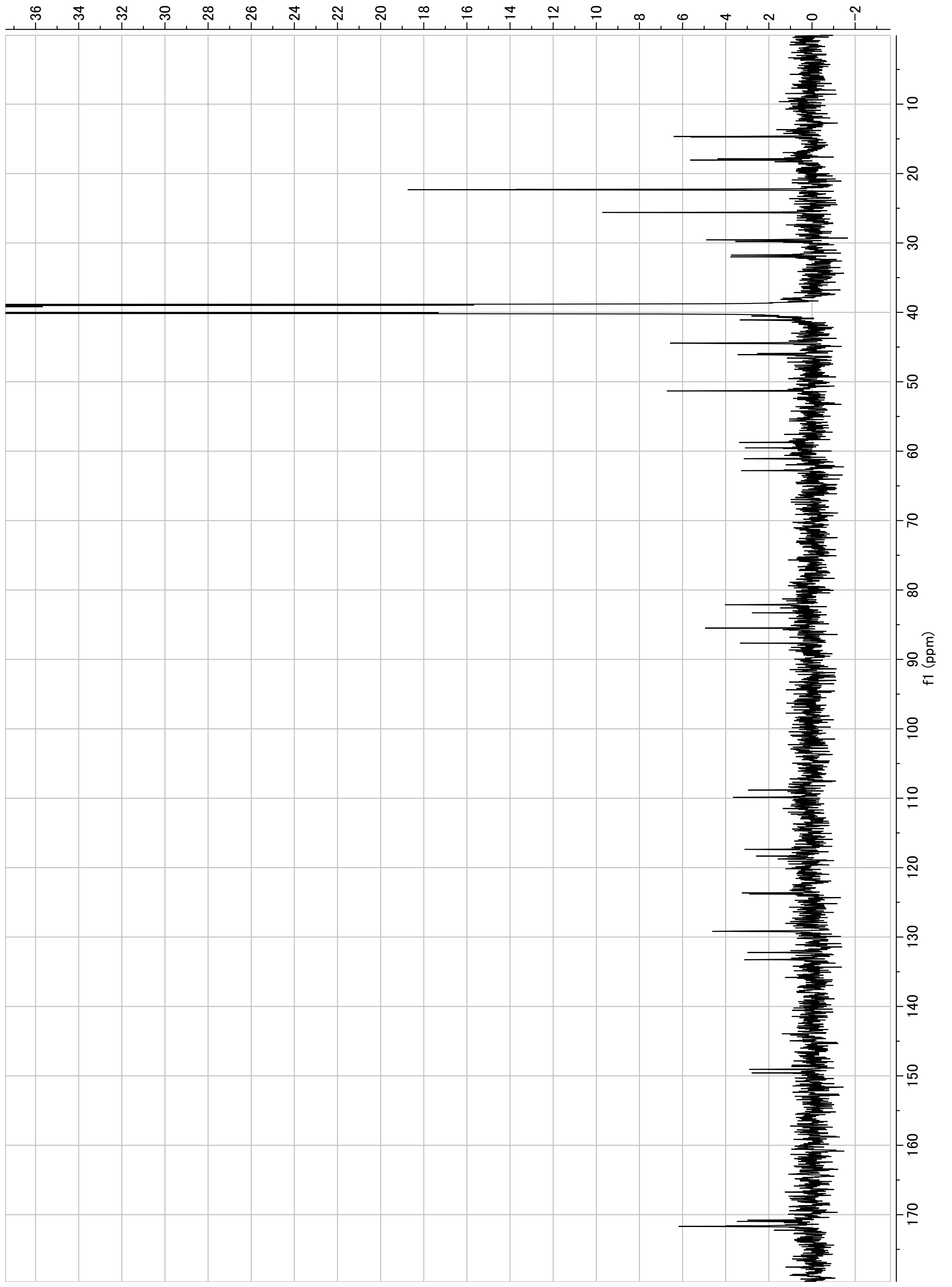

**Figure S15.** $^{13}C$ NMR of lentindole B (**2**) (100 MHz, DMSO-$d_6$)

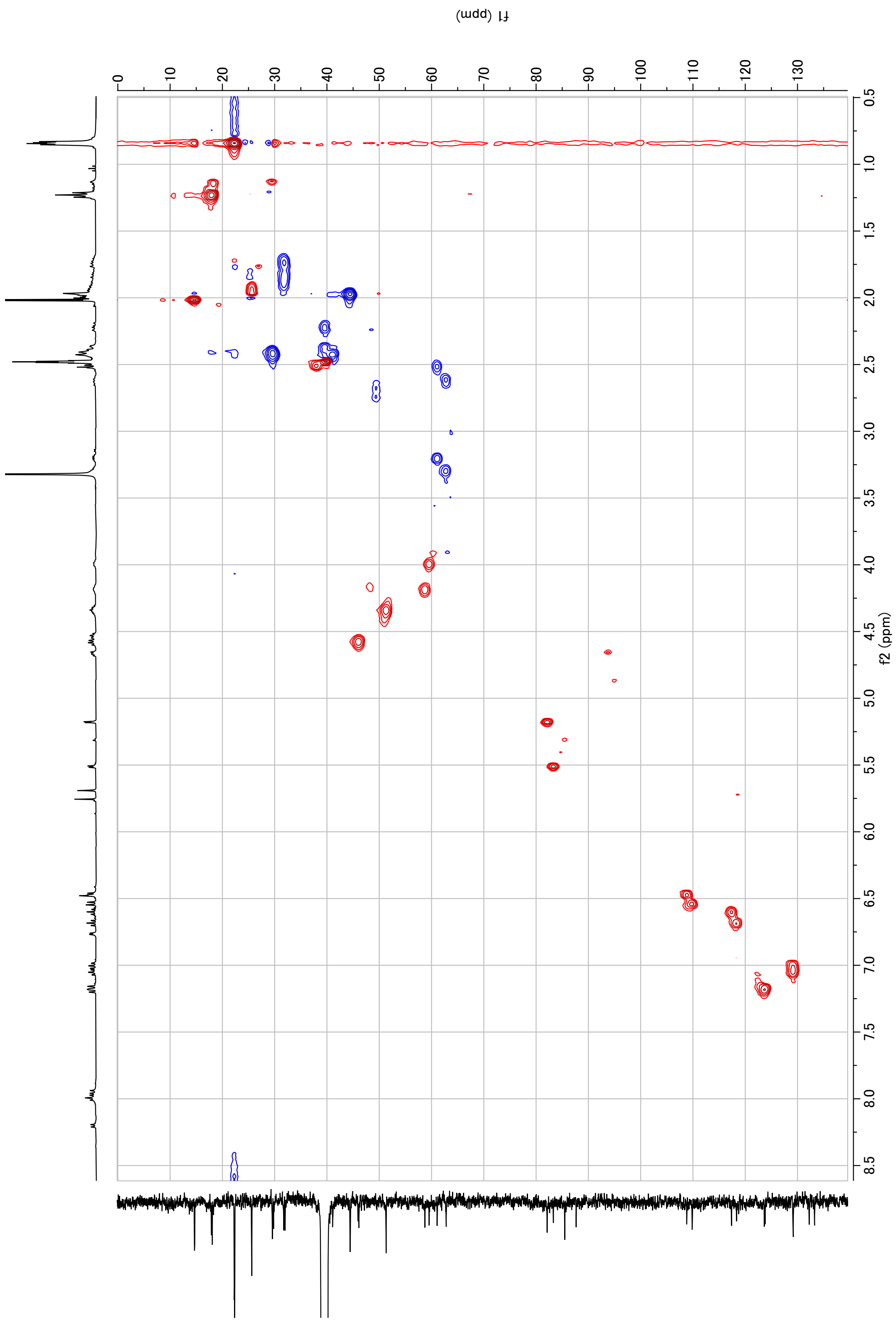

**Figure S16.** HSQC of lentindole B (**2**) ($^{1}$H:400 MHz, $^{13}$C100 MHz, DMSO-$d_6$)

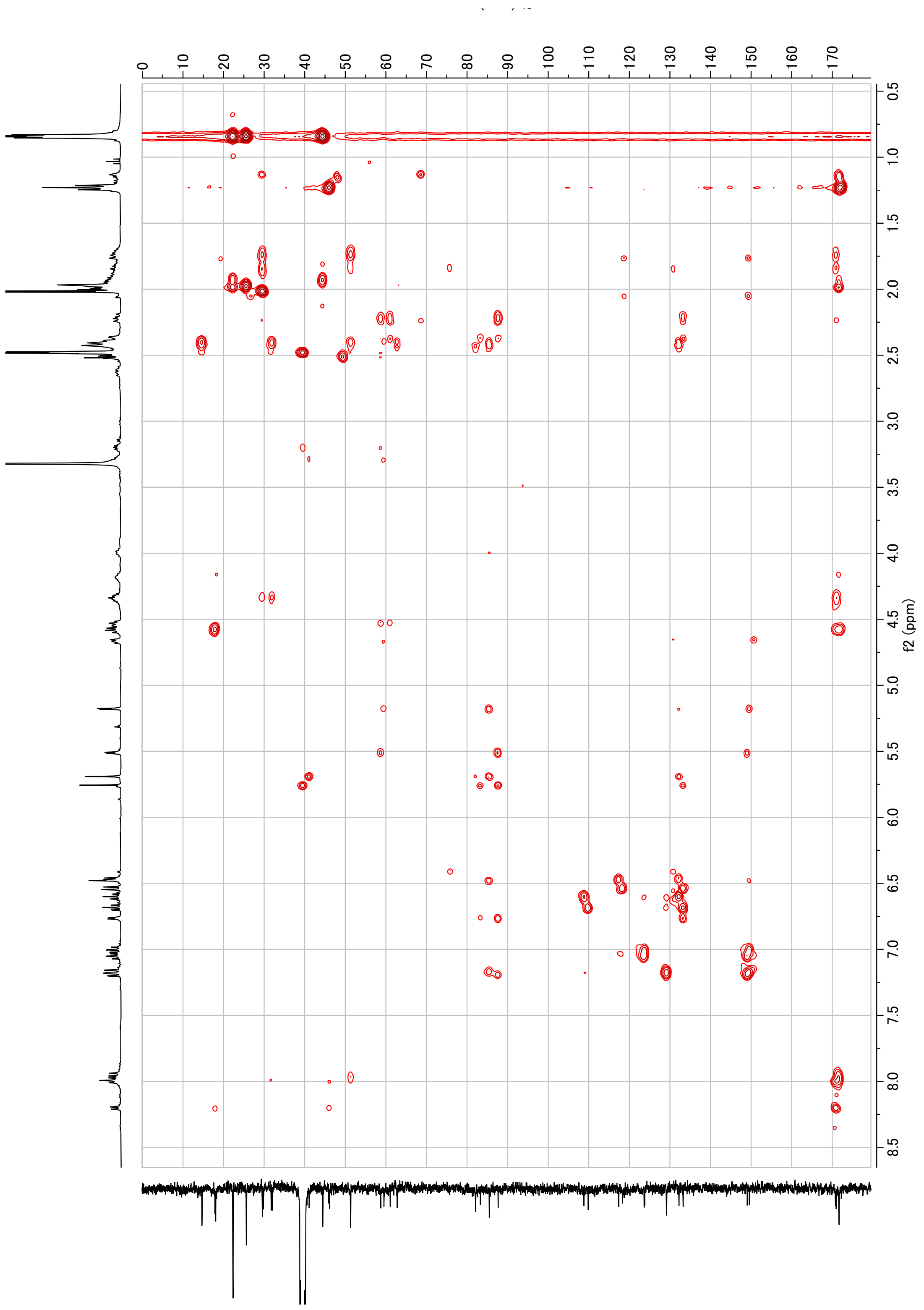

**Figure S17.** HMBC of lentindole B (**2**) ($^{1}$H:400 MHz, $^{13}$C100 MHz, DMSO-$d_6$)

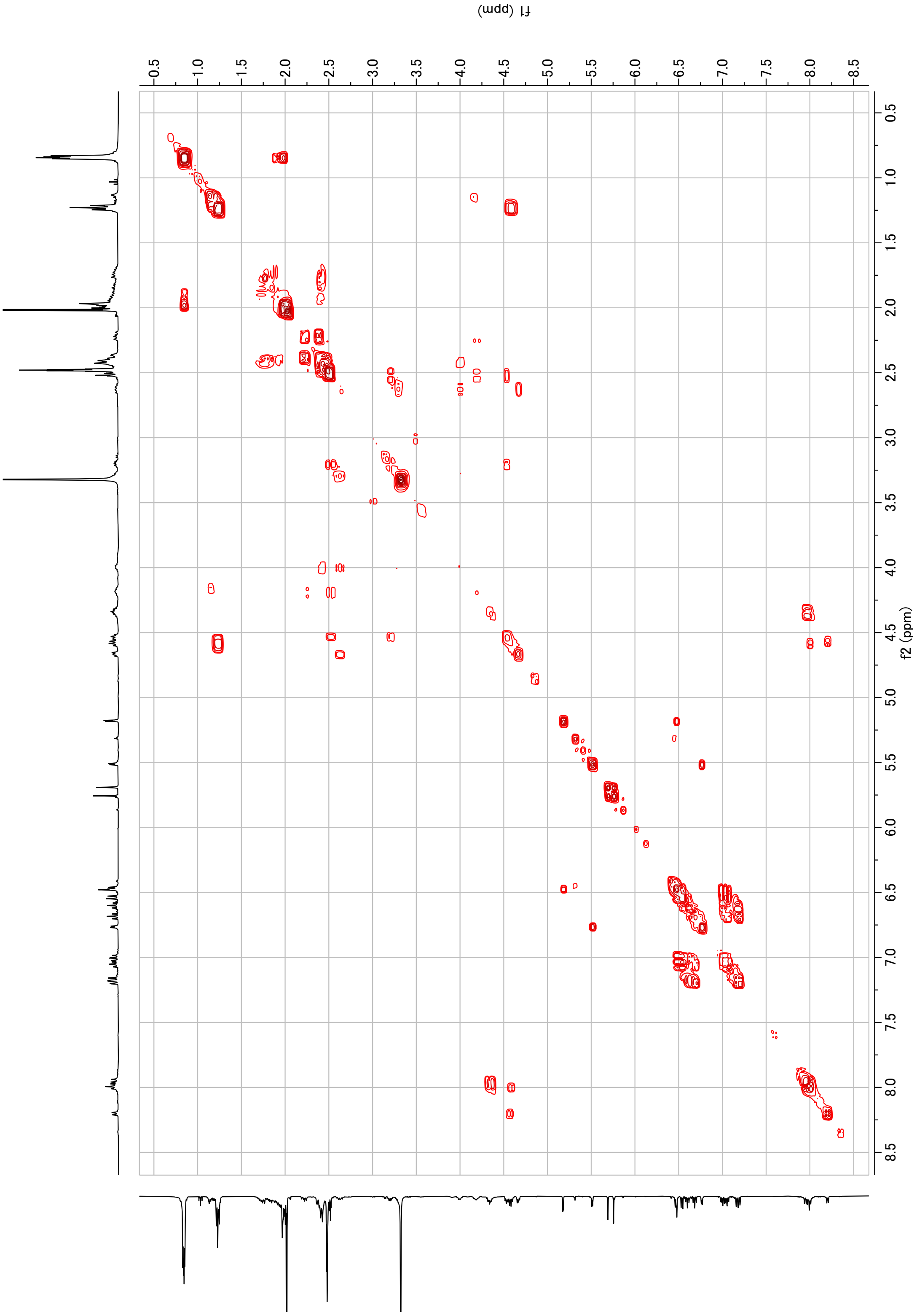

**Figure S18.** $^{1}H$-$^{1}H$ COSY of lentindole B (**2**) (400 MHz, DMSO-$d_6$)

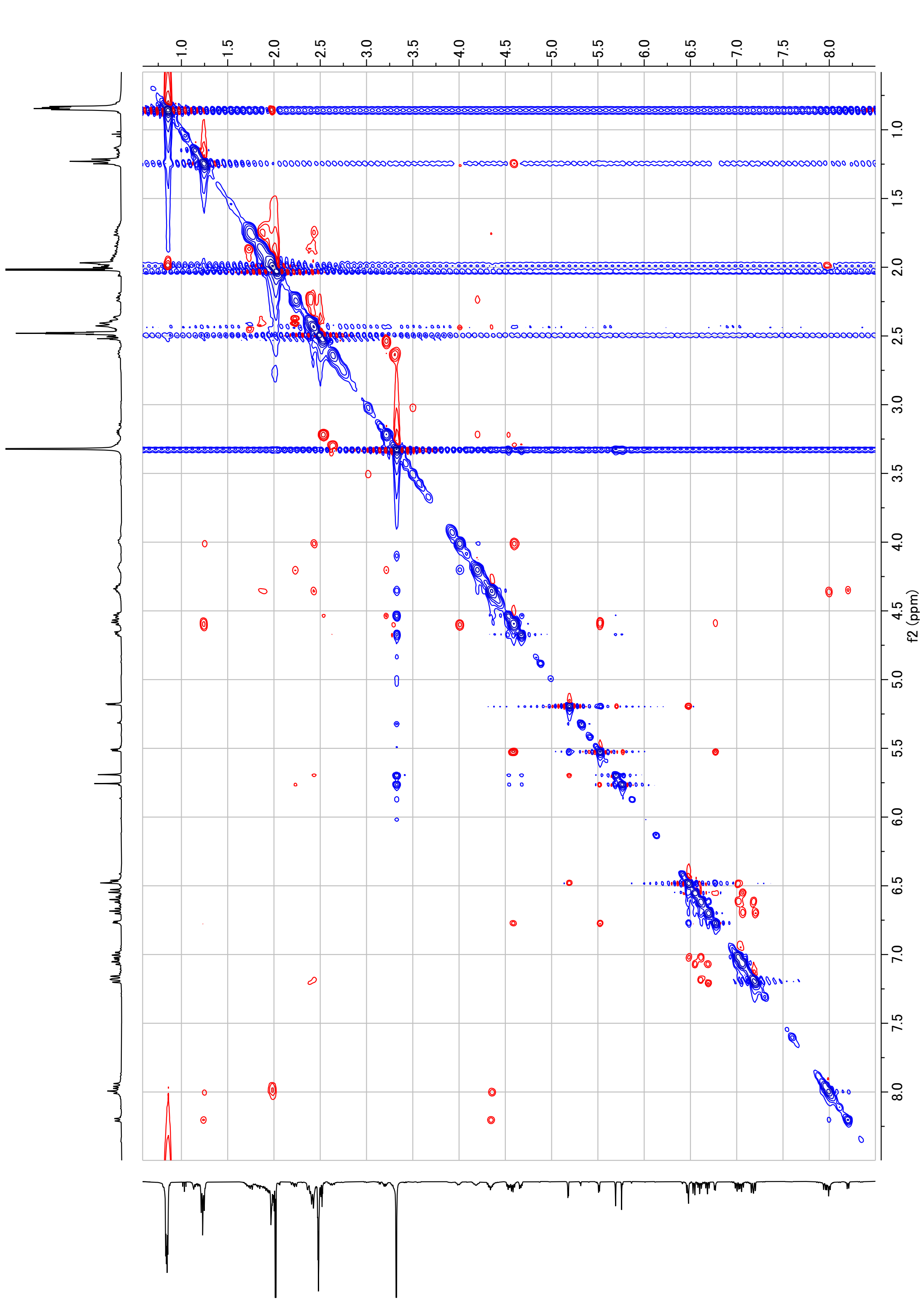

**Figure S19.** ROESY of lentindole B (**2**) (400 MHz, DMSO-$d_6$)

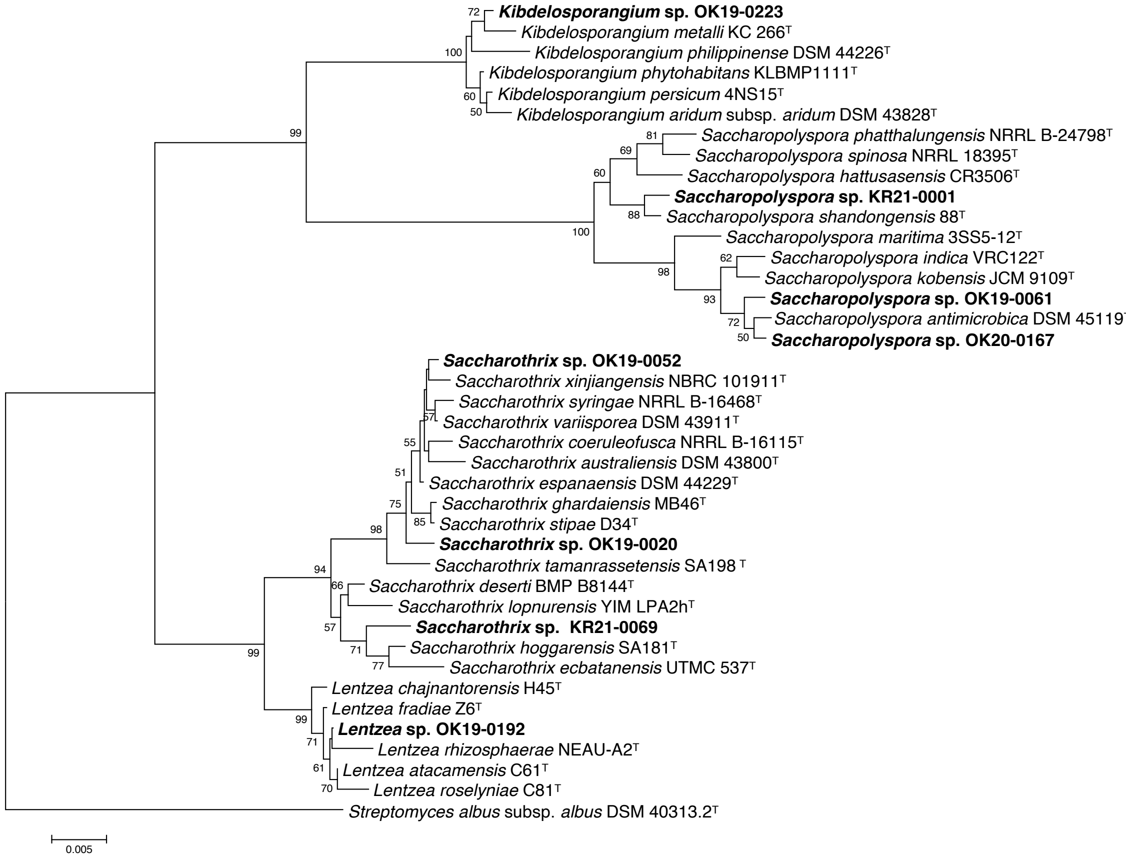

**Figure S20.** Phylogenetic tree among the strains used in this study and type strains of their closest species.

This tree was derived from 16S rRNA gene sequences, created using the neighbor-joining method. Only bootstrap values above 50 % (percentages of 1000 replications) are indicated. Bar, 0.005 substitutions per nucleotide position.

#### Supplementary Scheme

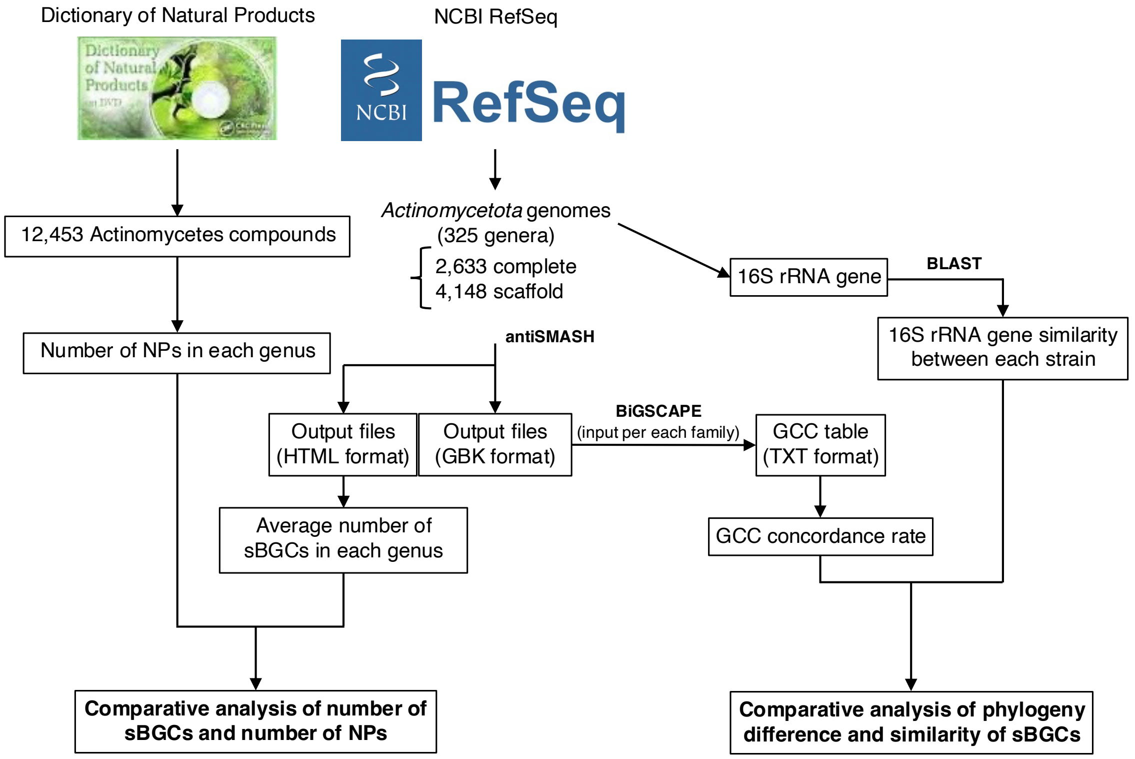

**Scheme S1**. The procedure for comparison of sBGC count and diversity.

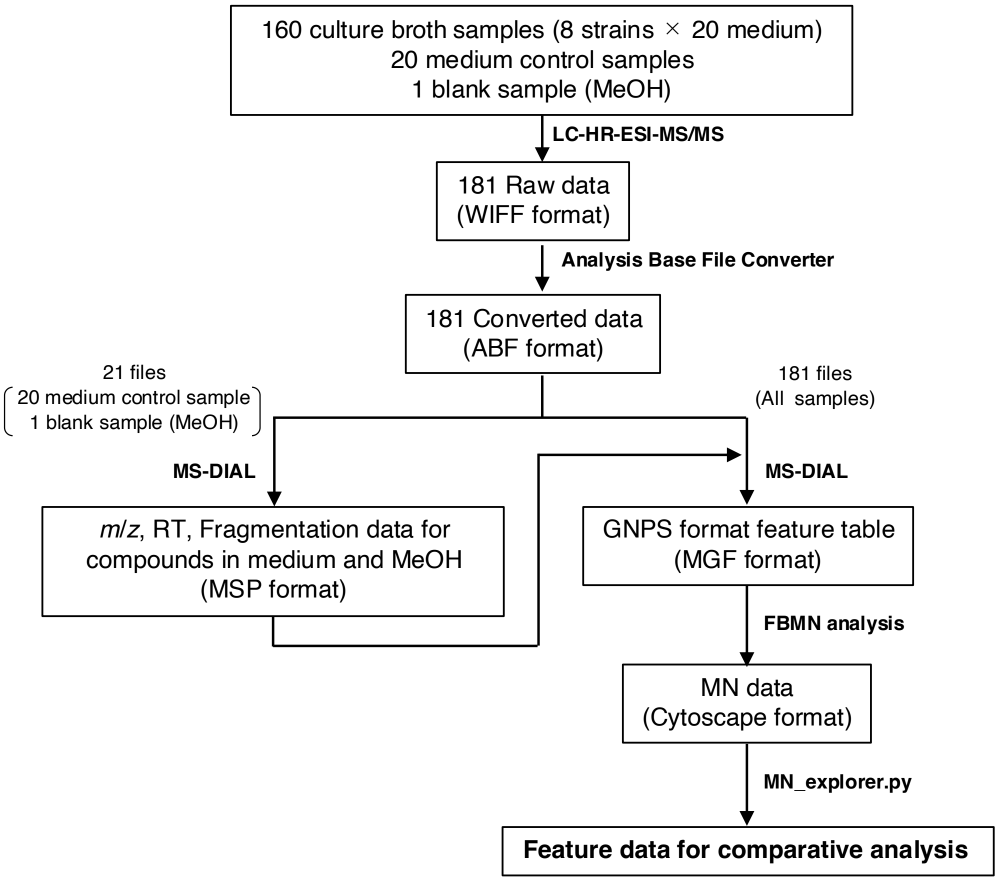

**Scheme S2**. The procedure for MN analysis

MN_explore.py is our original python script for comparison of MS intensity.

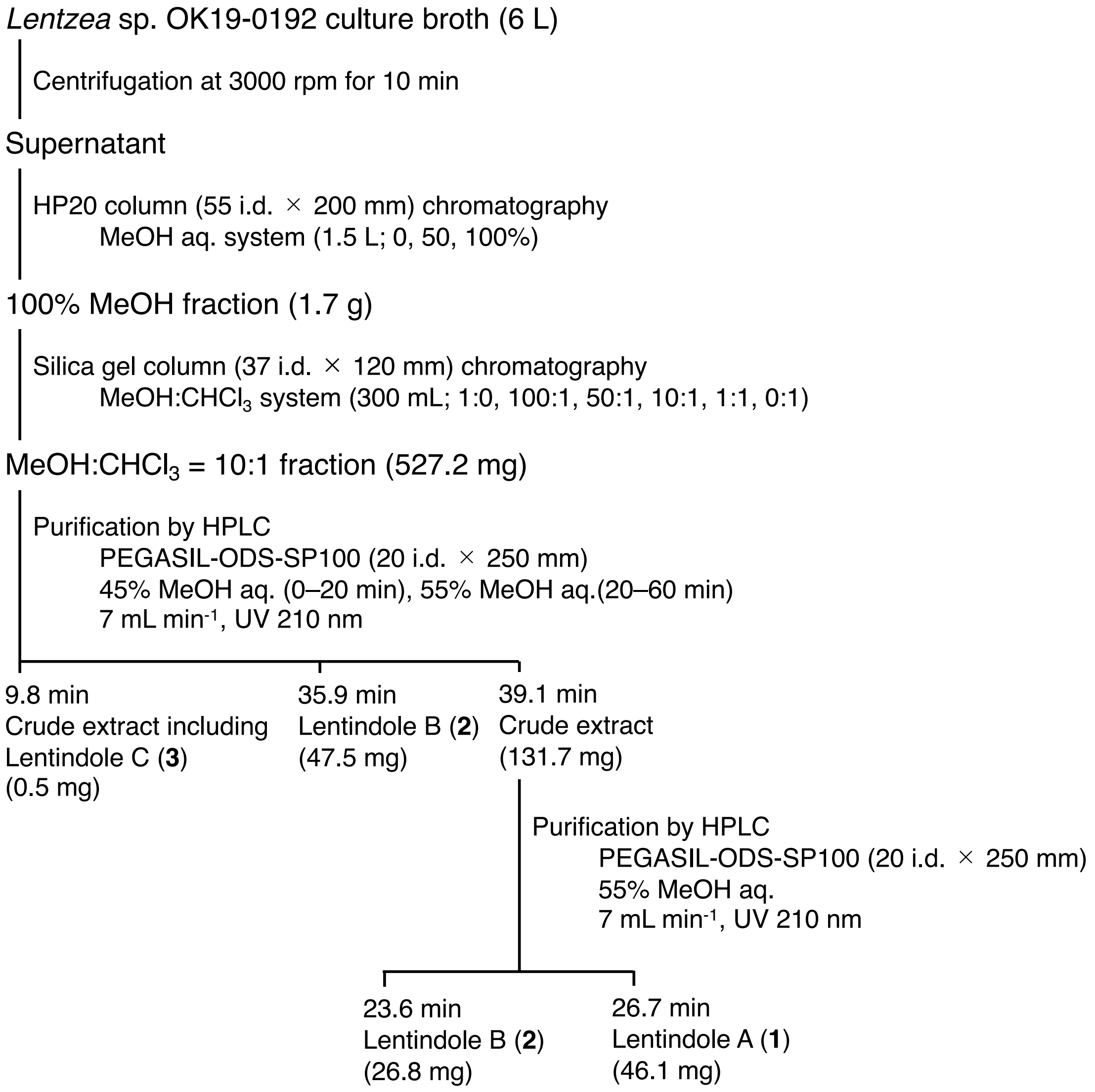

**Scheme S3**. The procedure for isolating lentindoles A–C (**1**–**3**)

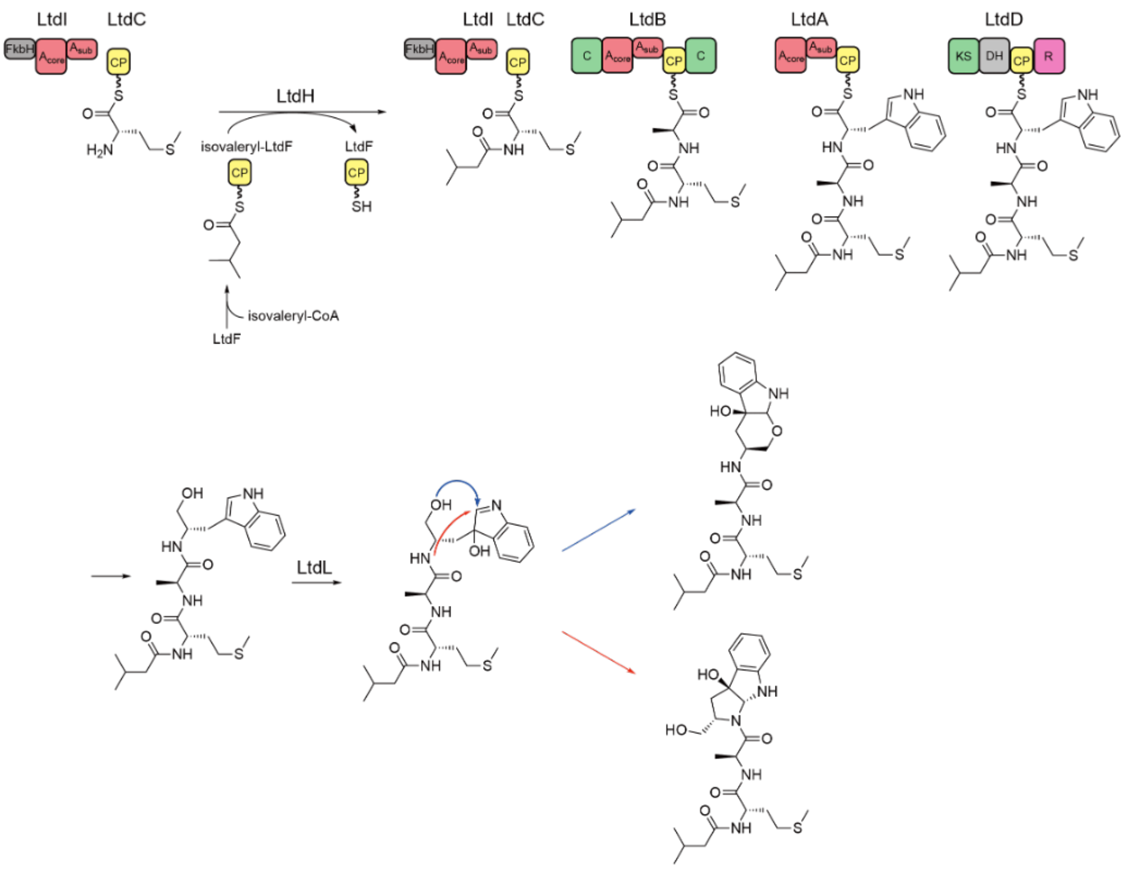

**Scheme S4**. Predicted biosynthetic pathway for lentindoles A (**1**) and B (**2**).

### Supplementary Table

**Table S1.** Biosynthetic types of sBGCs annotated by antiSMASH and their class in this study.

| Label | Product class |
|---|---|
| arylpolyene | PK |
| bottromycin | RiPP |
| cyanobactin | RiPP |
| epipeptide | RiPP |
| fungal-RiPP | RiPP |
| guanidinotides | RiPP |
| hglE-KS | PK |
| lantipeptide class I | RiPP |
| lantipeptide class II | RiPP |
| lantipeptide class III | RiPP |
| lantipeptide class IV | RiPP |
| lantipeptide class V | RiPP |
| LAP | RiPP |
| lassopeptide | RiPP |
| linaridin | RiPP |
| lipolanthine | RiPP |
| microviridin | RiPP |
| NAGGN | RiPP |
| nrps | NRP |
| nrps-like | NRP |
| PKS-like | PK |
| proteusin | RiPP |
| ranthipeptide | RiPP |
| RaS-RiPP | RiPP |
| RiPP-like | RiPP |
| RRE-containing | RiPP |
| saccharide | RiPP |
| sactipeptide | RiPP |
| spliceotide | RiPP |
| T1PKS | PK |
| T2PKS | PK |

| | |
|---|---|
| T3PKS | PK |
| terpene | Terpene |
| thioamide-NRP | RiPP |
| transAT-PKS | PK |
| transAT-PKS-like | PK |

**Table S2**. Average numbers of PNTR-BGCs and numbers of obtained NPs.

| Family | Genus | Ave. of PNTR-sBGC | NP |
|---|---|---|---|
| *Pseudonocardiaceae* | *Crossiella* | 48.3 | 0 |
| *Pseudonocardiaceae* | *Actinocrispum* | 46.0 | 0 |
| *Pseudonocardiaceae* | *Kibdelosporangium* | 42.3 | 33 |
| *Streptomycetaceae* | *Embleya* | 42.0 | 0 |
| *Pseudonocardiaceae* | *Allokutzneria* | 37.0 | 3 |
| *Pseudonocardiaceae* | *Kutzneria* | 36.0 | 9 |
| *Streptosporangiaceae* | *Thermocatellispora* | 34.0 | 0 |
| *Streptosporangiaceae* | *Bailinhaonella* | 34.0 | 0 |
| *Pseudonocardiaceae* | *Sciscionella* | 33.5 | 0 |
| *Pseudonocardiaceae* | *Saccharothrix* | 32.6 | 119 |
| *Streptomycetaceae* | *Kitasatospora* | 31.9 | 77 |
| *Pseudonocardiaceae* | *Longimycelium* | 31.0 | 0 |
| *Micromonosporaceae* | *Polymorphospora* | 30.0 | 3 |
| *Frankiaceae* | *Frankia* | 29.7 | 15 |
| *Streptosporangiaceae* | *Acrocarpospora* | 29.5 | 6 |
| *Pseudonocardiaceae* | *Actinokineospora* | 29.4 | 14 |
| *Streptosporangiaceae* | *Herbidospora* | 29.3 | 0 |
| *Pseudonocardiaceae* | *Lentzea* | 28.5 | 14 |
| *Pseudonocardiaceae* | *Labedaea* | 28.0 | 0 |
| *Streptosporangiaceae* | *Streptosporangium* | 27.6 | 74 |
| *Nocardiaceae* | *Nocardia* | 27.3 | 328 |
| *Streptosporangiaceae* | *Sinosporangium* | 27.0 | 0 |
| *Pseudonocardiaceae* | *Amycolatopsis* | 26.8 | 202 |
| *Pseudonocardiaceae* | *Umezawaea* | 26.5 | 2 |
| *Streptosporangiaceae* | *Microtetraspora* | 26.3 | 16 |
| *Streptosporangiaceae* | *Nonomuraea* | 26.3 | 57 |
| *Pseudonocardiaceae* | *Actinosynnema* | 25.5 | 62 |
| *Thermomonosporaceae* | *Actinomadura* | 25.2 | 289 |
| *Pseudonocardiaceae* | *Solihabitans* | 25.0 | 0 |
| *Streptosporangiaceae* | *Planobispora* | 25.0 | 1 |
| *Micromonosporaceae* | *Phytohabitans* | 24.5 | 1 |
| *Streptomycetaceae* | *Streptacidiphilus* | 24.2 | 6 |
| *Pseudonocardiaceae* | *Actinophytocola* | 24.0 | 0 |

| | | | |
|---|---|---|---|
| *Actinospicaceae* | *Actinospica* | 24.0 | 2 |
| *Streptosporangiaceae* | *Planomonospora* | 24.0 | 1 |
| *Streptomycetaceae* | *Streptomyces* | 23.3 | 9360 |
| *Pseudonocardiaceae* | *Saccharopolyspora* | 23.2 | 148 |
| *Pseudonocardiaceae* | *Alloactinosynnema* | 23.0 | 0 |
| *Streptosporangiaceae* | *Sphaerisporangium* | 22.5 | 9 |
| *Streptosporangiaceae* | *Microbispora* | 22.4 | 57 |
| *Micromonosporaceae* | *Mangrovihabitans* | 22.0 | 0 |
| *Micromonosporaceae* | *Micromonospora* | 21.8 | 529 |
| *Micromonosporaceae* | *Salinispora* | 21.6 | 134 |
| *Nocardiaceae* | *Antrihabitans* | 21.5 | 0 |
| *Pseudonocardiaceae* | *Actinoalloteichus* | 21.0 | 40 |
| *Catenulisporaceae* | *Catenulispora* | 21.0 | 4 |
| *Micromonosporaceae* | *Dactylosporangium* | 20.5 | 65 |
| *Micromonosporaceae* | *Verrucosispora* | 20.0 | 26 |
| *Micromonosporaceae* | *Plantactinospora* | 19.7 | 3 |
| *Streptomycetaceae* | *Yinghuangia* | 19.0 | 0 |
| *Pseudonocardiaceae* | *Herbihabitans* | 19.0 | 0 |
| *Micromonosporaceae* | *Longispora* | 18.0 | 1 |
| *Pseudonocardiaceae* | *Gandjariella* | 18.0 | 0 |
| *Nocardiopsaceae* | *Marinactinospora* | 18.0 | 13 |
| *Streptosporangiaceae* | *Thermoactinospora* | 18.0 | 0 |
| *Micromonosporaceae* | *Catellatospora* | 17.9 | 1 |
| *Actinopolysporaceae* | *Actinopolyspora* | 17.1 | 8 |
| *Micromonosporaceae* | *Hamadaea* | 17.0 | 0 |
| *Thermomonosporaceae* | *Actinocorallia* | 17.0 | 0 |
| *Micromonosporaceae* | *Pseudosporangium* | 16.5 | 0 |
| *Mycobacteriaceae* | *Mycobacteroides* | 16.5 | 0 |
| *Micromonosporaceae* | *Couchioplanes* | 16.0 | 0 |
| *Micromonosporaceae* | *Catelliglobosispora* | 16.0 | 0 |
| *Micromonosporaceae* | *Phytomonospora* | 16.0 | 0 |
| *Nocardiaceae* | *Rhodococcus* | 16.0 | 39 |
| *Nocardiopsaceae* | *Murinocardiopsis* | 15.0 | 0 |
| *Micromonosporaceae* | *Actinoplanes* | 14.4 | 143 |
| *Nocardiopsaceae* | *Nocardiopsis* | 14.3 | 190 |
| *Mycobacteriaceae* | *Mycobacterium* | 14.2 | 148 |

| | | | |
|---|---|---|---|
| *Micromonosporaceae* | *Natronosporangium* | 14.0 | 0 |
| *Micromonosporaceae* | *Asanoa* | 14.0 | 0 |
| *Gordoniaceae* | *Williamsia* | 13.5 | 1 |
| *Mycobacteriaceae* | *Mycolicibacterium* | 13.4 | 0 |
| *Thermomonosporaceae* | *Thermomonospora* | 13.2 | 4 |
| *Nocardiopsaceae* | *Marinitenerispora* | 12.5 | 0 |
| *Cryptosporangiaceae* | *Cryptosporangium* | 12.5 | 1 |
| *Tsukamurellaceae* | *Tsukamurella* | 12.3 | 28 |
| *Nocardiaceae* | *Prescottella* | 12.2 | 0 |
| *Nocardiaceae* | *Skermania* | 12.0 | 0 |
| *Nocardiopsaceae* | *Streptomonospora* | 12.0 | 4 |
| *Nocardiopsaceae* | *Actinorugispora* | 12.0 | 0 |
| *Micromonosporaceae* | *Actinocatenispora* | 11.7 | 0 |
| *Pseudonocardiaceae* | *Prauserella* | 11.4 | 0 |
| *Pseudonocardiaceae* | *Pseudonocardia* | 11.4 | 55 |
| *Glycomycetaceae* | *Stackebrandtia* | 11.0 | 0 |
| *Gordoniaceae* | *Gordonia* | 10.5 | 60 |
| *Nocardiopsaceae* | *Thermobifida* | 10.4 | 6 |
| *Mycobacteriaceae* | *Mycolicibacter* | 10.2 | 0 |
| *Nocardiaceae* | *Hoyosella* | 10.0 | 0 |
| *Pseudonocardiaceae* | *Saccharomonospora* | 9.7 | 16 |
| *Pseudonocardiaceae* | *Qaidamihabitans* | 9.0 | 0 |
| *Streptosporangiaceae* | *Thermopolyspora* | 9.0 | 0 |
| *Motilibacteraceae* | *Motilibacter* | 8.8 | 0 |
| *Pseudonocardiaceae* | *Haloechinothrix* | 8.5 | 0 |
| *Dermacoccaceae* | *Luteipulveratus* | 8.0 | 0 |
| *Glycomycetaceae* | *Natronoglycomyces* | 8.0 | 0 |
| *Streptomycetaceae* | *Carbonactinospora* | 8.0 | 0 |
| *Segniliparaceae* | *Segniliparus* | 7.5 | 0 |
| *Nocardiopsaceae* | *Spiractinospora* | 7.0 | 0 |
| *Treboniaceae* | *Trebonia* | 7.0 | 0 |
| *Vallicoccaceae* | *Vallicoccus* | 7.0 | 0 |
| *Kribbellaceae* | *Kribbella* | 6.7 | 7 |
| *Pseudonocardiaceae* | *Actinomycetospora* | 6.7 | 3 |
| *Geodermatophilaceae* | *Geodermatophilus* | 6.4 | 1 |
| *Jiangellaceae* | *Jiangella* | 6.4 | 7 |

| | | | |
|---|---|---|---|
| *Promicromonosporaceae* | *Myceligenerans* | 6.3 | 0 |
| *Mycobacteriaceae* | *Mycolicibacillus* | 6.0 | 0 |
| *Pseudonocardiaceae* | *Allosaccharopolyspora* | 6.0 | 0 |
| *Egibacteraceae* | *Egibacter* | 6.0 | 0 |
| *Iamiaceae* | *Aquihabitans* | 6.0 | 0 |
| *Nocardiaceae* | *Tomitella* | 6.0 | 0 |
| *Actinopolymorphaceae* | *Actinopolymorpha* | 6.0 | 3 |
| *Kineosporiaceae* | *Kineococcus* | 6.0 | 0 |
| *Microbacteriaceae* | *Clavibacter* | 5.8 | 1 |
| *Rubrobacteraceae* | *Rubrobacter* | 5.8 | 2 |
| *Microbacteriaceae* | *Curtobacterium* | 5.7 | 3 |
| *Glycomycetaceae* | *Glycomyces* | 5.7 | 2 |
| *Sporichthyaceae* | *Sporichthya* | 5.5 | 0 |
| *Microbacteriaceae* | *Rathayibacter* | 5.3 | 0 |
| *Microbacteriaceae* | *Plantibacter* | 5.3 | 0 |
| *Jiangellaceae* | *Phytoactinopolyspora* | 5.2 | 0 |
| *Baekduiaceae* | *Baekduia* | 5.0 | 0 |
| *Iamiaceae* | *Actinomarinicola* | 5.0 | 0 |
| *Ilumatobacteraceae* | *Ilumatobacter* | 5.0 | 0 |
| *Kineosporiaceae* | *Kineosporia* | 5.0 | 0 |
| *Micrococcaceae* | *Psychromicrobium* | 5.0 | 0 |
| *Geodermatophilaceae* | *Modestobacter* | 5.0 | 0 |
| *Kineosporiaceae* | *Quadrisphaera* | 5.0 | 0 |
| *Propionibacteriaceae* | *Microlunatus* | 5.0 | 1 |
| *Kineosporiaceae* | *Angustibacter* | 5.0 | 0 |
| *Solirubrobacteraceae* | *Solirubrobacter* | 4.7 | 0 |
| *Promicromonosporaceae* | *Cellulosimicrobium* | 4.6 | 0 |
| *Micrococcaceae* | *Auritidibacter* | 4.5 | 0 |
| *Dermatophilaceae* | *Austwickia* | 4.5 | 0 |
| *Actinopolysporaceae* | *Halopolyspora* | 4.0 | 0 |
| *Dermacoccaceae* | *Flexivirga* | 4.0 | 0 |
| *Micrococcaceae* | *Neomicrococcus* | 4.0 | 0 |
| *Geodermatophilaceae* | *Cumulibacter* | 4.0 | 0 |
| *Microbacteriaceae* | *Glaciibacter* | 4.0 | 0 |
| *Patulibacteraceae* | *Patulibacter* | 4.0 | 0 |
| *Dermacoccaceae* | *Yimella* | 4.0 | 0 |

| | | | |
|---|---|---|---|
| *Ruaniaceae* | *Occultella* | 4.0 | 0 |
| *Euzebyaceae* | *Euzebya* | 4.0 | 0 |
| *Jatrophihabitantaceae* | *Jatrophihabitans* | 4.0 | 0 |
| *Jiangellaceae* | *Haloactinopolyspora* | 4.0 | 0 |
| *Microbacteriaceae* | *Diaminobutyricibacter* | 4.0 | 0 |
| *Microbacteriaceae* | *Glaciihabitans* | 4.0 | 0 |
| *Micrococcaceae* | *Renibacterium* | 4.0 | 2 |
| *Propionibacteriaceae* | *Micropruina* | 4.0 | 0 |
| *Intrasporangiaceae* | *Tetrasphaera* | 4.0 | 0 |
| *Miltoncostaeaceae* | *Miltoncostaea* | 4.0 | 0 |
| *Dermabacteraceae* | *Devriesea* | 4.0 | 0 |
| *Microbacteriaceae* | *Marisediminicola* | 4.0 | 0 |
| *Propionibacteriaceae* | *Arachnia* | 4.0 | 0 |
| *Promicromonosporaceae* | *Oerskovia* | 3.7 | 0 |
| *Microbacteriaceae* | *Agreia* | 3.5 | 0 |
| *Nakamurellaceae* | *Nakamurella* | 3.5 | 0 |
| *Geodermatophilaceae* | *Klenkia* | 3.5 | 0 |
| *Microbacteriaceae* | *Frondihabitans* | 3.3 | 0 |
| *Promicromonosporaceae* | *Promicromonospora* | 3.3 | 6 |
| *Dietziaceae* | *Dietzia* | 3.3 | 4 |
| *Dermatophilaceae* | *Dermatophilus* | 3.3 | 0 |
| *Microbacteriaceae* | *Herbiconiux* | 3.3 | 0 |
| *Bogoriellaceae* | *Georgenia* | 3.2 | 0 |
| *Micrococcaceae* | *Paenarthrobacter* | 3.2 | 0 |
| *Microbacteriaceae* | *Cryobacterium* | 3.1 | 0 |
| *Cellulomonadaceae* | *Cellulomonas* | 3.1 | 2 |
| *Corynebacteriaceae* | *Corynebacterium* | 3.0 | 89 |
| *Micrococcaceae* | *Zhihengliuella* | 3.0 | 0 |
| *Microbacteriaceae* | *Planctomonas* | 3.0 | 0 |
| *Microbacteriaceae* | *Schumannella* | 3.0 | 0 |
| *Nocardioidaceae* | *Pimelobacter* | 3.0 | 0 |
| *Micrococcaceae* | *Sinomonas* | 3.0 | 0 |
| *Microbacteriaceae* | *Subtercola* | 3.0 | 0 |
| *Dermatophilaceae* | *Gephyromycinifex* | 3.0 | 0 |
| *Microbacteriaceae* | *Cnuibacter* | 3.0 | 0 |
| *Microbacteriaceae* | *Conyzicola* | 3.0 | 0 |

| | | | |
|---|---|---|---|
| *Microbacteriaceae* | *Diaminobutyricimonas* | 3.0 | 0 |
| *Microbacteriaceae* | *Gryllotalpicola* | 3.0 | 0 |
| *Microbacteriaceae* | *Humibacter* | 3.0 | 0 |
| *Promicromonosporaceae* | *Krasilnikoviella* | 3.0 | 0 |
| *Propionibacteriaceae* | *Granulicoccus* | 3.0 | 0 |
| *Sporichthyaceae* | *Epidermidibacterium* | 3.0 | 0 |
| *Streptosporangiaceae* | *Thermobispora* | 3.0 | 0 |
| *Geodermatophilaceae* | *Blastococcus* | 3.0 | 0 |
| *Cellulomonadaceae* | *Pseudactinotalea* | 3.0 | 0 |
| *Dermacoccaceae* | *Allobranchiibius* | 3.0 | 0 |
| *Egicoccaceae* | *Egicoccus* | 3.0 | 0 |
| *Promicromonosporaceae* | *Luteimicrobium* | 3.0 | 0 |
| *Dermatophilaceae* | *Mobilicoccus* | 3.0 | 0 |
| *Eggerthellaceae* | *Denitrobacterium* | 3.0 | 0 |
| *Intrasporangiaceae* | *Intrasporangium* | 3.0 | 0 |
| *Kineosporiaceae* | *Pseudokineococcus* | 3.0 | 0 |
| *Promicromonosporaceae* | *Puerhibacterium* | 3.0 | 0 |
| *Propionibacteriaceae* | *Aestuariimicrobium* | 3.0 | 0 |
| *Propionibacteriaceae* | *Auraticoccus* | 3.0 | 0 |
| *Microbacteriaceae* | *Leifsonia* | 2.9 | 0 |
| *Microbacteriaceae* | *Mycetocola* | 2.8 | 0 |
| *Brevibacteriaceae* | *Brevibacterium* | 2.8 | 37 |
| *Microbacteriaceae* | *Pseudoclavibacter* | 2.8 | 0 |
| *Microbacteriaceae* | *Salinibacterium* | 2.8 | 0 |
| *Eggerthellaceae* | *Raoultibacter* | 2.8 | 0 |
| *Microbacteriaceae* | *Frigoribacterium* | 2.7 | 0 |
| *Eggerthellaceae* | *Eggerthella* | 2.7 | 0 |
| *Micrococcaceae* | *Citricoccus* | 2.7 | 1 |
| *Micrococcaceae* | *Arthrobacter* | 2.6 | 61 |
| *Microbacteriaceae* | *Agrococcus* | 2.6 | 0 |
| *Micrococcaceae* | *Kocuria* | 2.6 | 2 |
| *Ruaniaceae* | *Ruania* | 2.5 | 0 |
| *Cellulomonadaceae* | *Actinotalea* | 2.5 | 0 |
| *Microbacteriaceae* | *Naasia* | 2.5 | 0 |
| *Ornithinimicrobiaceae* | *Ornithinimicrobium* | 2.5 | 1 |
| *Propionibacteriaceae* | *Propioniciclava* | 2.5 | 0 |

| | | | |
|---|---|---|---|
| *Microbacteriaceae* | *Microbacterium* | 2.5 | 9 |
| *Microbacteriaceae* | *Agromyces* | 2.4 | 3 |
| *Eggerthellaceae* | *Gordonibacter* | 2.4 | 0 |
| *Micrococcaceae* | *Glutamicibacter* | 2.4 | 0 |
| *Micrococcaceae* | *Pseudarthrobacter* | 2.4 | 0 |
| *Promicromonosporaceae* | *Isoptericola* | 2.4 | 1 |
| *Dermacoccaceae* | *Dermacoccus* | 2.4 | 11 |
| *Micrococcaceae* | *Nesterenkonia* | 2.3 | 4 |
| *Dermabacteraceae* | *Brachybacterium* | 2.3 | 0 |
| *Dermatophilaceae* | *Arsenicicoccus* | 2.3 | 0 |
| *Demequinaceae* | *Demequina* | 2.2 | 0 |
| *Microbacteriaceae* | *Microcella* | 2.2 | 0 |
| *Intrasporangiaceae* | *Terrabacter* | 2.2 | 0 |
| *Nocardioidaceae* | *Nocardioides* | 2.2 | 11 |
| *Micrococcaceae* | *Micrococcus* | 2.1 | 64 |
| *Actinomycetaceae* | *Actinomyces* | 2.1 | 101 |
| *Propionibacteriaceae* | *Cutibacterium* | 2.0 | 0 |
| *Micrococcaceae* | *Paeniglutamicibacter* | 2.0 | 0 |
| *Dermacoccaceae* | *Leekyejoonella* | 2.0 | 0 |
| *Micrococcaceae* | *Zafaria* | 2.0 | 0 |
| *Acidothermaceae* | *Acidothermus* | 2.0 | 3 |
| *Microbacteriaceae* | *Chryseoglobus* | 2.0 | 0 |
| *Microbacteriaceae* | *Marinisubtilis* | 2.0 | 0 |
| *Propionibacteriaceae* | *Enemella* | 2.0 | 0 |
| *Thermoleophilaceae* | *Thermoleophilum* | 2.0 | 9 |
| *Beutenbergiaceae* | *Miniimonas* | 2.0 | 0 |
| *Dermacoccaceae* | *Calidifontibacter* | 2.0 | 0 |
| *Dermacoccaceae* | *Metallococcus* | 2.0 | 0 |
| *Iamiaceae* | *Iamia* | 2.0 | 0 |
| *Intrasporangiaceae* | *Terracoccus* | 2.0 | 0 |
| *Microbacteriaceae* | *Homoserinibacter* | 2.0 | 0 |
| *Microbacteriaceae* | *Microterricola* | 2.0 | 0 |
| *Microbacteriaceae* | *Yonghaparkia* | 2.0 | 0 |
| *Promicromonosporaceae* | *Paraoerskovia* | 2.0 | 0 |
| *Propionibacteriaceae* | *Raineyella* | 2.0 | 0 |
| *Intrasporangiaceae* | *Phycicoccus* | 2.0 | 0 |

| | | | |
|---|---|---|---|
| *Acidimicrobiaceae* | *Ferrimicrobium* | 2.0 | 0 |
| *Acidimicrobiaceae* | *Ferrithrix* | 2.0 | 0 |
| *Bifidobacteriaceae* | *Parascardovia* | 2.0 | 0 |
| *Intrasporangiaceae* | *Knoellia* | 2.0 | 0 |
| *Jonesiaceae* | *Flavimobilis* | 2.0 | 0 |
| *Microbacteriaceae* | *Pontimonas* | 2.0 | 0 |
| *Microbacteriaceae* | *Leucobacter* | 2.0 | 0 |
| *Propionibacteriaceae* | *Acidipropionibacterium* | 1.9 | 0 |
| *Kytococcaceae* | *Kytococcus* | 1.8 | 0 |
| *Micrococcaceae* | *Rothia* | 1.8 | 3 |
| *Propionibacteriaceae* | *Tessaracoccus* | 1.7 | 0 |
| *Jonesiaceae* | *Sanguibacter* | 1.7 | 0 |
| *Actinomycetaceae* | *Actinobaculum* | 1.6 | 0 |
| *Nocardioidaceae* | *Mumia* | 1.5 | 0 |
| *Ornithinimicrobiaceae* | *Serinicoccus* | 1.5 | 3 |
| *Conexibacteraceae* | *Conexibacter* | 1.4 | 0 |
| *Actinomycetaceae* | *Flaviflexus* | 1.4 | 0 |
| *Microbacteriaceae* | *Aurantimicrobium* | 1.3 | 0 |
| *Microbacteriaceae* | *Protaetiibacter* | 1.3 | 0 |
| *Intrasporangiaceae* | *Janibacter* | 1.3 | 2 |
| *Nocardioidaceae* | *Aeromicrobium* | 1.3 | 0 |
| *Dermabacteraceae* | *Dermabacter* | 1.3 | 0 |
| *Nocardioidaceae* | *Marmoricola* | 1.3 | 0 |
| *Promicromonosporaceae* | *Xylanimonas* | 1.3 | 0 |
| *Microbacteriaceae* | *Gulosibacter* | 1.2 | 0 |
| *Jonesiaceae* | *Jonesia* | 1.2 | 0 |
| *Nitriliruptoraceae* | *Nitriliruptor* | 1.0 | 0 |
| *Pseudonocardiaceae* | *Thermocrispum* | 1.0 | 0 |
| *Microbacteriaceae* | *Lacisediminihabitans* | 1.0 | 0 |
| *Acidimicrobiaceae* | *Acidimicrobium* | 1.0 | 0 |
| *Microbacteriaceae* | *Rhodoluna* | 1.0 | 0 |
| *Micrococcaceae* | *Specibacter* | 1.0 | 0 |
| *Salsipaludibacteraceae* | *Salsipaludibacter* | 1.0 | 0 |
| *Actinomycetaceae* | *Actinotignum* | 1.0 | 0 |
| *Actinomycetaceae* | *Buchananella* | 1.0 | 0 |
| *Beutenbergiaceae* | *Beutenbergia* | 1.0 | 0 |

| | | | |
|---|---|---|---|
| *Intrasporangiaceae* | *Pedococcus* | 1.0 | 0 |
| *Lawsonellaceae* | *Lawsonella* | 1.0 | 0 |
| *Propionibacteriaceae* | *Propionicicella* | 1.0 | 0 |
| *Bifidobacteriaceae* | *Alloscardovia* | 0.8 | 0 |
| *Propionibacteriaceae* | *Propionibacterium* | 0.7 | 37 |
| *Eggerthellaceae* | *Adlercreutzia* | 0.7 | 0 |
| *Bifidobacteriaceae* | *Gardnerella* | 0.7 | 1 |
| *Actinomycetaceae* | *Gleimia* | 0.7 | 0 |
| *Bifidobacteriaceae* | *Scardovia* | 0.7 | 0 |
| *Actinomycetaceae* | *Arcanobacterium* | 0.6 | 0 |
| *Eggerthellaceae* | *Slackia* | 0.6 | 1 |
| *Actinomycetaceae* | *Bowdeniella* | 0.5 | 0 |
| *Atopobiaceae* | *Thermophilibacter* | 0.5 | 0 |
| *Bifidobacteriaceae* | *Bifidobacterium* | 0.4 | 14 |
| *Actinomycetaceae* | *Boudabousia* | 0.3 | 0 |
| *Atopobiaceae* | *Olsenella* | 0.2 | 0 |
| *Actinomycetaceae* | *Schaalia* | 0.2 | 0 |
| *Actinomycetaceae* | *Mobiluncus* | 0.1 | 0 |
| *Coriobacteriaceae* | *Collinsella* | 0.1 | 0 |
| *Actinomycetaceae* | *Trueperella* | 0.0 | 0 |
| *Eggerthellaceae* | *Berryella* | 0.0 | 0 |
| *Eggerthellaceae* | *Hugonella* | 0.0 | 0 |
| *Eggerthellaceae* | *Senegalimassilia* | 0.0 | 0 |
| *Micrococcaceae* | *Pseudoglutamicibacter* | 0.0 | 0 |
| *Actinomycetaceae* | *Nanchangia* | 0.0 | 0 |
| *Actinomycetaceae* | *Varibaculum* | 0.0 | 0 |
| *Actinomycetaceae* | *Winkia* | 0.0 | 0 |
| *Atopobiaceae* | *Atopobium* | 0.0 | 0 |
| *Atopobiaceae* | *Fannyhessea* | 0.0 | 0 |
| *Atopobiaceae* | *Lancefieldella* | 0.0 | 0 |
| *Atopobiaceae* | *Parafannyhessea* | 0.0 | 0 |
| *Atopobiaceae* | *Parolsenella* | 0.0 | 0 |
| *Coriobacteriaceae* | *Coriobacterium* | 0.0 | 0 |
| *Coriobacteriaceae* | *Enorma* | 0.0 | 0 |
| *Eggerthellaceae* | *Cryptobacterium* | 0.0 | 0 |
| *Eggerthellaceae* | *Phoenicibacter* | 0.0 | 0 |

| | | | |
|---|---|---|---|
| *Jonesiaceae* | *Timonella* | 0.0 | 0 |
| *Propionibacteriaceae* | *Propionimicrobium* | 0.0 | 0 |
| *Propionibacteriaceae* | *Vaginimicrobium* | 0.0 | 0 |
| *Tropherymataceae* | *Tropheryma* | 0.0 | 0 |

**Table S3.** *Pseudonocardiaceae* strains used in this study

| Strain | Isolation source |
|---|---|
| *Saccharothrix* sp. OK19-0052 | Soil under a pine tree on Minamidaito Island, Okinawa, Japan. |
| *Saccharothrix* sp. OK19-0020 | Soil under an *Asplenium antiquum* Makino on Minamidaito Island, Okinawa, Japan. |
| *Saccharothrix* sp. KR21-0069 | Soil under a *Ficus microcarpa* on Noho Island, Okinawa, Japan. |
| *Saccharopolyspora* sp. OK20-0167 | Soil under a pine tree on Noho Island, Okinawa, Japan. |
| *Saccharopolyspora* sp. OK19-0061 | Soil under a pine tree on Minamidaito Island |
| *Saccharopolyspora* sp. KR21-0001 | Soil under a pine tree on Oha Island, Okinawa, Japan. |
| *Lentzea* sp. OK19-0192 | Soil under a *Diospyros ferrea* Bakhauizen var. buxifolia Bakh on Tonaki Island, Okinawa, Japan. |
| *Kibdelosporangium* sp. OK19-0223 | Soil under a *Diospyros ferrea* Bakhauizen var. buxifolia Bakh on Tonaki Island, Okinawa, Japan. |

**Table S4.** Medium conditions

| Name | Component | Concentration |
|---|---|---|
| Medium 1 | Glucose | 1.00% (w/v) |
| | Starch | 2.00% (w/v) |
| | Yeast extract | 0.50% (w/v) |
| | Peptone | 0.50% (w/v) |
| | $CaCO_3$ | 0.40% (w/v) |
| Medium 2 | Mannitol | 3.00% (w/v) |
| | Glucose | 1.00% (w/v) |
| | Yesat extract | 0.20% (w/v) |
| | $KH_2PO_4$ | 0.10% (w/v) |
| | $CaCO_3$ | 0.10% (w/v) |
| | Ammonium succinate | 0.50% (w/v) |
| | Trace metal sol. [a] | 1 mL/L |
| Medium 3 | Peptone | 5.00% (w/v) |
| | NaCl | 0.50% (w/v) |
| | Trace metal sol. | 1 mL/L |
| Medium 4 | Soluble starch | 2.00% (w/v) |
| | Glucose | 1.00% (w/v) |
| | NZ-amine | 0.50% (w/v) |
| | Yeast extract | 0.50% (w/v) |
| | $CaCO_3$ | 0.10% (w/v) |
| | $CoCl_2 \cdot 6H_2O$ | 0.2 ppm (w/v) |
| Medium 5 | Starch | 2.40% (w/v) |
| | Glucose | 0.10 (w/v) |
| | Peptone | 0.30 (w/v) |
| | Ehrlich bonito extract | 0.30% (w/v) |
| | Yeast extract | 0.50% (w/v) |
| | $CaCO_3$ | 0.40% (w/v) |
| | Trace metal sol. | 5 mL/L |

| | | |
|---|---|---|
| Medium 6 | Starch | 2.00% (w/v) |

| | | |
|---|---|---|
| | Soybean meal | 1.00% (w/v) |
| | NaCl | 0.30% (w/v) |
| | $CaCO_3$ | 0.30% (w/v) |
| Medium 7 | Oatmeal | 2.00% (w/v) |
| Medium 8 | Glycerol | 3.00% (w/v) |
| | Ehrlich bonito extract | 2.00% (w/v) |
| | $CaCO_3$ | 0.20% (w/v) |
| Medium 9 | Maltose | 5.00% (w/v) |
| | Fermipan RED (Dry yeast) | 1.50% (w/v) |
| | Ebios tablets (dried brewer's yeast) | 2.50% (w/v) |
| | KBr | 1.00% (w/v) |
| | $KH_2PO_4$ | 0.05% (w/v) |
| | $CaCO_3$ | 0.05% (w/v) |
| Medium 10 | Glucose | 2.00% (w/v) |
| | L-Asparagine | 0.50% (w/v) |
| | $CaCO_3$ | 0.01% (w/v) |
| | $KH_2PO_4$ | 0.07% (w/v) |
| | Yeast extract | 0.10% (w/v) |
| Medium 11 | Glucose | 0.50% (w/v) |
| | Corn steep liquor powder | 0.50% (w/v) |
| | Oat meal | 1.00% (w/v) |
| | Pharmamedia | 1.00% (w/v) |
| | $KH_2PO_4$ | 0.50% (w/v) |
| | $MgSO_4 \cdot 7H_2O$ | 0.50% (w/v) |
| | Trace metal sol. | 1 mL/L |

| | | |
|---|---|---|
| Medium 12 | Glycerol | 2.00% (w/v) |
| | Soluble starch | 2.00% (w/v) |
| | Nutrient broth | 2.00% (w/v) |
| | Defatted wheat germ | 1.00% (w/v) |
| | $CaCO_3$ | 0.30% (w/v) |
| Medium 13 | Glucose | 2.00% (w/v) |
| | Peptone | 0.50% (w/v) |
| | Fermipan RED (Dry yeast) | 0.30% (w/v) |
| | Ehrlich bonito extract | 0.50% (w/v) |
| | NaCl | 0.50% (w/v) |
| | $CaCO_3$ | 0.30% (w/v) |
| Medium 14 | Soluble starch | 2.00% (w/v) |
| | Glycerol | 0.50% (w/v) |
| | Hi-gy B (defatted wheat germ) | 1.00% (w/v) |
| | Ehrlich bonito extract | 0.30% (w/v) |
| | Fermipan RED (Dry yeast) | 0.30% (w/v) |
| | $CaCO_3$ | 0.30% (w/v) |
| Medium 15 | Soluble starch | 2.00% (w/v) |
| | Yeast extract | 0.30% (w/v) |
| | New Silver (Gelatin) | 0.30% (w/v) |
| Medium 16 | Soluble starch | 1.00% (w/v) |
| | Sucrose | 2.00% (w/v) |
| | Yeast extract | 0.30% (w/v) |
| | New Silver (Gelatin) | 0.30% (w/v) |
| | Trace metal sol. | 0.5 mL/L |
| Medium 17 | Glucose | 1.00% (w/v) |
| | Maltose | 2.00% (w/v) |
| | Yeast ext. | 0.30% (w/v) |
| | New Silver (Gelatin) | 0.30% (w/v) |
| | Trace metal sol. | 0.5 mL/L |

| | | |
|---|---|---|
| Medium 18 | Glycerol | 2.00% (w/v) |
| | Soybean Meal | 2.00% (w/v) |
| | NaCl | 0.30% (w/v) |
| Medium 19 | Difco PDB | 2.40% (w/v) |
| Medium 20 | Glycerol | 2.00% (w/v) |
| | Molasses | 1.00% (w/v) |
| | Casein | 0.50% (w/v) |
| | Polypeptone | 0.50% (w/v) |
| | $CaCO_3$ | 0.10% (w/v) |
| Seed medium | Starch | 2.40% (w/v) |
| | Glucose | 0.10% (w/v) |
| | Peptone | 0.30% (w/v) |
| | Ehrlich bonito extract | 0.30% (w/v) |
| | Yeast extract | 0.50% (w/v) |
| | $CaCO_3$ | 0.40% (w/v) |

All media were prepared by addition the components described above to tap water and autoclaving at 121˚C for 20 min.

[a]Trace metal sol. was containing 0.1 ppm (w/v) each of $FeSO4 \cdot 7H_2O$, $MnCl_2 \cdot 4H_2O$, $ZnSO4 \cdot 7H_2O$, $CuSO_4 \cdot 5H_2O$ and $CoCl_2 \cdot 6H_2O$ in ion-exchanged water.

**Table S5**. Physicochemical properties of lentindoles A (**1**) and B (**2**).

| | Lentindole A (**1**) | Lentindole B (**2**) |
|---|---|---|
| Appearance | Pale-yellow powder | Yellow solid |
| Molecular formula | $C_{24}H_{36}N_4O_5S$ | $C_{24}H_{36}N_4O_5S$ |
| Molecular weight | 492 | 492 |
| HR-ESI-MS *m/z* | | |
| Calcd. | 493.2479 $[M+H]^+$ | 493.2479 $[M+H]^+$ |
| Found | 493.2468 $[M+H]^+$ | 493.2484 $[M+H]^+$ |
| $[\alpha]_D^{22}$ (*c* 0.1, MeOH) | -14° | 62° |
| UV $\lambda^{MeOH}$ nm (ε) | 240 (8572), 294 (2709) | 240 (8572), 294 (1970) |
| IR ν $cm^{-1}$ | 3296, 2958, 1641,1535, 1467, 1312, 1026, 936, 746 | 3296, 2957, 1635, 1546, 1431, 1312, 1042, 748 |
| Solubility | | |
| Soluble | DMSO, MeOH, $CHCl_3$ | DMSO, MeOH, $CHCl_3$ |

**Table S6.** NMR data of lentindole A (**1**) in DMSO-$d_6$

| Position | Type | $\delta_C$ | $\delta_H$ (mult., $J$ in Hz) |
|---|---|---|---|
| 1 | NH | - | 6.43 (br s) |
| 2 | CH | 93.8 | 4.68 (d,1.2) |
| 3 | C | 75.8 | - |
| 3a | C | 130.9 | - |
| 3b | OH | - | 5.33 (s) |
| 4 | CH | 123.0 | 7.17 (dd, 1.4, 7.4) |
| 5 | CH | 117.5 | 6.65 (ddd, 1.4, 7.4, 7.5) |
| 6 | CH | 129.1 | 7.06 (ddd, 1.4, 7.5, 7.7) |
| 7 | CH | 109.0 | 6.58 (dd, 1.4, 7.7) |
| 7a | C | 150.8 | - |
| 8 | $CH_2$ | 35.9 | 2.38 (m), 1.87 (m) |
| 9 | CH | 43.0 | 3.36 (m) |
| 10 | $CH_2$ | 63.7 | 3.51 (dd, 4.0, 10.9),3.03 (dd, 8.2, 10.9) |
| 11 | NH | - | 7.89 (d, 8.2) |
| 12 | C | 171.8 | - |
| 13 | CH | 48.1 | 4.18 (m) |
| 14 | $CH_3$ | 18.3 | 1.16 (d, 6.9) |
| 15 | NH | - | 7.93 (d, 7.5) |
| 16 | C | 171.0 | - |
| 17 | CH | 51.7 | 4.30 (m) |
| 18 | $CH_2$ | 31.6 | 1.87 (m), 1.75 (m) |
| 19 | $CH_2$ | 29.6 | 2.43 (m) |
| 20 | $CH_3$ | 14.6 | 2.02 (s) |
| 21 | NH | - | 8.00 (d, 8.0) |
| 22 | C | 171.9 | - |
| 23 | $CH_2$ | 44.4 | 2.01 (m) |
| 24 | CH | 25.6 | 2.02 (m) |
| 25 | $CH_3$ | 22.3 | 0.86[a] (dd, 3,4, 6.3) |
| 26 | $CH_3$ | 22.3 | 0.86[a] (dd, 3,4, 6.3) |

[a] Overlapping signals.

**Table S7.** NMR data of lentindole B (**2**) in DMSO-$d_{6f}$

| Position | Type | Conformer 1 (**2-1**) | | Conformer 2 (**2-2**) | |
|---|---|---|---|---|---|
| | | $\delta_C$ | $\delta_H$ (mult., $J$ in Hz) | $\delta_C$ | $\delta_H$ (mult., $J$ in Hz) |
| 1 | NH | - | 6.47 (d, 2.7) | - | 6.77 (d, 4.0) |
| 2 | CH | 82.1 | 5.18 (d, 2.7) | 83.3 | 5.51 (d, 4.0) |
| 3 | C | 88.5 | - | 87.7 | - |
| 3a | C | 132.2 | - | 133.3 | - |
| 3b | OH | - | 5.69 (s) | - | 5.76 (s) |
| 4 | CH | 123.6 | 7.17 (dd, 1.2, 7.5) | 123.8 | 7.19 (dd, 1.0, 7.6) |
| 5 | CH | 117.4 | 6.60 (ddd, 1.1, 7.5, 7.5) | 118.3 | 6.68 (ddd, 1.0, 7.6, 7.5) |
| 6 | CH | 129.2 | 7.00 (ddd, 1.2, 7.5, 7.7) | 129.2 | 7.05 (ddd, 1.0, 7.5, 7.8) |
| 7 | CH | 108.8 | 6.47 (dd, 1.1, 7.7) | 109.9 | 6.54 (dd, 1.0, 7.8) |
| 7a | C | 149.7 | - | 149.1 | - |
| 8 | $CH_2$ | 41.1 | 2.41 (m) | 39.7[a] | 2.22 (dd, 8.5, 12.9)<br>2.38 (dd, 2.6, 12.9) |
| 9 | CH | 59.5 | 3.99 (m) | 58.7 | 4.19 (m) |
| 10 | $CH_2$ | 62.8 | 2.62 (m), 3.29 (m) | 61.1 | 2.51 (m), 3.20 (m) |
| 11 | OH | - | 4.67 (dd, 4.6, 6.7) | - | 4.53 (dd, 5.1, 6.0) |
| 12 | C | 171.7[b] | - | 171.6[b] | - |
| 13 | CH | 46.1 | 4.59 (m) | 45.9 | 4.57 (m) |
| 14 | $CH_3$ | 18.1[c] | 1.22 (d, 6.6) | 17.9[c] | 1.24 (d, 6.8) |
| 15 | NH | - | 8.00 (d, 6.8) | - | 8.20 (d, 7.1) |
| 16 | C | 171.0[d] | - | 170.8[d] | - |
| 17 | CH | 51.3[e] | 4.36 (m) | 51.3[e] | 4.33 (m) |
| 18 | $CH_2$ | 32.0[f] | 1.83[g] (m), 1.74[h] (m) | 31.7[f] | 1.83[g] (m), 1.74[h] (m) |
| 19 | $CH_2$ | 29.8[i] | 2.41 (m) | 29.6[i] | 2.41 (m) |
| 20 | $CH_3$ | 14.7[j] | 2.01 (s) | 14.7[j] | 2.02 (s) |
| 21 | NH | - | 7.98[k] (d, 8.1) | - | 7.95[k] (d, 8.0) |
| 22 | Cf | 171.7[b] | - | 171.6[b] | - |
| 23 | $CH_2$ | 44.5[l] | 1.97[m] (m) | 44.4[l] | 1.97[m] (m) |
| 24 | CH | 25.6[n] | 1.97[m](m) | 25.6[n] | 1.97[m] (m) |
| 25 | $CH_3$ | 22.3[o] | 0.85[p] (d, 3.2) | 22.3[o] | 0.83[p] (d, 3.4) |
| 26 | $CH_3$ | 22.3[o] | 0.85[p] (d, 3.2) | 22.3[o] | 0.83[p] (d, 3.4) |

[a]Overlapping with DMSO-$d_6$ signal and determined by the HSQC experiment.

[b, c, d, f, i, k, l, p]Exchangeable signals.

[e, g, h, j, m, n, o]Overlapping signals.

**Table S8.** Crystallographic data determined by 3D ED/MicroED for lentindole A (**1**)

| Parameter | Value |
|---|---|
| Merged crystals | 3 |
| Resolution | 25.44 - 0.84 (0.87-0.84)[a] |
| Reflections | 22353 (2182) |
| Unique reflections | 2690 (273) |
| Unit cell (Å) | 9.50 (19), 11.68 (16), 24.7 (4) |
| Space group | $P2_12_12_1$ |
| Completeness | 97.0 (97.8) |
| $I$/sig($I$) | 21.87 (1.57) |
| Redundancy | 8.3 (8.0) |
| $R_{int}$ | 23.9 (86.8) |
| $R_{pim}$ | 8.7 (31.5) |
| $R_1$ | 16.31 |
| w$R_2$ | 38.08 |
| Goo$F$ | 1.791 |

[a]Values in parentheses are for the highest resolution shell.

**Table S9.** Deduced function of genes in sBGC of lentindoles.

| Gene | Function (Domain)[a] |
|---|---|
| *ltdA* | NRPS (A-CP) |
| *ltdB* | NRPS (C-A-CP-C) |
| *ltdC* | Carrier protein (CP) |
| *ltdD* | PKS (KS-DH-CP-R) |
| *ltdE* | Hydroxybutyryl-CoA dehydrogenase |
| *ltdF* | Carrier protein (CP) |
| *ltdG* | Acyl-CoA dehydrogenase |
| *ltdH* | Oxoacyl-(acyl carrier protein) synthase III |
| *ltdI* | FkbH-like protein-AMP-binding protein |
| *ltdJ* | Transporter |
| *ltdK* | Hypothetical protein |
| *ltdL* | Luciferase-like monooxygenase |
| *ltdM* | Flavin reductase |
| *ltdN* | Hypothetical protein |

[a]Abbreviation of each NRPS domain were as follow; A: adenylation domain, CP: carrier protein, C: condensation domain, KS: ketosynthase domain, DH: dehydratase domain, R: reductase domain.

**Table S10.** MS-DIAL parameter settings

| Parameter | Value |
| --- | --- |
| Project | |
| MS1 Data type | Profile |
| MS2 Data type | Profile |
| Ion mode | Positive |
| Target | Metabolomics |
| Mode | MSMS |
| Data collection parameters | |
| Retention time begin | 2 min |
| Retention time end | 13 min |
| Mass range begin | 250 Da |
| Mass range end | 2000 Da |
| MS2 mass range begin | 100 Da |
| MS2 mass range end | 1500 Da |
| Centroid parameters | |
| MS1 tolerance | 0.01 |
| MS2 tolerance | 0.01 |
| Isotope recognition | |
| Maximum charged number | 4 |
| Data processing | |
| Number of threads | 16 |
| Peak detection paramters | |
| Smoothing method | Linear Weighted Moving Average |
| Smoothing level | 4 |
| Minimum peak width | 5 |
| Minimum peak height | 3000 |
| Peak spotting parameters | |
| Mass slice width | 0.1 |
| Exclusion mass list (mass tolerance) | (None) |
| Deconvolution parameters | |
| Sigma window value | 0.5 |
| MS2Dec amplitude cut off | 0 |
| Exclude after precursor | TRUE |
| Keep isotope until | 0.5 |
| Keep original precursor isotopes | FALSE |

| | |
|---|---|
| **MSP file and MS/MS identification setting** | |
| MSP file | (In-house spectral library of production media) |
| Retention time tolerance | 0.2 |
| Accurate mass tolerance (MS1) | 0.01 |
| Accurate mass tolerance (MS2) | 0.05 |
| Identification score cut off | 70 |
| Using retention time for scoring | TRUE |
| Using retention time for filtering | TRUE |
| **Text file and post identification(retention time and accurate mass based) setting** | |
| Text file | (No file) |
| Retention time tolerance | 0.1 |
| Accurate mass tolerance | 0.01 |
| Identification score cut off | 85 |
| **Advanced setting for identification** | |
| Relative abundance cut off | 0 |
| Top candidate report | TRUE |
| Adduct ion setting | $[M+H]^+$, $[M+NH_4]^+$, $[M+Na]^+$, $[M+H-H_2O]^+$, $[2M+H]^+$, $[2M+NH_4]^+$, $[2M+Na]^+$, $[M+2H]^{2+}$, $[M+H+NH_4]^{2+}$, $[M+H+Na]^{2+}$, $[M+3H]^{3+}$ |
| **Alignment parameters setting** | |
| Reference file | (Data of MeOH blank sample) |
| Retention time tolerance | 0.2 |
| MS1 tolerance | 0.01 |
| Retention time factor | 0.5 |
| MS1 factor | 0.5 |
| Peak count filter | 0 |
| N% detected in at least one group | 0 |
| Remove feature based on peak height fold-change | FALSE |
| Sample max / blank average | 5 |
| Sample average / blank average | 5 |
| Keep identified and annotated metabolites | TRUE |

| | |
|---|---|
| Keep removable features and assign the tag for checking | TRUE |
| Gap filling by compulsion | TRUE |
| **Tracking of isotope labels** | |
| Tracking of isotopic labels | FALSE |
| **Ion mobility** | |
| Ion mobility data | FALSE |
